# AP2/ERF transcription factor NbERF-IX-33 is involved in the regulation of phytoalexin production for the resistance of *Nicotiana benthamiana* to *Phytophthora infestans*

**DOI:** 10.1101/2021.11.30.468885

**Authors:** Sayaka Imano, Mayuka Fushimi, Maurizio Camagna, Akiko Tsuyama-Koike, Hitoshi Mori, Akira Ashida, Aiko Tanaka, Ikuo Sato, Sotaro Chiba, Kazuhito Kawakita, Makoto Ojika, Daigo Takemoto

## Abstract

Plants recognize molecular patterns unique to a certain group of microbes to induce effective resistance mechanisms. Elicitins are secretory proteins produced by plant pathogenic oomycete genera including *Phytophthora* and *Pythium*. Treatment of INF1 (an elicitin produced by *P. infestans*) induces a series of defense responses in *Nicotiana* species, including reactive oxygen species (ROS) production, transient induction of ethylene production, hypersensitive cell death and accumulation of the sesquiterpenoid phytoalexin capsidiol. In this study, we analyzed the expression profiles of *N. benthamiana* genes after INF1 treatment by RNAseq analysis. Based on their expression patterns, *N. benthamiana* genes were categorized into 20 clusters and 4,761 (8.3%) out of 57,140 genes were assigned to the clusters for INF1-induced genes. All genes encoding enzymes dedicated to capsidiol production, 5-*epi*-aristolochene (EA) synthase (*NbEAS*, 10 copies) and EA dehydrogenase (*NbEAH*, 6 copies), and some genes for ethylene production, such as 1-aminocyclopropane 1-carboxylate (ACC) synthase (*NbACS*) and ACC oxidase (*NbACO*), were significantly upregulated by INF1 treatment. Analysis of *NbEAS1* and *NbEAS4* promoters revealed that AGACGCC (GCC box-like motif) is the essential cis-element required for INF1-induced expression of *NbEAS* genes. Given that the GCC box is known to be targeted by ERF (ethylene-responsive factor) transcription factors, we created a complete list of *N. benthamiana* genes encoding AP2/ERF family transcription factors, and identified 45 out of 337 *AP2/ERF* genes in the clusters for INF1-induced genes. Among INF1-induced *NbERF* genes, silencing of *NbERF-IX-33* compromised resistance against *P. infestans* and INF1-induced production of capsidiol. Recombinant NbERF-IX-33 protein can bind to the promoter sequence of *NbEAS4*, suggesting that NbERF-IX-33 is a transcription factor directly regulating the expression of genes for phytoalexin production.

## INTRODUCTION

Plants recognize a variety of molecules derived from pathogens, including components of microbial cell walls, membranes and secreted proteins (Ranf, 2017; Monjil et al., 2021). Elicitins are small secretory proteins produced by plant pathogenic oomycete genera such as *Phytophthora, Pythium* and *Hyaloperonospora* (Tyler, 2002; Takemoto et al., 2005; Cabral et al., 2011). Treatment with elicitins induces typical defense responses such as hypersensitive cell death, expression of *PR* genes and production of phytoalexins (Milat et al., 1991; Bonnet et al., 1996; Matsukawa et al., 2013). Reports of plant species and cultivars distinctly responsive to elicitins include *Nicotiana* spp., some cultivars of *Brassica rapa* and *Raphanus sativus*, and *Solanum microdontum* (Bonnet et al., 1996; Takemoto et al., 2005; Du et al., 2015). Elicitins from different *Phytophthora* species elicit defense responses in these plants responsive to elicitin, indicating that responsive plants recognize elicitins as a molecular pattern of oomycete pathogens. The most virulent isolates of *Phytophthora parasitica* (syn. *P. nicotianae*) have lost the production of canonical elicitins, or downregulate the expression of the elicitin gene during infection (Ricci et al., 1992; Colas et al., 2001), indicating that the downregulation of elicitin represents a strategy for the pathogen to avoid recognition by the host plant. Elicitins are considered as an essential factor for the reproduction of *Phytophthora* and *Pythium* species (Blein et al., 2002). Elicitins contain a hydrophobic pocket similar to a lipid transfer protein and can bind to phytosterols to take up sterols from plant plasma membranes (Boissy et al., 1996; Mikes et al., 1998). Procuring sterols from the plant plasma membrane may be the primary function of elicitins, which could explain why elicitins are essential for reproduction, as *Phytophthora* and *Pythium* species are unable to produce sterols themselves (Hendrix, 1970).

Potato late blight caused by *Phytophthora infestans* is one of the most devastating and economically important plant diseases. *P. infestans* was the causal agent of the Irish potato famine in the 1840s, and even in present times, worldwide yield losses caused by this pathogen exceed $6 billion per year (Haverkort et al., 2008). *P. infestans* has at least six genes for elicitins (Jiang et al., 2006), and INF1 is the most abundantly produced elicitin. Although INF1 treatment does not elicit a noticeable defense response in most *Solanum* species, including potato (*S. tuberosum*), a receptor-like protein ELR was isolated from the wild potato *S. microdontum*, which is able to recognize INF1 and induce a defense response (Du et al., 2015). Recognition of INF1 is important for the non-host resistance of *Nicotiana benthamiana* to *P. infestans*. Gene silencing of *inf1* in *P. infestans* enhanced the virulence of the pathogen on *N. benthamiana* (Kamoun et al., 1998), suggesting that recognition of INF1 is essential for the defense induction in *N. benthamiana* against *P. infestans*. However, in contrast to *S. microdontum*, the gene for the recognition of INF1 has not yet been identified in *N. benthamiana*.

*N. benthamiana* is an ideal model host plant to study the molecular mechanisms underlying non-host resistance of Solanaceae plants against *P. infestans*. Previously, we performed virus-induced gene silencing (VIGS)-based screenings to isolate genes essential for the resistance of *N. benthamiana* against *P. infestans* (Matsukawa et al., 2013; Ohtsu et al., 2014; Shibata et al., 2016; Takemoto et al., 2018; Mizuno et al., 2019a). So far, thirty-three genes have been identified from approx. 3,000 randomly gene-silenced plants as the essential genes for full resistance of *N. benthamiana* against *P. infestans*. Besides random gene silencing, *N. benthamiana* can also readily be used for targeted gene-silencing to further examine the involvement of homologs of known defense-related genes in the resistance to *P. infestans* (Shibata et al., 2010, 2011, 2016).

A major group of genes isolated from the VIGS screening are genes coding for enzymes in the mevalonate (MVA) pathway, for the production of sterols and a wide variety of isoprenoid-derived secondary metabolites (Shibata et al., 2016). In addition, silencing of genes for *NbEAS* (*5-epi-aristolochene synthase*) and *NbEAH* (*5-epi-aristolochene dihydroxylase*), encoding the enzymes dedicated to the production of the sesquiterpenoid phytoalexin capsidiol, compromised resistance of *N. benthamiana* to *P. infestans*. Six genes related to ethylene biosynthesis were also isolated as essential genes for the resistance of *N. benthamiana* against *P. infestans*. Expression of *NbEAS* and *NbEAH* genes was induced by the treatment with ethylene, and suppressed by gene silencing of *NbEIN2*, a central regulator of ethylene signaling, indicating that phytoalexin production is regulated by ethylene in *N. benthamiana* (Shibata et al., 2010, 2016; Ohtsu et al., 2014). However, the detailed mechanism of how ethylene signaling regulates the genes for phytoalexin production in *N. benthamiana* has not been fully elucidated.

In a previous study, we performed RNAseq analysis of *N. benthamiana* treated with INF1 at a single time point (24 h after treatment, Rin et al., 2020). While these findings were insightful, they merely represent a single snapshot and are therefore insufficient to capture the dynamic expression processes during pathogen defense. In the present study, we performed RNAseq analysis of INF1 elicited *N. benthamiana* for various time points to obtain more detailed expression profiles of the genes involved in defense response. The activity of *NbEAS* gene promoters was analyzed to identify the cis-acting element essential for the INF1-induced expression of *NbEAS* genes. Moreover, we created a complete catalog of the AP2/ERF (APETALA 2/ethylene-responsive element binding factor) transcription factor genes for *N. benthamiana* and analyzed their expression profiles to identify the transcription factor directly regulating the expression of *NbEAS* genes.

## MATERIALS AND METHODS

### Biological Materials, Growth Conditions, Inoculation and Treatment

*N. benthamiana* line SNPB-A5 (Shibata et al., 2016) was grown in a growth room at 23°C with 16 h of light per day. *P. infestans* isolate 08YD1 (Shibata et al., 2010) was maintained on rye-media at 20°C in the dark. Preparation of zoospore suspension of *P. infestans* and inoculation of *N. benthamiana* leaves (approx. 45 days old) with a zoospore suspension of *P. infestans* was performed as described previously (Shibata et al., 2010). INF1 elicitor was prepared from *Escherichia coli* (strain DH5α) carrying an expression vector for INF1, pFB53, as previously reported (Kamoun et al., 1997; Shibata et al., 2010). *N. benthamiana* leaves were treated with 150 nM INF1 solution as previously described (Shibata et al., 2010).

### Measurement of Reactive Oxygen Species (ROS) Production

The relative intensity of ROS generation was determined by counting photons from L-012-mediated chemiluminescence. For the detection of ROS production at early time points (for Figure 1A), *N. benthamiana* leaf discs (4 mm in diameter) were excised with a cork borer and floated on 100 μl distilled water in a 96-well microplate (Nunc 96F microwell white polystyrene plate, Thermo Fisher Scientific, Waltham, MA, USA) overnight in a growth chamber (23°C). Just before the measurement, water in each well was replaced with 50 μl of water or 150 nM INF1 containing 1 mM L-012 (Wako Pure Chemical, Osaka, Japan). Chemiluminescence was measured by a multiplate reader (TriStar LB941; Berthold Technologies, Bad Wildbad, Germany). For the detection of ROS production at relatively later time points (for Figure 1B), *N. benthamiana* leaves were infiltrated with water or 150 nM INF1 and incubated in a growth room at 23°C, and 0.5 mM L-012 was allowed to infiltrate to the intercellular space of leaves before the measurement. ROS production was measured as chemiluminescence using Lumino Graph II EM (ATTO, Tokyo, Japan).

**FIGURE 1.**
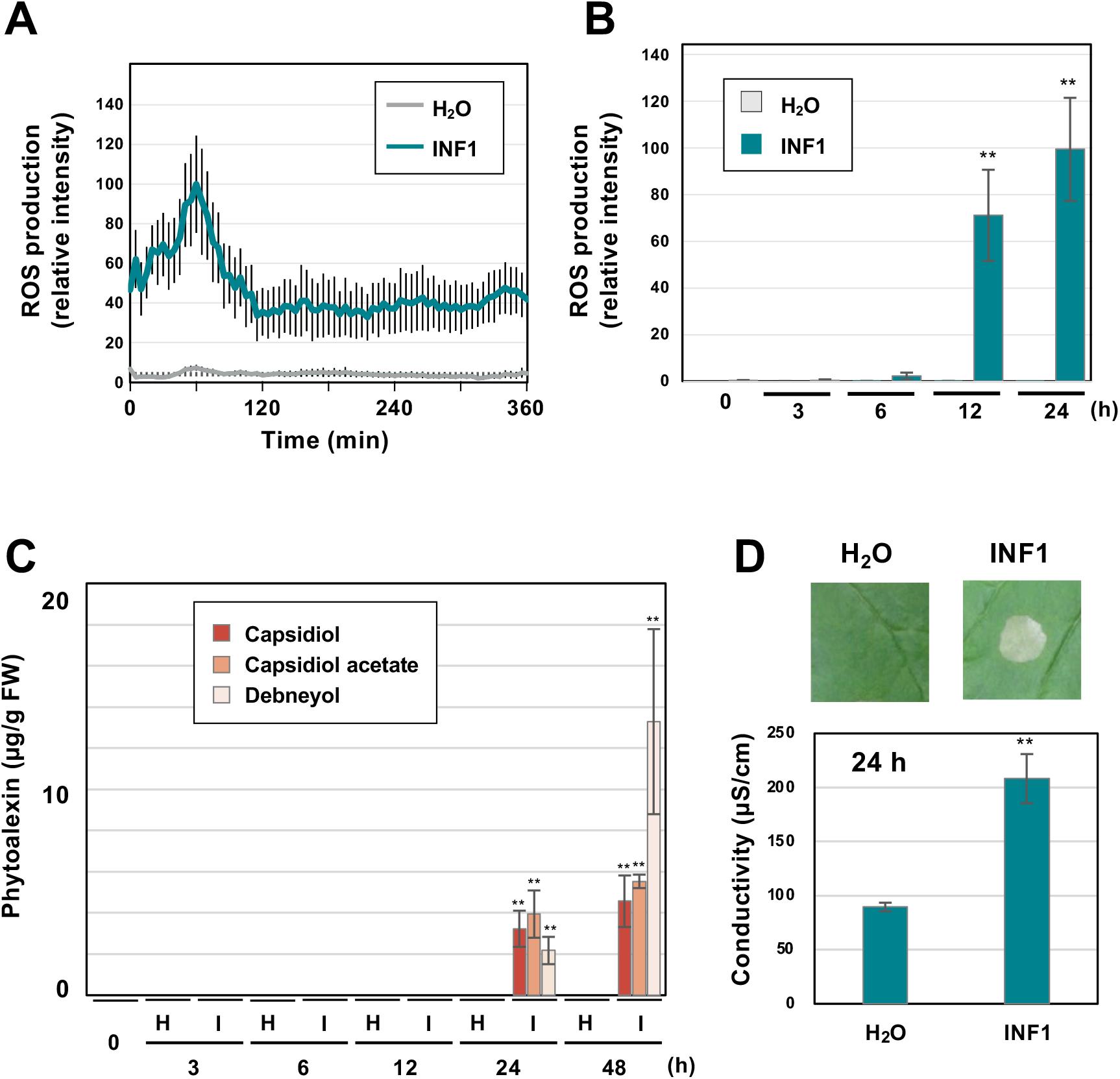
Induction of defense responses of *Nicotiana benthamiana* leaves treated with INF1, a secretory protein derived from *Phytophthora infestans*. **(A)** INF1-induced reactive oxygen species (ROS) production in *N. benthamiana*. Leaf discs were placed in water (H_2_O) or 150 nM INF1 solution containing L-012, and L-012 mediated chemiluminescence was measured. Data are means ± standard error (SE) (n = 8). **(B)** Leaves of *N. benthamiana* were treated with water, or 150 nM INF1 by infiltration and ROS production was detected at indicated time after the treatment. Data are means ± standard error (SE) (n = 6). **(C)** Phytoalexins were extracted at the indicated time after water (H) or 150 nM INF1 (I) treatment and quantified by LC/MS. Data are means ± SE (n = 3). **(D)** Leaves of *N. benthamiana* were treated with water or 150 nM INF1, and electrolyte leakage was quantified at 24 h after INF1 treatment. Data are means ± SE (n = 6). Photographs were taken 72 h after the treatment. Data marked with asterisks are significantly different from control as assessed by the two-tailed Student’s *t*-test: \*\**P* < 0.01.

### Ion Leakage Assay

The severity of cell death induced by INF1 treatment was quantified by Ion leakage assay as previously described (Mizuno et al., 2019b). Four leaf disks (5 mm diameter) of *N. benthamiana* treated with water or 150 nM INF1 were floated on 1 ml water under light condition for 6 h. Ion conductivity was measured by LAQUAtwin compact conductivity meter (Horiba, Kyoto, Japan).

### Quantitative Analysis of Phytoalexin Production by Liquid Chromatography (LC)

For the quantification of phytoalexins shown in Figure 1C, leaves of *N. benthamiana* (50 mg, fresh weight) treated with elicitor were soaked in 1.5 ml ethyl acetate with gentle shaking for an hour. After the evaporation of ethyl acetate, the dried substances were redissolved in 100 μl of 1:1 acetonitrile: water (v/v) and produced phytoalexins were measured by LC/MS (Accurate-Mass Q-TOF LC/MS 6520, Agilent Technologies, Santa Clara, CA, USA) with ODS column Cadenza CD-C18, 75 x 2 mm (Imtakt, Kyoto, Japan). For the quantification of phytoalexins shown in Figure 8C, extraction and quantification of capsidiol were performed as previously described (Matsukawa et al., 2013).

### RNA-seq Analysis and Clustering of *N. benthamiana* Genes

Total RNA was extracted from *N. benthamiana* leaves using the RNeasy Plant Mini Kit (QIAGEN, Hilden, Germany). Libraries were constructed using KAPA mRNA Capture Kit (Roche Diagnostics, Tokyo, Japan) and MGIEasy RNA Directional Library Prep Set (MGI, Shenzhen, China), and sequenced on DNBSEQ-G400RS (MGI) with 150 bp paired-end protocol. The RNA-seq reads were filtered using trim-galore v.0.6.6 (Martin, 2011, bioinformatics.babraham.ac.uk) and mapped to the *N. benthamiana* genome (ver. 1.0.1, https://solgenomics.net, Bombarely et al., 2012) using HISAT2 v.2.2.1 (Kim et al., 2019) and assembled via StringTie v.2.1.7 (Kovaka et al., 2019). Significant differential expression was determined using DESeq2 v.1.32.0 (Love et al., 2014). All software used during RNA-seq analysis was run with default settings. For the clustering of *N. benthamiana* genes, K-means clustering was performed using TimeSeriesKMeans from the tslearn (v.0.5.2) Python package (k=20). Before clustering, the log2-fold expressions for each gene were pre-processed, so that the mean expression of each time series equaled 0. RNA-seq data reported in this work are available in GenBank under the accession number DRA013037.

### Quantitative Analysis of Ethylene Production by Gas Chromatography (GC)

Ethylene production was quantified as previously described (Matsukawa et al., 2013). Leaf disks (10 mm in diameter) of *N. benthamiana* were excised with a cork borer and placed in a 5-ml GC vial sealed with a rubber plunger. After 3 h of incubation, a 1-ml air sample in the vial was collected using a glass syringe. The quantity of ethylene in the collected air was measured using a gas chromatograph equipped with a flame thermionic detector (GC-353, GL Sciences, Tokyo, Japan) and CP-Permabond Q column (Varian Inc., Middelburg, Netherlands). GC was performed with column temperature at 50°C, injection temperature at 120°C, and detector temperature at 150°C using N_2_ as the carrier gas.

### Construction of Vectors and Transformation of *Agrobacterium tumefaciens*

Base vectors, primer sets and templates for the PCR amplification of DNA fragments for vector construction are listed in Supplementary Table 1. Sequences of primers used for the construction of vectors are listed in Supplementary Table 2. Gene fragments were amplified using PrimeStar HS DNA polymerase (Takara Bio, Kusatsu, Japan) and cloned into the vector using In-Fusion HD Cloning Kit (Takara Bio). Gene fragments in pTV00 vectors for VIGS induction were assessed using the SGN VIGS tool (Fernandez-Pozo et al., 2015) to exclude unexpected off-target effects. Vectors for transient expression or VIGS were transformed into *A. tumefaciens* (strain C58C1) by electroporation with a MicroPulser electroporator (Bio-Rad, Hercules, CA, USA). *A. tumefaciens* transformants were selected on LB media containing 50 μg/ml rifampicin, 50 μg/ml kanamycin, and 50 μg/ml ampicillin at 28 °C.

### Transient Gene Expression by Agroinfiltration

*A. tumefaciens* carrying expression vectors (pNPP40 and pNPP243 derivatives, Supplementary Table S1) were cultured to saturation in LB medium at 28 °C and bacterial cells were collected by centrifugation at 16,000 x g for 1 min. The bacterial cells were then resuspended in MMA infiltration solution to a final OD_600_ of 0.5 (MMA: 5 mg/ml MS salts, 1.95 mg/ml MES, 20 mg/ml sucrose, 200 μM acetosyringone, pH = 5.6) and incubated at 28°C for 2 h. The suspensions were allowed to infiltrate the intercellular space of *N. benthamiana* leaves using a syringe without a needle.

### Quantitative RT-PCR

Total RNAs were isolated from *N. benthamiana* leaves using TRIzol Reagent (Thermo Fisher Scientific) and cDNA synthesis was conducted using ReverTra Ace-α- (Toyobo, Osaka, Japan). Quantitative RT-PCR (qRT-PCR) analysis was performed using LightCycler Quick System 350S (Roche Applied Science, Penzberg, Germany) with Thunderbird SYBR qPCR Mix (Toyobo). The expression of *N. benthamiana EF-1α* gene was used as an internal standard. Gene-specific primers used for expression analysis were listed in Supplementary Table S2.

### Fluorescence Microscopy

To visualize the activation of *NbEAS4* promoter by GFP marker, fluorescence images were recorded using BX51 fluorescence microscope (Olympus, Tokyo, Japan) equipped with color CMOS camera Wraycam-NF500 (Wraymer, Osaka, Japan).

### Virus-induced Gene Silencing (VIGS)

The induction of VIGS was carried out as previously reported (Ratcliff et al., 2001; Shibata et al., 2010). *A. tumefaciens* strain GV3101 delivering the binary TRV RNA1 construct pBINTRA6, and the TRV RNA2 vector pTV00 or its derivatives (Supplementary Table 1), were cultured to saturation in LB media. Bacterial cells were collected by centrifugation at 16,000 x *g* for 1 min. The bacterial cells were then resuspended in 10 mM MES-NaOH (pH 5.6), 10 mM MgCl_2_ and 150 μm acetosyringone (final OD_600_ = 0.5) and incubated at room temperature for 2 h. The cultures were mixed in a 1:1 ratio (RNA1/RNA2), to infiltrate leaves of *N. benthamiana* using a syringe without a needle. After 3-4 weeks of infiltration, the upper leaves of the inoculated plants were used for experiments.

### Expression of Recombinant Proteins in *E. coli*

The MBP-NbERF-IX-33a fusion protein was prepared using the pMAL Protein Fusion and Purification System (New England Biolabs). *E. coli* cells (DH5) carrying pMAL-c5x (New England Biolabs) or pMAL-NbERF-IX-33a were cultured overnight at 37°C in rich medium (1% tryptone, 0.5% yeast extract, 0.5% NaCl, 0.2% glucose) containing 100 μg/ml ampicillin and grown until the optical density (OD_600_=0.6). Production of the protein was induced by adding 0.3 mM IPTG for 5 h. The culture (20 ml) was centrifuged to collect *E. coli* cells. The cells were then resuspended in 6 ml of column buffer [20 mM Tris-HCl (pH 7.5), 200 mM NaCl, 1 mM EDTA] and frozen at −20°C. Frozen cells were thawed on ice and then homogenized using a sonicator (Sonicator cell disruptor Model W-225R, Heat Systems-Ultrasonics Inc., Plainview, NY, USA). The crude cell extracts were centrifuged at 10,000 x g, 4°C for 5 min, and the supernatant was adsorbed to a 0.4 ml amylose resin (New England Biolabs) at 4°C. The column was washed with 4 ml of column buffer, followed by elution of the fusion protein with 1 ml of column buffer containing 10 mM maltose. The fractions of purified proteins were assessed by SDS-PAGE and eluted fractions were dialyzed against water with the dialysis tubing (SnakeSkin dialysis tubing, 3.5 kD molecular mass cutoff, Thermo Fisher Scientific) overnight at 4°C.

### Electrophoresis Mobility Shift Assay (EMSA)

Electrophoresis mobility shift assay Kit (Thermo Fisher Scientific) was used to detect binding of NbERF-IX-33a to the *NbEAS4* promoter. The DNA fragment of *NbEAS4* promoter (30 ng) and MBP or MBP-NbERF-IX-33a proteins (1 μg) were mixed and incubated in 10 μl binding buffer [125 mM HEPES-KOH, 20 mM KCl, 0.5 mM EDTA, 50% (v/v) glycerol, 5 mM DTT, pH7.5] for 20 min at 4°C. 2 μl of 6 x loading buffer was added to stop the reaction. The complexes were resolved on 6% non-denaturing polyacrylamide gels (Thermo Fisher Scientific) and stained with SYBR green dye.

## RESULTS

### Induction of ROS Production, Phytoalexin Accumulation and Cell Death in *N. benthamiana* treated with INF1

Prior to the RNAseq analysis of *N. benthamiana* genes induced by INF1 treatment, typical defense responses induced by INF1 treatment were observed under our experimental conditions. Induction of reactive oxygen species (ROS) production is one of the common phenomena observed during the induction of plant disease resistance. Using small leaf disks of *N. benthamiana*, transient ROS production was detected within 60 min after INF1 treatment (Figure 1A). Significant increase in ROS production was also observed at 12 h and 24 h after treatment (Figure 1B). These results are consistent with previous studies that reported biphasic ROS production in plants during the plant defense (Chai and Doke, 1987; Levine et al., 1994; Yuan et al., 2021). The production of sesquiterpenoid phytoalexins, capsidiol, debneyol and capsidiol 3-acetate were significantly increased at 24 h and 48 h after treatment with 150 nM INF1, while the production of phytoalexins was below the detection limit at earlier times (Figure 1C). Induction of visible cell death became obvious at around 48 h after INF1 treatment, while the increase of ion leakage from the INF1-treated plant tissue (indicating the initiation of cell death) was already detectable at 24 h (Figure 1D).

### Clustering of *N. benthamiana* Genes Based on Their Expression Patterns after INF1 Treatment

To investigate the expression profile of all *N. benthamiana* genes during the induction of defense responses by INF1 treatment, RNA-seq analysis was performed for *N. benthamiana* leaves treated with water or INF1 at 0, 3, 6, 12 and 24 h after the treatment. Based on their time dependent expression patterns, all genes were categorized into 20 groups using K-means clustering. Among the 57,140 *N. benthamiana* annotated genes, 14,390 genes (25.2%) were not assigned to any cluster because no expression could be determined at any point in time. Among the 20 clusters, clusters 2, 4, 10, 14 and 16 were the clusters that contained significantly more INF1-inducible genes, and these include 4,761 (8.3%) out of 57,140 genes (Figure 2). Especially, the genes in cluster 14 (contains 336 genes) were significantly up-regulated after INF1 treatment, with more than 300-fold induction at 24 h (average in cluster 14) compared to control (Figure 2). The lists of genes in clusters for INF1-induced *N. benthamiana* genes are shown in Supplementary Tables 3-7.

**FIGURE 2.**
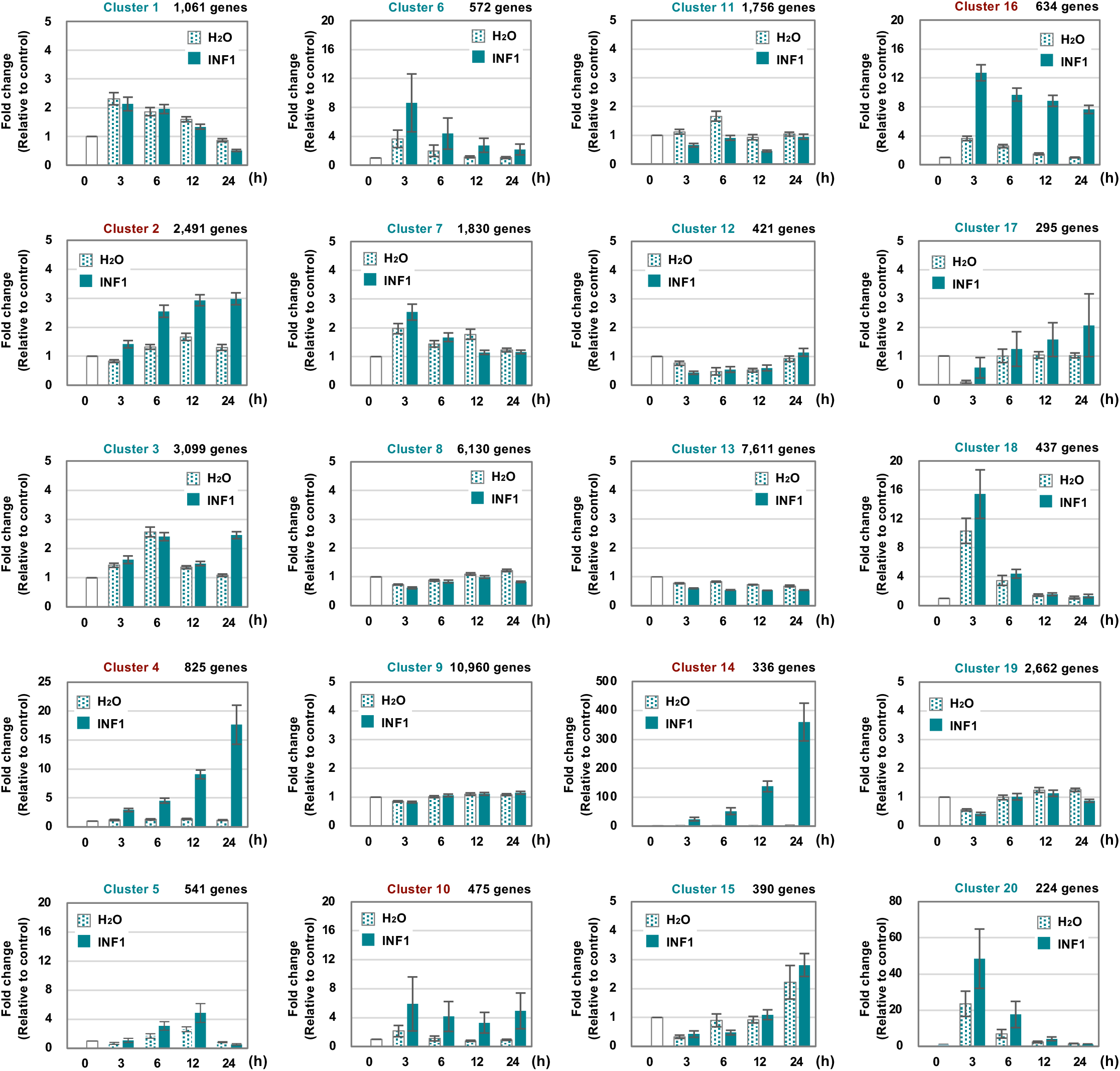
Clustering of 57,140 *Nicotiana benthamiana* genes based on their expression profiles in leaves treated with water or 150 nM INF1. 14,390 (25.2%) of genes were not assigned to any cluster because no or significantly lower expression was detected. See method for detailed procedure of the gene clustering. Graphs shown are average expression profiles of 100 genes (selected based on their high average FPKM value) from each cluster. Data are means ±SE (n = 100). Clusters for INF1 induced genes are shown in red letters.

Cluster 14 includes all 10 copies of *NbEAS* and 5 out of 6 *NbEAH* genes which are genes specific for the capsidiol production (Figure 3, Supplementary Table 8). Only negligible expression of genes in cluster 14 was observed in control and water-treated leaves, indicating that genes specifically involved in the production of phytoalexin are under strict regulation.

**FIGURE 3.**
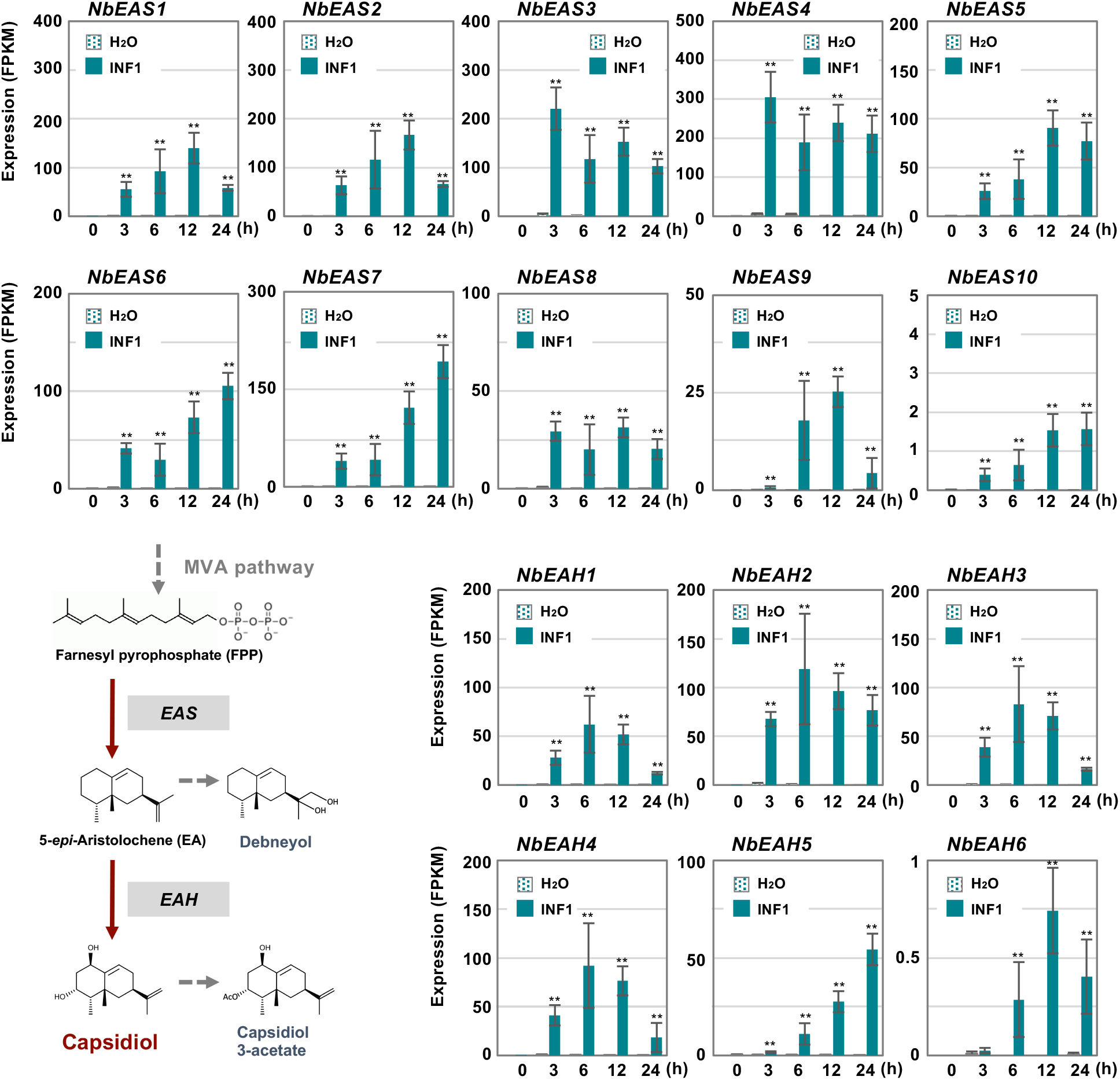
The time course of expression of *Nicotiana benthamiana* genes encoding dedicated enzymes for phytoalexin production. The gene expression (FPKM value) was determined by RNA-seq analysis of *N. benthamiana* leaves treated with water or 150 nM INF1 for 0 h, 3 h, 6 h, 12 h and 24 h. Data are mean ± SE (*n* = 3). Asterisks indicate a significant difference from the control (water-treated) as assessed by two-tailed Student’s *t*-test, \*\**P* < 0.01. *EAS* 5-*epi*-aristolochene synthase, *EAH* 5-*epi*-aristolochene-1,3-dihydroxylase

In contrast, most genes for the enzymes in the MVA pathway (required for the production of capsidiol precursor, farnesylpyrophosphate, FPP) were categorized into other clusters as most of them have a basal constitutive expression in control and water treated leaves, probably reflecting that FPP is a common precursor of isoprenoids and phytosterol (Supplementary Table 9, Figure 4). Among five *acetyl-CoA thiolase* (*ACAT*) genes, only the expression of *NbACAT1b* was highly upregulated from an early time point. Out of five *3-hydroxy-3-methylglutaryl-CoA* (HMG-CoA) synthase genes, *NbHMGS1a* and *1b* were found to be induced by INF1 treatment. Similarly, among five *HMG-CoA reductase* (*NbHMGR*) genes, *NbHMGR2* and *1a* were significantly upregulated by INF1 treatment. In the case of genes for mevalonate-5-kinase (*NbMVK*, 2 copies), phosphomevalonate kinase (*NbPMVK*, 2 copies), and mevalonate-5-pyrophosphate decarboxylase (*NbMVD*, 2 copies), they were all upregulated by INF1 treatment. Likewise, two out of four genes encoding isopentenyl pyrophosphate (IPP) isomerase, *NbIPPI1b* and *2a*, were appeared to mainly contribute to the activation of the conversion of IPP to dimethylallyl pyrophosphate (DMAPP) during the induction of capsidiol production, while stable and constant expression was observed for *NbIPPI2b*. Finally, among 4 genes for FPP synthase (*NbFPPS*), *NbFPPS1a, NbFPPS1b, NbFPPS2b* were significantly upregulated (Figure 4).

**FIGURE 4.**
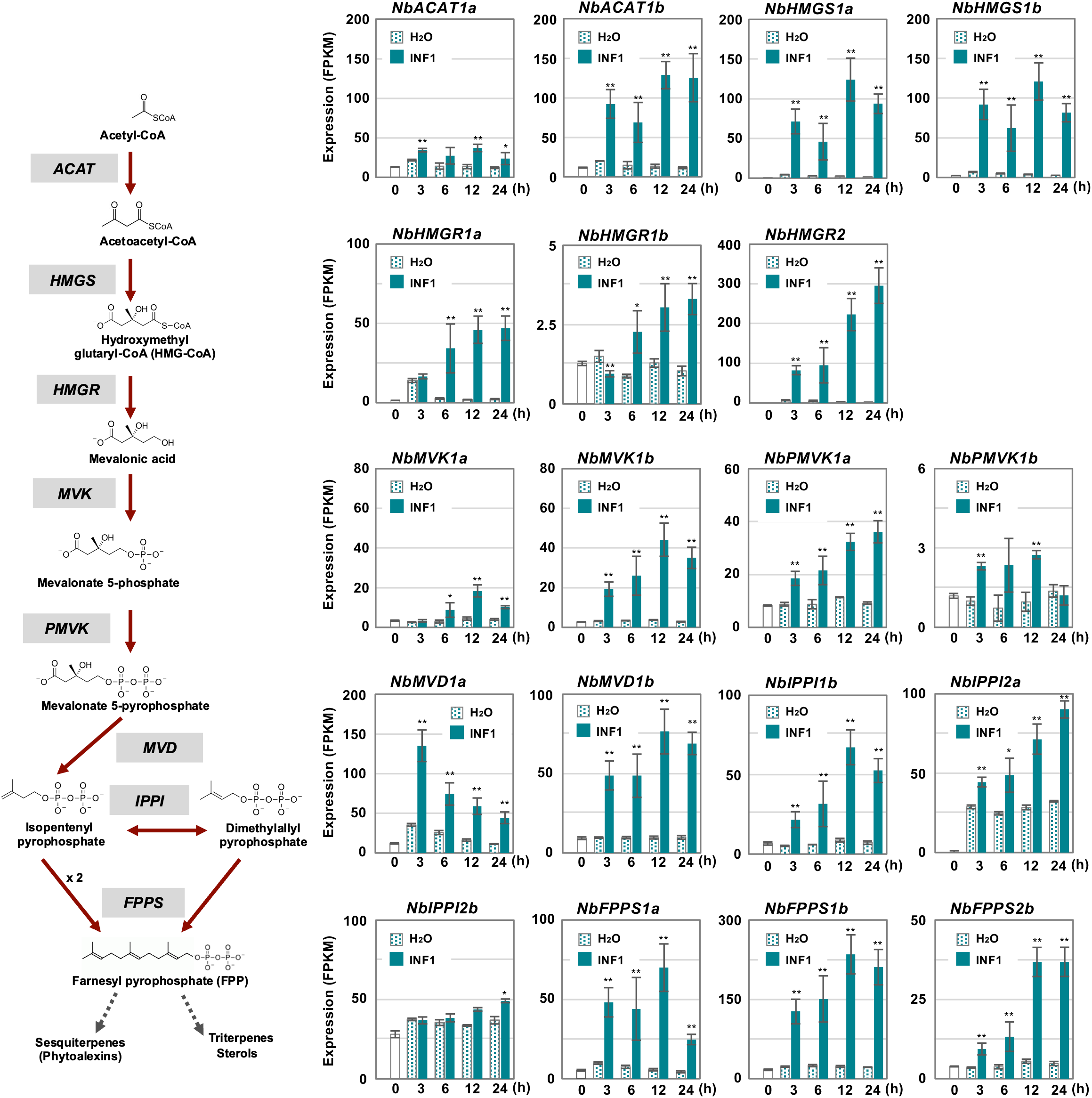
The time course of expression of *Nicotiana benthamiana* genes encoding enzymes in the mevalonate pathway. Gene expression (FPKM value) was determined by RNA-seq analysis of *N. benthamiana* leaves treated with water (H_2_O) or 150 nM INF1 for 3 h, 6 h, 12 h and 24 h. Data are means ±SE (n = 3). Data marked with asterisks are significantly different from control as assessed by the two-tailed Student’s *t*-test: \*\**P* < 0.01 \**P* < 0.05. *ACAT*, Acetoacetyl-CoA thiolase; *HMGS*, Hydroxymethylglutaryl-CoA synthase; *HMGR*, Hydroxymethylglutaryl-CoA-reductase; *MVK*, Mevalonate-5-kinase; *PMVK*, Phosphomevalonate kinase; *MVD*, Mevalonate-5-pyrophosphate decarboxylase; *IPPI*, Isopentenyl pyrophosphate isomerase; *FPPS*, Farnesyl pyrophosphate synthase.

### Identification of Cis-element for INF1-induced Expression of *NbEAS* Genes

To further investigate the strict regulation of *NbEAS* genes, we analyzed promoters of *NbEAS1* and *NbEAS4* in this study. To evaluate the promoter activity of these genes, plasmid vectors containing the *GFP* gene under the control of *NbEAS1* or *NbEAS4* promoter (P_*NbEAS1:GFP* and P_*NbEAS4:GFP*) were transiently transformed into *N. benthamiana* leaves by Agroinfiltration. Basal expression of GFP was detected after agroinfiltration using *A. tumefaciens* containing these promoter:GFP vectors under fluorescence microscopy, presumably because plant defense was weakly induced by the infection with *A. tumefaciens*. Nonetheless, the relative expression levels of GFP were significantly enhanced by treatment with INF1. When the promoter length exceeded 260 bp upstream from the start codon (−260) was used, *GFP* expression was induced by INF1 treatment for both *NbEAS1* and *NbEAS4* promoters. When the *NbEAS1* promoter was shortened to a length of 230 bp, INF1-induced *GFP* expression was no longer observed, whereas the *NbEAS4* promoter remained functional. INF1-induced *GFP* expression ceased however when the *NbEAS4* promoter was further trimmed to a length of 200 bp (Figure 5A). From these findings it follows that the regions essential for the induction of *NbEAS1* and *NbEAS4* expression by INF1 are located between −260 and −230 of the *NbEAS1* promoter, and between −230 and −200 of the *NbEAS4* promoter. In the corresponding region of *NbEAS1* and *NbEAS4* promoter, a sequence similar to the ethylene-responsive element GCC box (AGCCGCC), AGACGCC, was identified (Figure 5B). To confirm the importance of the GCC box-like motif on the promoter activity, we introduced mutations to the motif in the P_*NbEAS4:GFP* vector, which led to substitutions of G to T (A<u>T</u>AC<u>T</u>CC). We designated the vector carrying the mutated GCC box-like motif as P_*NbEAS4* (TT). G to T mutations in the GCC box-like motif significantly reduced both the fluorescence of GFP in *N. benthamiana* leaves 48 h after Agroinfiltration, as well as the relative expression level of *GFP* (Figure 5C), indicating that the GCC-box like motif is the cis-acting element essential for the INF1-induced expression of *EAS4*. GCC-box like motifs can be found in most of *NbEAS* and *NbEAH* genes (Supplementary Figure 1), while the GCC-box was not found in the promoter regions of genes in the MVA pathway, indicating that production of the capsidiol precursor FPP and capsidiol production are probably controlled via distinctive regulatory mechanisms.

**FIGURE 5.**
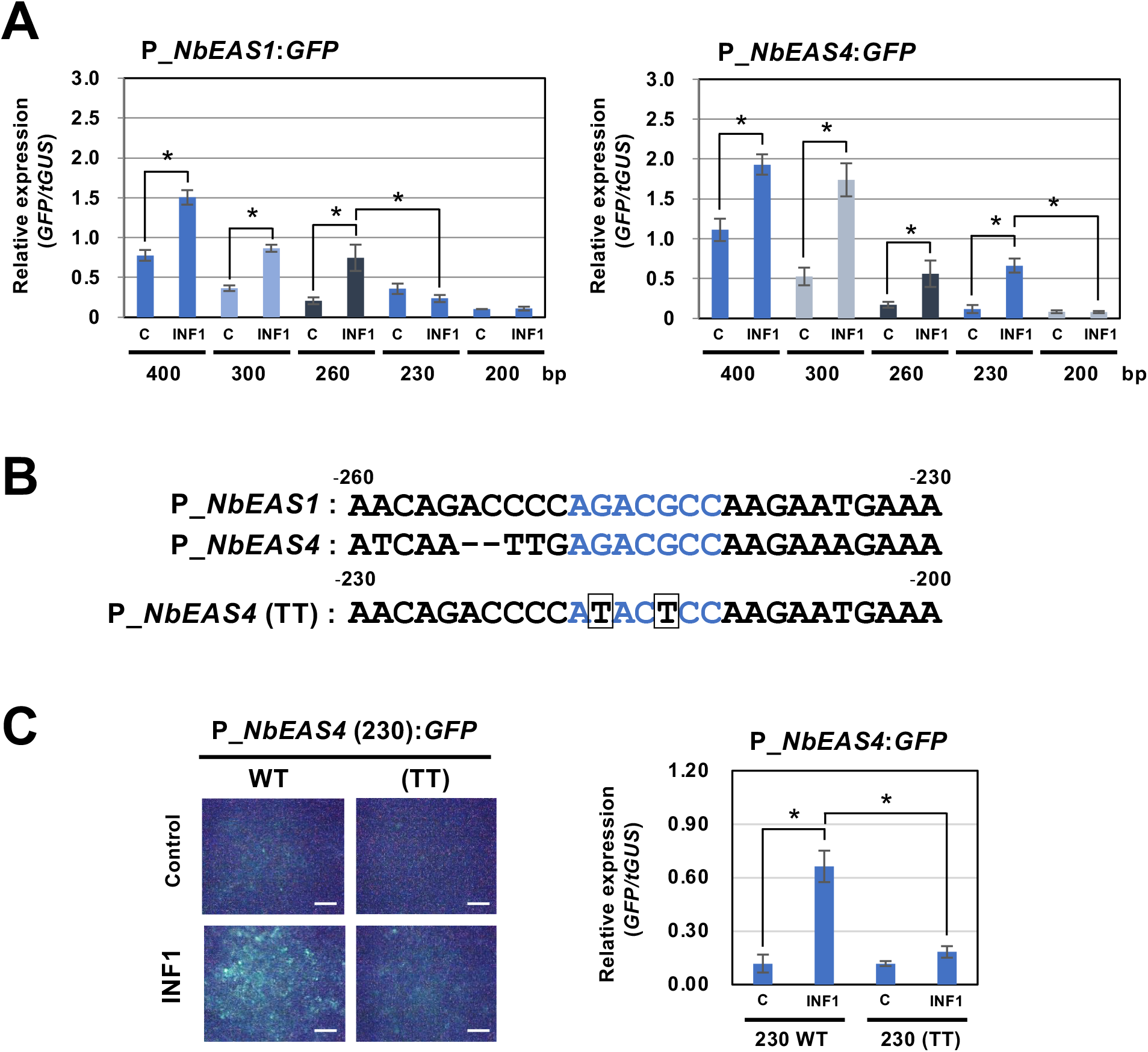
Identification of cis-element for INF1-induced expression of *NbEAS* genes. **(A)** Expression of *GFP* gene under the control of the indicated length of *NbEAS1* and *NbEAS4* promoter (P_*EAS*). Total RNA was isolated from *N. benthamiana* leaves 48 h after co-inoculation with *A. tumefaciens* strains containing expression vectors for P_*EAS*:*GFP*, INF1 or control (C) vector, and truncated *GUS (tGUS*) gene under the control of 35S promoter as internal standard. Expression of the *GFP* gene was assessed by qRT-PCR with *GFP* primers and values were normalized to the expression of *tGUS*. Means ±SE (n = 3). Data marked with asterisks are significantly different as assessed by the two-tailed Student’s *t*-test: \**P* < 0.05. **(B)** Alignment of −260 to −230 of *NbEAS1* promoter and −230 to −200 of *NbEAS4* promoter. GCC box-like motifs were shown in red letters. Mutations introduced in *NbEAS4* (TT) were shown below the alignment by boxes. **(C)** Expression of *GFP* gene under the control of 230 bp of wild type or mutated (TT) *NbEAS4* promoter. (Left) Fluorescence micrographs of *N. benthamiana* leave 48 h after *A. tumefaciens* inoculation containing indicated expression vectors. Bars = 500 μm. (Right) Total RNA was isolated from *N. benthamiana* leaves 48 h after co-inoculation with *A. tumefaciens* strains containing expression vectors. Expression of the *GFP* gene was assessed by qRT-PCR with *GFP* primers and values were normalized to the expression of *tGUS*. Means ± SE (n = 3). Data marked with asterisks are significantly different as assessed by the two-tailed Student’s *t*-test: \**P* < 0.05.

### Expression Profile of *N. benthamiana* Genes for Enzymes Involved in the Ethylene Production

Previously, we have isolated three genes essential for resistance of *N. benthamiana* to *P. infestans* belonging to the methionine cycle, which is related to the production of the ethylene precursor SAM (S-adenosyl-L-methionine) (Shibata et al., 2016, Figure 6). In this study, we created a list of genes for putative enzymes in the methionine cycle and for ethylene biosynthesis (Supplementary Table 10) to investigate their expression profiles after INF1 treatment. Given that *N. benthamiana* has an allopolyploid genome (Goodin et al., 2008), two highly homologous genes were frequently found, and which were predicted to be derived from the two ancestral *Nicotiana* species of *N. benthamiana*. Such highly homologous pairs of genes were named by appending either a or b to the gene name (Matsukawa et al., 2013; Shibata et al., 2016; Rin et al., 2020). For example, the highly homologous genes for cystathionine γ-synthase (CGS) were designated as *NbCGS1a* and *NbCGS1b*.

**FIGURE 6.**
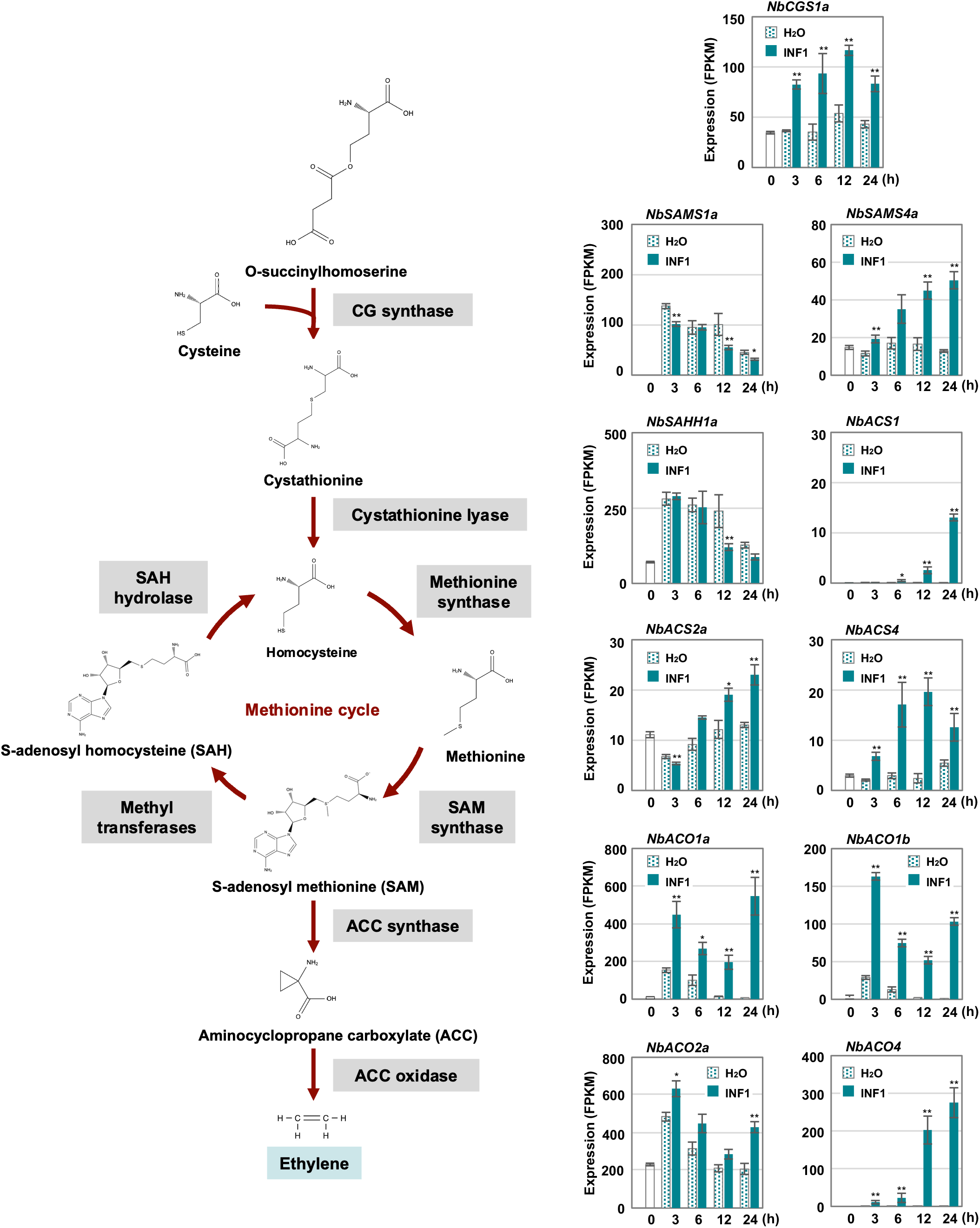
Expression profiles of *Nicotiana benthamiana* genes encoding enzymes for ethylene production. The gene expression (FPKM value) was determined by RNA-seq analysis of *N. benthamiana* leaves treated with water and 150 nM INF1 for 0 h, 3 h, 6 h, 12 h, 24 h. Data are means ±SE (n = 3). Data marked with asterisks are significantly different from control as assessed by the two-tailed Student’s *t*-test: \*\**P* < 0.01, \**P* < 0.05. CGS, Cystathionine gamma-synthase; SAMS, S-adenosylmethionine synthetase; SAHH, S-adenosyl homocysteine hydrolase; ACS, Aminocyclopropane carboxylate synthase; ACO, Aminocyclopropane carboxylate oxidase.

Among the 32 putative methionine cycle genes (Supplementary Table 10), only two genes (*NbCGS1a* and *NbSAM4a*) were moderately upregulated by INF1 treatment (Figure 6). For genes encoding enzymes specific for ethylene production, 24 1-aminocyclopropane-1-carboxylate (ACC) synthase (*NbACS*) and 16 ACC oxidase (*NbACO*) genes were identified in the genome of *N. benthamiana* (Supplementary Table 10). Among the 24 *NbACS* genes, three genes, *NbACS1, 2a* and *4* were upregulated by INF1 treatment. Four out of the 16 *NbACO* genes were categorized as INF1 inducible genes, of which *NbACO1a, 1b* and *4* were highly upregulated (Figure 6). Ethylene production was induced within 3 h after treatment of INF1, and gene silencing of *NbACO* genes (using a conserved region of *NbACO* genes) compromised INF1 induced production of ethylene (Supplementary Figure 2A). *NbACO*-silenced *N. benthamiana*, as well as *NbEIN2-silenced* plants, showed enhanced disease symptoms by *P. infestans* compared with the control plants (Supplementary Figure 2B), confirming the importance of ethylene production in disease resistance of *N. benthamiana*.

### *N. benthamiana* NbERF-IX-33 is Involved in the Production of Capsidiol and Resistance to *P. infestans*

The ERF (ethylene response factor) family transcription factors are known to be involved in the responses of plants to a number of environmental stresses (Müller and Munné-Bosch, 2015). Previous studies indicated that some ERF transcription factors can directly bind to a cis-acting element, typically the GCC box (AGCCGCC), to respond to pathogen attack (Ohme-Tagaki and Shinshi, 1995), while some studies also indicated that another member of the ERF transcription factors can bind to the dehydration responsive element (DRE, typically TACCGAC) to respond to drought stress (Yamaguchi-Shinozaki and Shinozaki, 1994). To investigate the activation of ethylene signaling during the defense induction in *N. benthamiana*, we created a complete list of predicted genes for AP2/ERF transcription factors found in the genome of *N. benthamiana*, using 147 *A. thaliana* AP2/ERF (Nakano et al., 2006) as queries for Blastp search against *N. benthamiana* predicted proteins (Sol genomics, Fernandez-Pozo et al., 2015). In the genome of *N. benthamiana*, we found 337 genes predicted to encode AP2/ERF transcription factors (Supplementary Tables 11 and 12). Based on the phylogenetic analysis of *A. thaliana* and *N. benthamiana AP2/ERF* genes, 47 genes were assigned to the AP2 family (including Soloist I) and 7 genes were classified in the RAV family (Figure 7 and Supplementary Table 11). The 283 *N. benthamiana ERF* genes were classified into 10 major groups (ERF-I to -X as in the case of *A. thaliana*, Nakano et al., 2006) and two small subgroups (Soloist II and III). Almost all groups of AP2/ERF transcription factors have corresponding members in Arabidopsis and *N. benthamiana*, suggesting that their functions are conserved in these two dicotyledonous plants. Among these, group ERF-IX contained the highest number of elicitor-responsive genes, and 22 out of the 34 genes (assigned to a cluster) were classified into INF1-induced clusters (Figure 7 and Supplementary Table 12). Because a substantial number of genes for NbERF were elicitor-inducible, it was expected that these NbERFs may have redundant functions in the induction of disease resistance.

**FIGURE 7.**
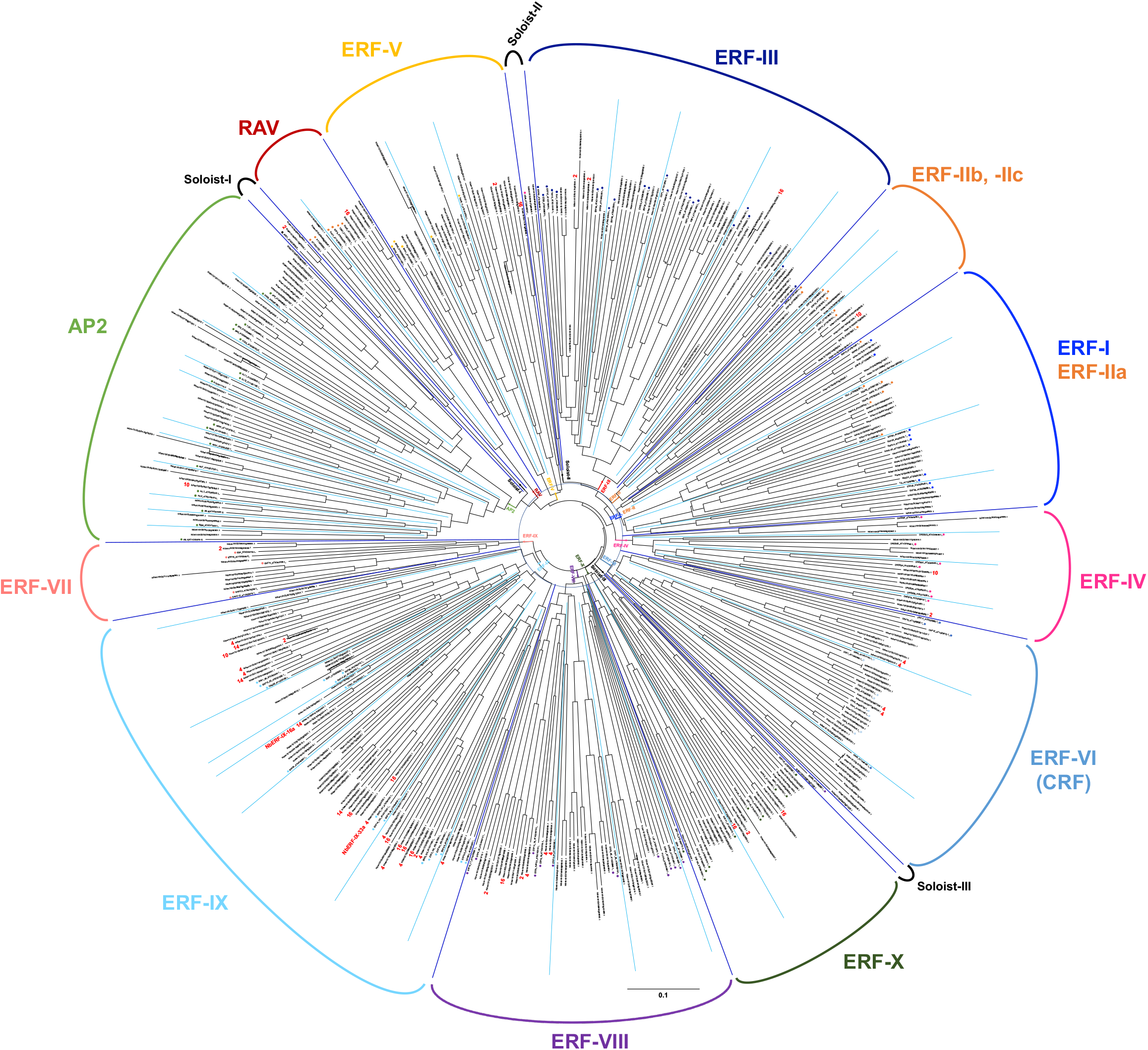
An unrooted phylogenetic tree of AP2/ERF transcription factors form *Nicotiana benthamiana* and *Arabidopsis thaliana*. The deduced amino acid sequences of AP2/ERF transcription factors were aligned by ClustalW (Thompson et al., 1994), and the phylogenetic tree was constructed using the neighbor-joining (NJ) method (Saitou and Nei, 1987). Classification of AP2/ERF transcription factors by Nakano et al. (2006) are indicated. The cluster numbers of INF1-inducible genes are shown in red letters.

To investigate the role of ERF-type transcription factors in the production of capsidiol during the induction of disease resistance, two *NbERF* were selected for further analysis in this study. *NbERF-IX-33a* (Niben101Scf00454g04003, cluster 4) whose expression level (FPKM value) was the highest among all *AP2/ERF* genes at 12 h after INF1 treatment, and *NbERF-IX-16a* (Niben101Scf01212g03005, cluster 14) whose expression level was the highest at 24 h after INF1 treatment (Figure 8A). For the specific silencing of these target genes, silencing vectors based on unique sequences of *NbERF-IX-33* and *NbERF-IX-16* were constructed. Both *NbERF-IX-33* and *NbERF-IX-16* occur in the form of highly similar *a* and *b* homologs, as described above. Due to the high sequence similarity of *a* and *b* homologs, a functional redundancy is to be expected and we therefore designed the silencing vectors to target the *a* and *b* homologs of both genes. Analysis using the SGN VIGS tool (Fernandez-Pozo et al., 2015) confirmed that the constructed vectors are specifically targeting *NbERF-IX-33* and *NbERF-IX-16* genes, and no potential off-target effect was detected.

**FIGURE 8.**
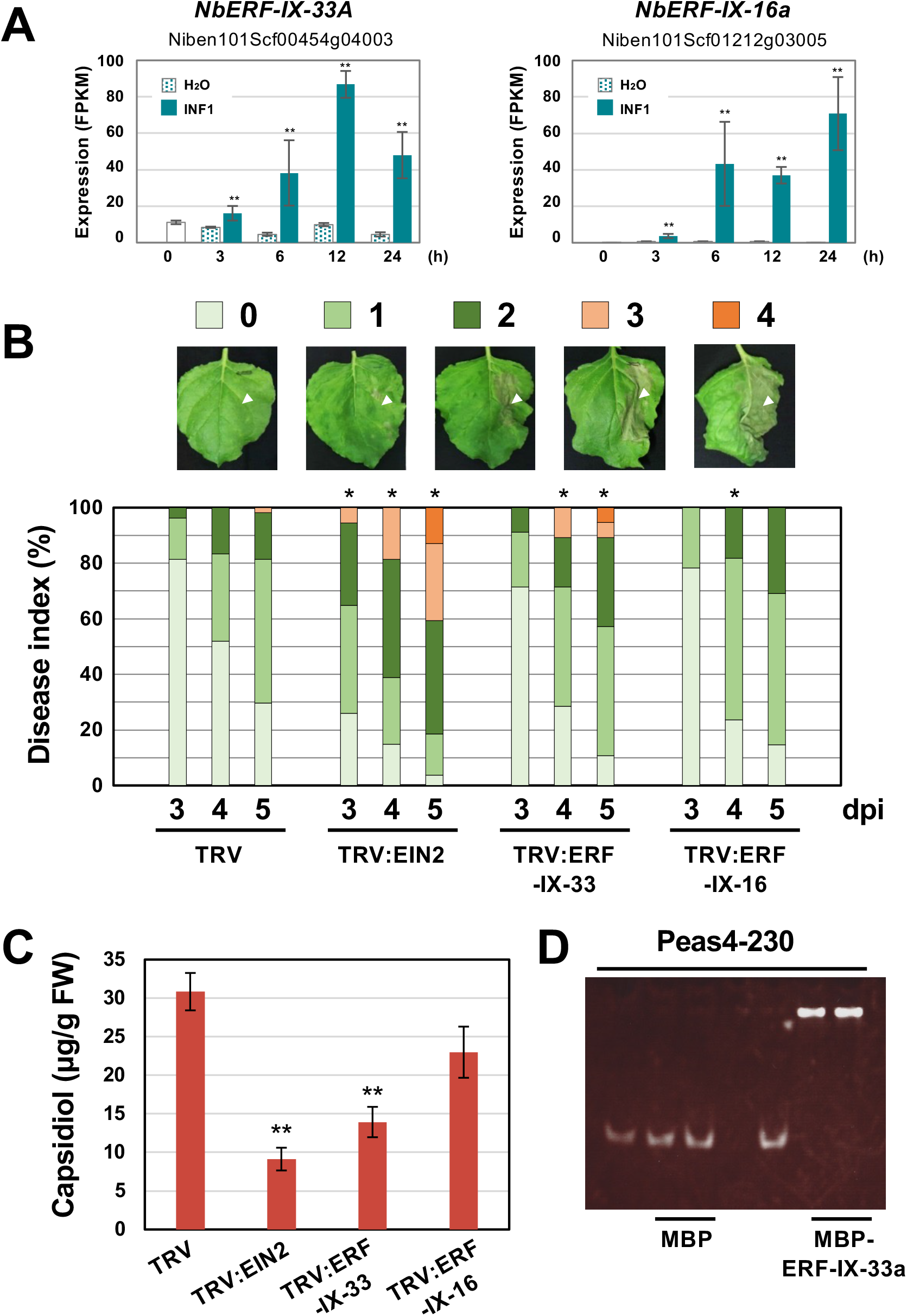
**(A)** Expression profiles of *Nicotiana benthamiana* genes encoding two selected ERF transcription factors. Gene expression (FPKM value) was determined by RNA-seq analysis of *N. benthamiana* leaves treated with water (H_2_O) or 150 nM INF1. Data are means ±SE (n = 3). Data marked with asterisks are significantly different from control (water-treated). \*\**P* < 0.01. **(B)** *N. benthamiana* were inoculated with TRV, TRV:ERF-IX-33 or TRV:ERF-IX-16 and the right side of leaves of control or gene-silenced plants were inoculated with *P. infestans*. The appearance of disease symptoms was categorized into 5 classes according to the severity of disease symptoms. 0, no visible symptom; 1, small wilted spots in inoculated area; 2, browning <50% of the inoculated side of leaf; 3, browning >50% of the inoculated side of leaf; 4, development of disease symptoms over central leaf vein. Plot showing percentage of *N. benthamiana* leaves with disease symptom severities represented in the five classes as shown in the upper panels, for leaves of control and gene-silenced plants inoculated with *P. infestans* from 3 to 5 days post-inoculation (dpi). At least 50 leaves from each control and gene-silenced plant were scored. Data marked with asterisks are significantly different from control as assessed by one-tailed Mann-Whitney U tests: \**P* < 0.05. **(C)** Production of phytoalexin in TRV-inoculated or gene-silenced *N. benthamiana*. Leaves were harvested 24 hours after 150 nM INF1 treatment and extracted phytoalexins were detected by HPLC. Data are means ±SE (n = 4). Data marked with asterisks are significantly different from control. \*\**P* < 0.01. **(D)** Binding of NbERF-IX33 to the *NbEAS4* promoter. MBP (maltose-binding protein) or MBP-ERF-IX-33a was incubated with *NbEAS4* promoter fragments. Samples were separated in a non-denaturing polyacrylamide gel. The gel was stained with SYBR Green EMSA stain.

*NbERF-IX-33* and *NbERF-IX-16*-silenced *N. benthamiana* plants were inoculated with *P. infestans* and disease symptoms on control and gene-silenced plants were scored as visible development of disease symptoms. Within the first 5 days after the inoculation of *P. infestans*, there was no obvious development of disease symptoms for control and *NbERF-IX-16*-silenced plants was detected, whereas *NbEIN2*- and *NbERF-IX-33*-silenced plants showed severe disease symptoms on the inoculated leaves (Figure 8B). To investigate the effect of the *NbERF* gene silencing of these *NbERF* on phytoalexin production, capsidiol was quantified in gene-silenced *N. benthamiana*. At 24 h after treatment with 150 nM INF1, capsidiol was extracted from control and gene-silenced *N. benthamiana* leaves, and the amount of capsidiol produced was quantified using HPLC. While capsidiol production in *NbERF-IX-16*-silenced *N. benthamiana* was not significantly different from that of the control (TRV) plant, capsidiol amounts in *NbERF-IX-33*- and *NbEIN2*-silenced plants were reduced (Figure 8C). These results indicated that *NbERF-IX-33* is a transcription factor essential for disease resistance through the production of capsidiol. The function of *NbERF-IX-33* appears to be partially compensated by other NbERF transcription factors in *NbERF-IX-33* silenced plants, given that both disease resistance and capsidiol production were more pronounced in *NbEIN2*-silenced plants.

To investigate whether NbERF-IX-33 acts directly on the *NbEAS* promoter, an electrophoresis mobility shift assay (EMSA) was conducted. To this end, we expressed and purified a recombinant fusion protein of maltose-binding protein (MBP) and NbERF-IX33 (MBP-NbERF-IX-33a) from *E. coli* (Supplementary Figure 3). The mobility shift assay was performed using the purified MBP-NbERF-IX-33a protein with *NbEAS4* promoter fragments. Mobility shift was observed for the 230 bp *NbEAS4* promoter fragment incubated with the NbERF-IX-33a protein, in contrast to the control experiment using MBP (Figure 8D). These results suggested that NbERF-IX-33 directly binds to the promoter region of *NbEAS4* to increase the production of phytoalexin production.

## DISCUSSIONS

Salicylic acid, jasmonic acid, and ethylene are commonly known as second messengers that play important roles in plant disease responses, but which plant hormones are essential for effective resistance induction varies among plant-pathogen combinations (Glazebrook, 2005). In general, salicylic acid plays an important role in the activation of defense against biotrophic and hemibiotrophic pathogens, while jasmonic acid and ethylene are usually implicated in the defense against necrotrophic pathogens. Since similarities in gene expression patterns have been noted between jasmonic acid and ethylene treatments in Arabidopsis (Schenk et al., 2000), both plant hormones are often regarded as having similar functions in disease resistance.

In the case of the interaction between *N. benthamiana* and hemibiotrophic *P. infestans*, it is presumed that ethylene, but not jasmonic acid, is the crucial plant hormone essential for the resistance against the pathogen. Although RNAseq analysis in this study detected the INF1-induced expression of some genes related to jasmonic acid production (data not shown), silencing of *NbCOI1*, a component of the jasmonic acid receptor, had no effect on the resistance of *N. benthamiana* to *P. infestans* (Shibata et al., 2010). The VIGS-based screening for defense-related genes identified six ethylene-related genes, but no gene involved in the production of jasmonic acid was isolated (Shibata et al., 2016). Thus, the importance of ethylene in *P. infestans* resistance of *N. benthamiana* is evident, but it has not been clarified how ethylene is involved in the induction of resistance.

In this study, we investigated the expression profiles of all *N. benthamiana* genes in leaves after treatment with INF1, a PAMP of oomycete pathogens. Among the genes induced by INF1, those in cluster 14 showed a particularly clear upregulation by INF1 treatment (Figure 2). The main gene group included in cluster 14 was PR (pathogenesis-related) protein genes, such as *PR-1, PR-2* (*β-1,3-glucanase*), *PR-3* (*chitinase*), *PR-4* and PR-5 (*Osmotin*) (Supplementary Table 6). In the previous VIGS screening for *N. benthamiana* genes required for the resistance against *P. infestans*, however, none of the genes for PR proteins were isolated from 3,000 randomly gene-silenced plants, in contrast, several genes involved in the capsidiol synthase had been identified (Shibata et al., 2016). Targeted gene silencing of *PR1* did not show any detectable effect on the resistance of *N. benthamiana* against *P. infestans* (Shibata et al., 2010). These results indicate that not all genes whose expression is induced by INF1 elicitor treatment necessarily function in *P. infestans* resistance. Nevertheless, cluster 14 contains genes for *NbEAS* and *NbEAH*, which are required for the resistance of *N. benthamiana* to *P. infestans*. The expression of *NbEAS* and *NbEAH* was markedly induced by INF1 treatment, however, the expression of these genes was not stimulated at all by water treatment. (Figure 3). Since there is a trade-off between the expression of genes for disease resistance and plant growth (Denancé et al., 2013), it is important to strictly control the expression of genes involved in plant defense. In fact, many genes for photosynthesis (such as *rbcS* and *Lhcb* genes) showed a tendency to be down-regulated in INF1-treated leaves (data not shown).

Promoter analysis of *NbEAS1* and *NbEAS4* revealed that the deletion of the GCC box-like sequence compromises the INF-induced expression of *NbEAS* genes (Figure 5), indicating the direct regulation of phytoalexin production by ERF. Previous reports described the presence of a GCC box in the promoter region of the pepper and tobacco *EAS* genes (Maldonado-Bonilla et al., 2008), but its role in regulating *EAS* expression had not been analyzed. This study proves that the GCC box-like motif is indispensable for the induction of the *NbEAS* gene, but does not rule out the possibility that other cis sequences are included in the promoter region. W-box (C/T)TGCA(C/T) motifs, previously shown to correlate with the defense induction in *N. benthamiana* (Ishihama et al., 2011), are found in the promoter region of some *NbEAS* and *NbEAH* genes (Supplementary Figure 1). We also noticed that when we shortened the length of the *NbEAS1* promoter from 400 to 300 (300 to 260 in *NbEAS4*) for the analysis of promoter activities, the basal expression level decreased. This region may therefore contain sequences with which auxiliary transcriptional regulators interact to increase the overall amount of expression. The promoter regions of *NbEAS3* and *NbEAS4* have two GCC-box like motifs (Supplementary Figure 1). *NbEAS3* and *NbEAS4* are the most highly expressed homologs of *NbEAS* (Figure 3), thus this second GCC-box-like sequence might be responsible for the stronger INF1-induced gene expression.

In control samples we observed a constant expression of MVA pathway genes, which was to be expected, since besides producing the capsidiol precursor FPP, the pathway is of central importance for a large variety of isoprenoid compounds. A quick and ample production of capsidiol would require the MVA pathway to step up production, which appears to be the case, as INF1 elicitation could be shown to up-regulate several MVA genes (Figure 4). Despite this, the GCC-box-like motif was not found in the promoters of MVA genes, not even *NbHMGR2*, which was contained in cluster 14 with several *NbEAS* genes. This is somewhat surprising, since the upregulation of the MVA pathway during pathogenesis is typically not observed in plants that do not produce terpenoid defense compounds, and we would therefore assume that this regulatory mechanism had co-evolved with terpenoid phytoalexin synthesis. An explanation of why the GCC-box motif is absent in MVA promoters is purely speculative at this point, but one reason could be that terpenoid phytoalexins are preferentially produced against necro- and hemibiotrophic pathogens, and the MVA pathway may need to be tightly regulated during defense against biotic pathogens as well. Ishihama et al. (2011) has reported that *NbHMGR2* is under the control of a defense-related transcription factor, NbWRKY8, which targets W-boxes, but further investigation of the activation mechanism of genes in the MVA pathway would be an interesting subject of future research.

AP2/ERF family is a conserved group of plant transcription factors, being defined by a central AP2/ERF domain consisting of approx. 60 amino acid residues, which bind to cis-element targets (Licausi et al., 2013). Although they are called “ethylene-responsive factors” due to how they were first discovered (Ohme-Tagaki and Shinshi, 1995), ERFs are involved in the regulation of diverse phenomena, including development, morphogenesis, and biotic/abiotic stress responses mediated via different plant hormones. For instance, Arabidopsis PUCHI (belonging to subfamily ERF-VIII) is involved in lateral root initiation and development via auxin-mediated signaling (Hirota et al., 2007), while CRFs (cytokinin response factors, belonging to ERF-VI) are induced by cytokinin, which functions in the regulation of root growth, embryo development, leaf senescence, and hypocotyl elongation (Kim, 2016). In this study, we listed all genes predicted to encode AP2/ERF transcription factors in the *N. benthamiana* genome to identify the transcription factors that directly control phytoalexin production via GCC-box-like sequences in *NbEAS1* and *NbEAS4* promoters. A total of 283 *ERF* genes were found, of which 43 (15.2 %) were assigned to the cluster for INF1-inducible genes (Supplementary Table 12). Given that INF1 treatment enhanced the production of ethylene within 9 h (Supplementary Figure 2), a substantial number of ERF gene groups were shown not to be induced by ethylene.

The functional analysis of two NbERFs (both belonging to ERF-IX), whose expression was significantly increased by INF1 treatment, indicated that NbERF-IX-33 is an essential transcription factor for the INF1-induced production of capsidiol and the resistance to *P. infestans* (Figure 8). Gene silencing of *NbERF-IX-16* may reduce the capsidiol production (this could not be shown to be statistically significant), and the *NbERF-IX-16*-silenced plant were slightly more susceptible to *P. infestans* (Figure 8). The reduction in disease resistance and capsidiol production in *NbEIN2*-silenced plants was more pronounced than in *NbERF-IX-33*-silenced plants, suggesting that EIN2 may act upstream of NbERF regulation, affecting multiple NbERFs involved in the induction of phytoalexin production.

In this study, we reported that NbERF-IX-33a binds to the GCC-box like motif in the promoter region of *NbEAS4*, which is supported by many studies that have shown ERFs that bind to the GCC box. Four ERFs were originally isolated from tobacco as transcription factors that directly bind to the GCC box, the conserved motif found in the promoter region of ethylene-inducible *PR* genes (Ohme-Takagi and Shinshi, 1995). In Arabidopsis, ORA59 and ERF1/ERF92 (subfamily IX) have been shown to activate jasmonic acid- and ethylene-mediated expression of the defensin gene *AtPDF1.2* by directly binding to two GCC boxes in its promoter region (Zarei et al., 2011), while AtERF3 and AtERF4 can downregulate their target gene via a GCC box (Fujimoto et al., 2000). In rice, OsERF83 positively regulates the disease resistance against rice blast pathogen by upregulating *PR* genes, whereas OsERF922 activates abscisic acid biosynthesis-related genes which has a negative effect on the disease resistance (Liu et al., 2012; Tezuka et al. 2019). Tobacco ERF189 and four related ERFs (subfamily IX) can bind to a GCC-box in the promoter region of *PMT* (putrescine N-methyltransferase) for jasmonate-inducible nicotine synthesis, and all enzyme genes for nicotine production are under the control of ERF189 (Shoji et al., 2010). Wheat ERF transcription factor TaERF3 is involved in salt and drought tolerance via the GCC boxes in stress-related genes (Rong et al., 2014). Expression of genes for the production of anti-insect steroidal glycoalkaloids (SGA) in tomato is under the control of jasmonate-responsive ERF, JRE4, which bind to GCC box-like elements in the promoter of SGA biosynthetic genes (Thagun et al., 2016). These reports indicate that GCC-box and ERF combinations are involved in the regulation of diverse phenomena in different plant species.

The synthesis of bioactive compounds requires a high degree of coordination, especially in cases where other pathways rely on the same substrate (such as FPP), and where its depletion can therefore have widespread negative effects. This poses a problem during the evolution of a novel pathway, since it must assure that other pathways are not disrupted. Shoji and Yuan (2021) suggest that the diversified combinations of a collection of enzyme genes under the control of the same transcriptional factors can lead to the invention of a new metabolic pathway in different plant species. In addition to the promoters of genes for NbEAS and NbEAH, which consist of a short metabolic pathway for capsidiol production, the GCC-box like motifs are also conserved in the promoter region of *NbABCG1* and 2 genes, which is involved in the secretion of capsidiol at the sites of pathogens attack (Shibata et al., 2016; Rin et al., 2017). Thus, this theory may be valid beyond the enzyme genes of the metabolic pathways. Rishitin, a sesquiterpenoid phytoalexin produced by potato and tomato, is expected to be produced by a more complex metabolic pathway compared with capsidiol, but the pathway for rishitin production is still largely unknown. Since the two primary rishitin-producing enzymes also share a GCC box-like promoter sequence (Takemoto et al., 2018), the similarity of expression patterns and the commonality of the regulating transcription factors may provide a clue to the elucidation of this yet unknown biosynthetic pathway.

## Author contributions

DT designed the research. SI, MF, AT-K, HM, AA, AT and DT conducted the experiments. SI, MF, MC, AA and DT analyzed data, A Tanaka, IS, SC, KK, MO and DT supervised the experiments. SI and DT wrote the manuscript. MC and DT edited the manuscript. SI, MC and DT contributed to the discussion and interpretation of the results.

## Acknowledgments

We thank Prof. David C. Baulcombe (University of Cambridge, USA) for providing pTV00 and pBINTRA6 vectors, Prof. Sophien Kamoun (The Sainsbury Laboratory, UK) for the pFB53 vector, and Prof. Gregory B. Martin (Cornell University, USA) for the access to the *N. benthamiana* genome database. We also thank Dr. David Jones (The Australian National University, Australia) for *N. benthamiana* seeds, Ms. Kayo Shirai (Hokkaido Central Agricultural Experiment Station, Japan), and Dr. Seishi Akino (Hokkaido University, Japan) for providing *P. infestans* isolate 08YD1. We also thank Dr. Kenji Asano and Mr. Seiji Tamiya (National Agricultural Research Center for Hokkaido Region, Japan) and Mr. Yasuki Tahara (Nagoya University, Japan) for providing tubers of potato cultivars.

## SUPPLEMENTAL DATA

**SUPPLEMENTARY FIGURE 1 |** Nucleotide sequence of the promoter region of *NbEAS* and *NbEAH* genes. The 7-bp sequences for the GCC box-like motifs are shown in red letters, the 6-bp sequences for the W-box motifs are shown in blue letters, and the start codon of *NbEAS* and *NbEAH* are shown in orange letters. Note that the promoter sequence for *NbEAS10* (−350 to −237) is not available.

**SUPPLEMENTARY FIGURE 2 | (A)** Accumulation of ethylene in *N. benthamiana* leaves treated with INF1. Leaves of control (TRV) or *NbACO*-silenced (TRV:ACO) were treated with water (H_2_O), or 150 nM INF1 and the amount of ethylene produced was measured by gas chromatography. Data are means ± SE (n = 4). **(B)***N. benthamiana* were inoculated with TRV, TRV:EIN2 or TRV:ACO and leaves of control or *NbACO*-silenced plants were inoculated with *P. infestans*. The appearance of disease symptoms was categorized into 3 classes according to the severity of disease symptoms. 0, no visible symptom; 1, small wilted spots in inoculated area; 2, browning >50% of the inoculated side of the leaf. Plot showing percentage of *N. benthamiana* leaves with disease symptom severities represented in the three classes as shown in the upper panels, for leaves of control and gene-silenced plants inoculated with *P. infestans* at 7days post-inoculation (dpi). At least 12 leaves from each control and gene-silenced plant were scored. Data marked with asterisks are significantly different from control as assessed by one-tailed Mann-Whitney U tests: \**P* < 0.05.

**SUPPLEMENTARY FIGURE 3 |** Expression and purification of NbERF-IX-33a proteins in *Escherichia coli. E. coli* with pMAL-c5x or pMAL-c5x containing *NbERF-IX-33* gene were cultured in LB medium with IPTG for the induction of protein expression. Cultured *E. coli* cells were harvested 5 hours after IPTG treatment and MBP (maltose-binding protein) or MBP-NbERF-IX-33a were purified using amylose resin. Eluted fractions were separated by SDS-PAGE and stained with CBB. Protein size markers are shown in kDa. The concentrations of the purified proteins were adjusted and used for the experiments in Figure 8D.

**Supplementary Table 1** Plasmids used in this study.

**Supplementary Table 2** Primers used in this study.

**Supplementary Table 3** *Nicotiana benthamiana* genes categorized in cluster 2.

**Supplementary Table 4** *Nicotiana benthamiana* genes categorized in cluster 4.

**Supplementary Table 5** *Nicotiana benthamiana* genes categorized in cluster 10.

**Supplementary Table 6** *Nicotiana benthamiana* genes categorized in cluster 14.

**Supplementary Table 7** *Nicotiana benthamiana* genes categorized in cluster 16.

**Supplementary Table 8** Predicted *Nicotiana benthamiana* genes for enzymes specifically involved in the production of capsidiol.

**Supplementary Table 9** Predicted genes for enzymes in the mevalonate pathway and farnesyl-pyrophosphate synthase in *Nicotiana benthamiana*.

**Supplementary Table 10** Predicted genes for enzymes involved in the methionine cycle and ethylene production in *Nicotiana benthamiana*.

**Supplementary Table 11** Gene list for predicted AP2 (RAV) family transcription factors in *Nicotiana benthamiana*.

**Supplementary Table 12** Gene list for predicted AP2 (RAV) family transcription factors in *Nicotiana benthamiana*.

