## Supplemental Figures and Tables for "AP2/ERF transcription factor NbERF-IX-33 is involved in the regulation of phytoalexin production for the resistance of *Nicotiana benthamiana* to *Phytophthora infestans*"

### P_*NbEAS1*

-350 ATTTAACTGCTACTTTAAATAATACTATCCACTTAACCAAAAAATGAAAA
ATAAAGTACATAAACTTTAAATAATAGGGAAATTTGGATCAACAGACCCC
-250 AGACGCCAAGAATGAATTAATAGGCTGCTGGTTGGCTGACTAGCTAGTTA
GTGTAAAGTCAAGTAAGGCAACTGGGAAATGATTAGTTGTTTAAATAATT
-150 GGCTGCACTTTTCTCACAACTATATATATATATATATACACTTGTCCCTT
CTCTTCCATTCAAATCATCAGCAATTCAGAGTTCCTAATTTCCTTCTTCC
-50 TTAAAACGAACAAAAACAATACCCTCATCTTTTAATTTATTAGCAATATA
1 ATG

### P_*NbEAS2*

-350 ATTTAACTGCTACTTTAAATAATACTATCCACTTAACCAAAAAATGAAAA
ATAAAGTACATAAACTTTAAATAATAGGGAAATTTGGATCAACAGACCCC
-250 AGACGCCAAGAATGAATTAATAGGCTGCTGGTTGGCTGACTAGCTAGTTA
GTGTAAAGTCAAGTAAGGCAACTGGGAAATGATTAGTTGTTTAAATAATT
-150 GGCTGCACTTTTCTCACAACTATATATATATATATATACACTTGTCCCTT
CTCTTCCATTCAAATCATCAGCAATTCAGAGTTCCTAATTTCCTTCTTCC
-50 TTAAAACGAACAAAAACAATACCCTCATCTTTTAATTTATTAGCAATATA
1 ATG

### P_*NbEAS3*

-350 TAACTAAGGTGGCTAGACATTTATAATAAAGAGTAGTTAGTGTAAACATT
ATTTTAGAGACGTAGGATGTCCCACAACACAAAATTTTTTATGTAAGAAA
-250 TACTGCTCCTATTATACTGGATCAATTGAGACGCCAAGAAAGAAATCAAA
AGACGCCAAGGAAGAAATATTGTATCAGTAGACTACAGTCAAGTAAGGCA
-150 ACTGAAATGAAGAAATAAAAAAACACTATAAATACTTATGCCTTCTCTTC
CATTTGGGTCATCAGTCTACTTTCTTTTTCTTCCTCGGAGAATTAAGAAG
-50 CCAAAAATTCTCCTATCATTTGTGGTACTAAGAGTATAAATTCTATAGCA
1 ATG

### P_*NbEAS4*

-350 TAACTAAGGTGGCTAGACATTTATAATAAAGAGTAGTTAGTGTAAACATT
ATTTTAGAGACGTAGGATGTCCCACAACACAAAATTTTTTATGTAAGAAA
-250 TACTGCTCCTATTATACTGGATCAATTGAGACGCCAAGAAAGAAATCAAA
AGACGCCAAGGAAGAAATATTGTATCAGTAGACTACAGTCAAGTAAGGCA
-150 ACTGAAATGAAGAAATAAAAAAACACTATAAATACTTATGCCTTCTCTTC
CATTTGGGTCATCAGTCTACTTTCTTTTTCTTCCTCGGAGAATTAAGAAG
-50 CCAAAAATTCTCCTATCATTTGTGGTACTAAGAGTATAAATTCTATAGCA
1 ATG

### P_*NbEAS5*

-350 CAATTAAATCTCGTCAATTTATTTTTAACCAATCCAAATAAGAAGTTGGA
GAAAGAAAAGACTAAGAAATAGAAGGGAGTTGAATACTCTTAACTAAAGA
-250 GTAGTTACTAGGAAAGTAAACATTGACACAATAAAAAATATCCGTAAGAA
ATACTACTACAATTATATTGGATTCACTGAGACGCCAATAAAGAAATCAA
-150 AAGACGCTGCTGATGATTGGTATATATGGTCAAGTAAGGCAACTGAAATG
AAGAAATAAAAAAGTATTATAAATACTTATGCCTTCTCTACCAATTGTGT
-50 CATCACTCAGAGAATTAATAACACAAAATTCTCCTATAACTTTTATAGCA
1 ATG

### P_*NbEAS6*

```
-350 TCTTAATTAAGGCGGCTAGAAATTTAAAGTTTATAATAAAGACCAGTTAC
     TACTATATATTATTTTCCCACGACACTAATTTTTTTTCCGTAAGAAATAA
-250 TACTACTATTCAATGAGACACCAATAAAGAAATCAAAAAGCTGGTGATGA
     GTGGTATATATGGTCAAGTAAGGCAACTGAAATGAAGAAATAAAAAAGCA
-150 TTATAAATACTTATGCCTTCTCTACCAATTGTGTCATCACTCAGAGAATT
     AATAACACAAAATTCTCCTATAACTTCTATAGCAATGGCCGCAGCAGCAG
-50  TTGGCANCAGAGAATTAATAACACAAAATTCTCCTATGACTTTTATAGCA
1    ATG
```

### P_*NbEAS7*

```
-350 TCAATTAATTTCCAACGGAGAGACTAAGAAATGGAAGAGAGTTGAATACT
     CTTAATTAAGGCGGCTAGATATTTAAAATTTATAATAAAGACCAGTTACT
-250 ACTATATATTATTTGCCCACGACACAAAATTTTTTTGCGTAAGAAATAAT
     ACTACTATTCAATGAGACGCCAATAAAGAAATCAAAAAGCTGGTGATGAG
-150 TGGTATATATGGTCAAGTAAGGCAACTGAAATGAAGAAATAGAAAAGCAT
     TATAAATACTTATGCCTTCTCTACCAATTGTGTCATCACTCCACTTTCTT
-50  TCTTCCCCAGAGAATTAATAACACAAAATTCTCCTATGACTTTTATAGCA
1    ATG
```

### P_*NbEAS8*

```
-350 CGTTTATAATAAAGAGTTGTTACTGGGAAAGTAAACATTATTGTATGATG
     TCCCACAACACAATTTATTTTTAAGTAAGAAATACTACTCGTATTATATT
-250 GGATCAATTGAGACGCCAAGAAAGAAATCAAAAGACGCCAATGAAGAAAT
     ATTGGAACAGTAGACTATATGATATATAGTCAAGTAAGGCAACTGAAATG
-150 AAGAAATAAAAAAATCACTATAAATACCTATGCGTTCTCTTCCGTTTGGG
     TCATCTCACTCATCAGTCTACTTTCTTTCTTCCTCGGAGAATTAAGAACC
-50  AAAAAATTCTCTTATCATTTGTGGTACTAAGAGTATAACTTTTATTAGCA
1    ATG
```

### P_*NbEAS9*

```
-350 GCAATTTGGCTATTTAAATAATATTATTCAGTTTACGAAAAAAGAATAAA
     GTAATAGACTTTAAATAATTGAGAAACTGGATGAATAGACCACAGACGCC
-250 AACAATGAATCAAAAGGCTGCTGCCTAATGTAAAGTCAAGTAAGACAACT
     GGGAATTTGATTAGGTGATTTTGATCAGTTAAAATATCAACGTGACGTAG
-150 TTGTTTTAAATAATAATTACATGAACAAAAATTAGAGAATTGGCTTTGGG
     TCTCTCACCACTATATATACTTGTCCCTTGCCTTCCATTTAAGTCAACAA
-50  CTGAGAGTTCCTTAAAATAACAAAAACAATACTCTCATCTTTTATTAGCA
1    ATG
```

### P_*NbEAS10*

```
-350 --------------------------------------------------
     --------------------------------------------------
-250 --------------CGCCCCGTCGCCGACTTCTCCCCTAGTCTTTGGGGT
     GATCAGTTCCTTTCATTCTCCATTGACAATCAGGTAATTTATACTTTCTT
-150 CGTGCAAGCATGCATGGTTTATTTCTTCTTTGTATTTATTGAAAGACATT
     AATTTTGCTAACTTCACTTTTCTTTTTTTAAATACGTTATAAATAGGTTG
-50  CAGAAACGTATGCTAAAGAGATTGAAGCACTGAAGGAGCAAACGAGGAAT
1    ATG
```

### P_*NbEAH1*

-350 AAGTATTTAGTGAGGAGTTAGTATCAAATTTGTAGCCGCCAAATATGAAA
GTCTTTGTGGCTATACTTTCGGATTATATAGAAAAAAAGAATATATTGCT
-250 TCACAATAATCAGTCCACAATACTAATAATTAATTACATCATTATTGCTC
TCTAGTCAGCTGAATATGAATAATTAATGGTCAACAGGAAAGTAAACAAA
-150 GTCTGCTTCAATGAGATGTTGATCAACAAAAGTTGCAGAAAATATTAGTA
TATTTATCTTTCCATTTCTTGCCAAAACAAATGCCTATAAAAGACCATAC
-50 GCATCCAGACTTTGAAGAATCATCGAATAATCCTCCATTTATCTCCGAAA
1 ATG

### P_*NbEAH2*

-350 TTTTATCCATATTTTACCCATTTTTAAAAATTTAGTCGTCCAACCCATTT
TTGGTGAATTATATTAATGGATAATTATTTTTTTAAATCCATTTCGCCGC
-250 CGCTACTCCGAGATTATTAGAGAATTTCCCCCGCATTGCTGTTCTTTGAG
TCAGCTGAGTTGTTTTGACTCTGAAGGAAGTTGAACAGTTAAGTAAACAA
-150 AGTCTGCTTTGACAAACAAATGTTGCAGAAAAATACTACTCTTTCTTTGC
ATATCCCATTCTGGCCAAAACAAAAGCCTATAAAAGACCACACACAACTG
-50 ACCTTTGAATTATATCATCGAACCAAATACTCCTCCATTTATCTCCCAAA
1 ATG

### P_*NbEAH3*

-350 AAGTATTTAGTGAGGAGTTAGTATCAAATTTGTAGCCGCCAAATATGAAA
GTCTTTGTGGCTATACTTTCGGATTATATAGAAAAAAAGAATATATTGCT
-250 TCACAATAATCAGTCCACAATACTAATAATTAATTACATCATTATTGCTC
TCTAGTCAGCTGAATATGAATAATTAATGGTCAACAGGAAAGTAAACAAA
-150 GTCTGCTTCAATGAGATGTTGATCAACAAAAGTTGCAGAAAATATTAGTA
TATTTATCTTTCCATTTCTTGCCAAAACAAATGCCTATAAAAGACCATAC
-50 GCATCCAGACTTTGAAGAATCATCGAATAATCCTCCATTTATCTCCGAAA
1 ATG

### P_*NbEAH4* (Niben101Scf00072g05004.1)

-350 AAGTATTTAGTGAGGAGTTAGTATCAAATTTGTAGCCGCCAAATATGAAA
GTCTTTGTGGCTATACTTTCGGATTATATAGAAAAAAAGAATATATTGCT
-250 TCACAATAATCAGTCCACAATACTAATAATTAATTACATCATTATTGCTC
TCTAGTCAGCTGAATATGAATAATTAATGGTCAACAGGAAAGTAAACAAA
-150 GTCTGCTTCAATGAGATGTTGATCAACAAAAGTTGCAGAAAATATTAGTA
TATTTATCTTTCCATTTCTTGCCAAAACAAATGCCTATAAAAGACCATAC
-50 GCATCCAGACTTTGAAGAATCATCGAATAATCCTCCATTTATCTCCGAAA
1 ATG

### P_*NbEAH5*

-350 CTAATCCTTTGTTACTACCAAATTTCTGGCCCCTTAGTTAAGAGTCTTTC
CCGTCCGCAGATTACTAGGAATTTGCCAGAACAGTGATAAAATATTCATA
-250 TTTTGGCTTAAATATTACATCATTATGGTTCTTTTAGTCAGCTGTACTGT
TTTGACCCTTGAGAAAAATTAGTTCCACAGAAAAGTAAACAAAGTCTGCT
-150 TTGATCAGATTTTGAAATATATATATATATATCACTATTGCTTTGTATCT
GCATATTCCATCTCTTGCCAAACCTGCAGCCTATAAAAGAGTATATATAT
-50 GCAACTGAGCTTTGAAGAATACCAACTAACCCTCCCTTTCTCTCCCAATT
1 ATG

### P_*NbEAH6*

-350 TCCCTATTCTCTCTCTATATTGTTCTTATTCTTGCATTATTGTTCTTGTC
TTTATATTATTTCATAACATGATTAAAATATTAAATAGTTCTCACAAAGT
-250 TTTGTATTTTTTATAGCCAGCTCAAATATTCCTCTGTCATATTTATTAGTT
TTGACTAGAAAATTCCCATTTTCTTGAGAAAAATTATTCAATTGTTTAGT
-150 TTGTTTTCTTCTTTATTAAAATTATGAAAAGCTACAGCTGCCATGGTTGA
CAGCTGAATAAACAAAGTTGCTGAAAAATGCATCTGTCATTTAAAATGAG
-50 CATAAATACCCAACAACACTTTGAAGAATCTATCAAACCAAATAACCAAA
1 ATG

**SUPPLEMENTARY FIGURE 1 |** Nucleotide sequence of promoter region of *NbEAS* and *NbEAH* genes. The 7-bp sequence for the GCC box-like motifs are shown in red letters, 6-bp sequence for the W-box motifs are shown in blue letters, and the start codon of *NbEAS* and *NbEAH* are shown in orange letters. Note that the promoter sequence for *NbEAS10* (-350 to -237) was not available.

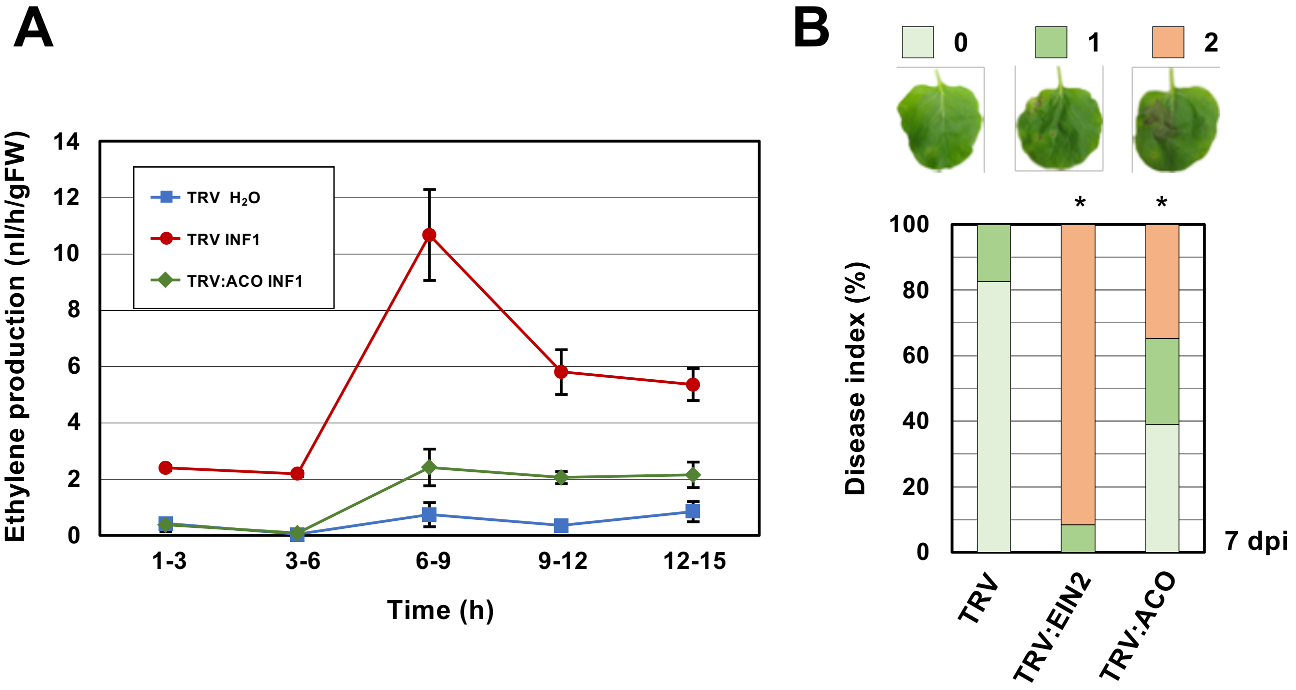


**SUPPLEMENTARY FIGURE 2 | (A)** Accumulation of ethylene in *N. benthamiana* leaves treated with INF1. Leaves of control (TRV) or *NbACO*-silenced (TRV:ACO) were treated with water ($H_2O$), or 150 nM INF1 and the amount of ethylene produced was measured by gas chromatography. At least 5 samples from each control and gene-silenced plants were scored for all time points. **(B)** *N. benthamiana* were inoculated with TRV, TRV:EIN2 or TRV:ACO and leaves of control or *NbACO*-silenced plants were inoculated with *P. infestans*. The appearance of disease symptoms was categorized into 3 classes according to the severity of disease symptoms. 0, no visible symptom; 1, small wilted spots in inoculated area; 2, browning >50% of the inoculated side of the leaf. Plot showing percentage of *N. benthamiana* leaves with disease symptom severities represented in the three classes as shown in the upper panels, for leaves of control and gene-silenced plants inoculated with *P. infestans* at 7 days post inoculation (dpi). At least 12 leaves from each control and gene-silenced plants were scored. Data marked with asterisks are significantly different from control as assessed by one-tailed Mann-Whitney U tests: $*P < 0.05$.

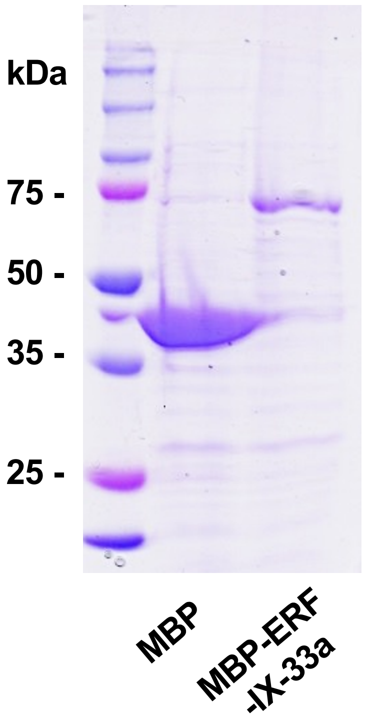


**SUPPLEMENTARY FIGURE 3 |** Expression and purification of NbERF-IX-33a proteins in *Escherichia coli. E. coli* with pMAL-c5x or pMAL-c5x containing *NbERF-IX-33a* gene were cultured in LB medium with IPTG for the induction of protein expression. Cultured *E. coli* cells were harvested 5 hours after IPTG treatment and MBP (maltose-binding protein) or MBP- NbERF-IX-33a were purified using amylose resins. Eluted fractions were separated by SDS-PAGE and stained with CBB. Protein size markers are shown in kDa. The concentrations of the purified proteins were adjusted and used for the experiments in Figure 8D.

**Supplementary Table 1.** Plasmids used in this study.

| Vector name | Base vector | Restriction site | Insert | Primers used to amplify insert | References | note |
|---|---|---|---|---|---|---|
| **Base vectors** | | | | | | |
| pNPP40-GFP | - | - | - | - | Shibata et al. 2016 | Base vector for *Agrobacterium* mediated expression of GFP; Kan[R] |
| pBINTRA6 | - | - | - | - | Ratcliff et al. 2001 | VIGS vector encoding TRV RNA1; Kan[R] |
| pTV00 | - | - | - | - | Ratcliff et al. 2001 | VIGS vector encoding TRV RNA2 with multi cloning site; Kan[R] |
| pMAL-c5x | - | - | - | - | New England Biolabs | Base vector for the expression of MBP fusion protein in *E. coli* ; Amp[R] |
| pNPP243 | pNPP40-GFP | - | - | EGFP-F, pNPP40-R [a] | RIN et al. 2017 | Linealized vector for promoter:*GFP* constructs; Kan[R] |
| **Plasmids for promoter analysis** | | | | | | |
| pNPP40-INF1 | - | - | - | - | Shibata et al. 2016 | Vector for *Agrobacterium*- mediated expression of INF1 |
| pNPP40-tGUS | - | - | - | - | RIN et al. 2017 | Vector for *Agrobacterium*- mediated expression of truncated *GUS* gene. |
| pNPP243:P_NbEAS1-400:GFP | pNPP243 | Linealized by PCR | P_*NbEAS1-400* | IF_Peas1-400-F, IF_Peas1-R | This study | Expression vector for *GFP* under control of 400-bp *NbEAS1* promoter. |
| pNPP243:P_NbEAS1-300:GFP | pNPP243 | Linealized by PCR | P_*NbEAS1-300* | IF_Peas1-300-F, IF_Peas1-R | This study | Expression vector for *GFP* under control of 300-bp *NbEAS1* promoter. |
| pNPP243:P_NbEAS1-260:GFP | pNPP243 | Linealized by PCR | P_*NbEAS1-260* | IF_Peas1-260-F, IF_Peas1-R | This study | Expression vector for *GFP* under control of 260-bp *NbEAS1* promoter. |
| pNPP243:P_NbEAS1-230:GFP | pNPP243 | Linealized by PCR | P_*NbEAS1-230* | IF_Peas1-230-F, IF_Peas1-R | This study | Expression vector for *GFP* under control of 230-bp *NbEAS1* promoter. |
| pNPP243:P_NbEAS1-200:GFP | pNPP243 | Linealized by PCR | P_*NbEAS1-200* | IF_Peas1-200-F, IF_Peas1-R | This study | Expression vector for *GFP* under control of 200-bp *NbEAS1* promoter. |
| pNPP243:P_NbEAS4-400:GFP | pNPP243 | Linealized by PCR | P_*NbEAS4-400* | IF_Peas4-400-F, IF_Peas4-R | This study | Expression vector for *GFP* under control of 400-bp *NbEAS4* promoter. |
| pNPP243:P_NbEAS4-300:GFP | pNPP243 | Linealized by PCR | P_*NbEAS4-300* | IF_Peas4-300-F, IF_Peas4-R | This study | Expression vector for *GFP* under control of 300-bp *NbEAS4* promoter. |
| pNPP243:P_NbEAS4-260:GFP | pNPP243 | Linealized by PCR | P_*NbEAS4-260* | IF_Peas4-260-F, IF_Peas4-R | This study | Expression vector for *GFP* under control of 260-bp *NbEAS4* promoter. |
| pNPP243:P_NbEAS4-230:GFP | pNPP243 | Linealized by PCR | P_*NbEAS4-230* | IF_Peas4-230-F, IF_Peas4-R | This study | Expression vector for *GFP* under control of 230-bp *NbEAS4* promoter. |
| pNPP243:P_NbEAS4-200:GFP | pNPP243 | Linealized by PCR | P_*NbEAS4-200* | IF_Peas4-200-F, IF_Peas4-R | This study | Expression vector for *GFP* under control of 200-bp *NbEAS4* promoter. |
| pNPP243:P_NbEAS4-230 (TT):GFP | pNPP243 | Linealized by PCR | P_*NbEAS4-230 (TT)* | IF_Peas4-230TT-F, IF_Peas4-R | This study | Expression vector for *GFP* under control of 230-bp *NbEAS4* promoter with mutations in GCC box. |
| **Plasmids for VIGS** | | | | | | |
| pTV00-EIN2 | pTV00 | - | - | - | Shibata et al. 2010 | Silencing vector for *NbEIN2* |
| pTV00-ACO | pTV00 | *Bam* HI/*Sma* I | *NbACO2a* (Partial, 385 bp) | NbACO2-BamHI-F, NbACO2-SmaI-R | This study | Silencing vector for *NbACO* genes |
| pTV00-NbERF-IX-33 | pTV00 | *Sma* I | *NbERF-IX-33* (Partial, 300 bp) | IF_pTV00_Nb36116-F, IF_pTV00_Nb36116-R | This study | Silencing vector for *NbERF-IX-33* |
| pTV00-NbERF-IX-16 | pTV00 | *Sma* I | *NbERF-IX-16* (Partial, 300 bp) | IF_pTV00_Nb25008-F, IF_pTV00_Nb25008-R | This study | Silencing vector for *NbERF-IX-16* |
| **Plasmid for protein expression in E. *coli*** | | | | | | |
| pMAL-NbERF-IX-33a | pMAL-c5x | *Eco* RV | *NbERF-IX-33a* | IF_pMAL-36116-F, IF_pMAL-36116-R | This study | Expression of MBP-NbERF-IX-33a fusion protein in *E. coli* |

[a] Primers used to amplify linierlized vector.

**Supplementary Table 2.** Primers used in this study.

| Primer | Sequence | Note |
|---|---|---|
| **Primers for construction of expression vectors for promoter analysis** | | |
| EGFP-F | ATGGTGAGCAAGGGCGAGGA | for amplification of pNPP243 |
| pNPP40-R | CTAGATGTAGTTGTAGAATG | for amplification of pNPP243 |
| IF_Peas1-400-F | ACCCTCACTAAAGGGAAATTTCGGCTTCTAATAGT | for construction of pNPP243:P_NbEAS1-400:GFP |
| IF_Peas1-300-F | ACCCTCACTAAAGGGATAAAGTACATAAACTTTAA | for construction of pNPP243:P_NbEAS1-300:GFP |
| IF_Peas1-260-F | ACCCTCACTAAAGGGAACAGACCCCAGACGCCAAG | for construction of pNPP243:P_NbEAS1-260:GFP |
| IF_Peas1-230-F | ACCCTCACTAAAGGGTAGGCTGCTGGTTGGCTGAC | for construction of pNPP243:P_NbEAS1-230:GFP |
| IF_Peas1-200-F | ACCCTCACTAAAGGGGTGTAAAGTCAAGTAAGGCA | for construction of pNPP243:P_NbEAS1-200:GFP |
| IF_Peas1-R | GCCCTTGCTCACCATTATATTGCTAATAAATTAAA | for construction of pNPP243:P_NbEAS1:GFP |
| IF_Peas4-400-F | ACCCTCACTAAAGGGGAAAGTTTTCAAATATCACT | for construction of pNPP243:P_NbEAS4-400:GFP |
| IF_Peas4-300-F | ACCCTCACTAAAGGGATTTTAGAGACGTAGGATGT | for construction of pNPP243:P_NbEAS4-300:GFP |
| IF_Peas4-260-F | ACCCTCACTAAAGGGATGTAAGAAATACTGCTCCT | for construction of pNPP243:P_NbEAS4-260:GFP |
| IF_Peas4-230-F | ACCCTCACTAAAGGGATCAATTGAGACGCCAAGAA | for construction of pNPP243:P_NbEAS4-230:GFP |
| IF_Peas4-200-F | ACCCTCACTAAAGGGAGACGCCAAGGAAGAAATAT | for construction of pNPP243:P_NbEAS4-200:GFP |
| IF_Peas4-230TT-F | ACCCTCACTAAAGGGATCAATTGATACTCCAAGAA | for construction of pNPP243:P_NbEAS4-230TT:GFP |
| IF_Peas4-R | GCCCTTGCTCACCATTGCTATAGAATTTATACTCT | for construction of pNPP243:P_NbEAS4:GFP |
| **Primers for construction of VIGS vectors** | | |
| NbACO2-BamHI-F | TACATAGGATCCTGGAGAGTTTCCCAGTGG | for construction of pTV00-ACO |
| NbACO2-SmaI-R | CAGAGCCCCGGGAAAAGTTGCTCTGCTAAT | for construction of pTV00-ACO |
| IF_pTV00_Nb36116-F | ACTAGTGGATCCCCCGATGAAGAAGCTGCAATT | for construction of pTV00-NbERF-IX-33 |
| IF_pTV00_Nb36116-R | GAATTCCTGCAGCCCTTAACTGACTATTAATTG | for construction of pTV00-NbERF-IX-33 |
| IF_pTV00_Nb25008-F | ACTAGTGGATCCCCCCAGGTCATGGGAAAGAAG | for construction of pTV00-NbERF-IX-16 |
| IF_pTV00_Nb25008-R | GAATTCCTGCAGCCCCTAGGTCCTGTCATAACC | for construction of pTV00-NbERF-IX-16 |
| **Primers for qRT-PCR** | | |
| RT-GUS-F | TCAAAAAACTCGACGGCCTG | for qRT-PCR of tGUS |
| RT-GUS-R | TTCGGTATAAAGACTTCGCG | for qRT-PCR of tGUS |
| RT-GFP-F | CAACTACAACAGCCACAACG | for qRT-PCR of GFP |
| RT-GFP-R | TCTTTGCTCAGGGCGGACTG | for qRT-PCR of GFP |
| **Primers for construction of protein expression vector** | | |
| IF_pMAL-36116-F | ATGGGCGGCCGCGATATGAATCATTCTATTTATAC | for construction of pMAL-NbERF-IX-33a |
| IF_pMAL-36116-R | GGATCCGTCGACGATTTAACTGACTATTAATTGAT | for construction of pMAL-NbERF-IX-33a |

Extension sequences for In-fusion reaction are in red letters.

Mismatches to introduce mutations are highlighted in blue letters.

Restriction sites used for the construction of vectors are underlined.

**Supplementary Table 3.** *Nicotiana benthamiana* genes categorized in cluster 2.

| Gene ID* | Expression (FPKM value) | | | | | | | | | Annotation (BlastX) |
|---|---|---|---|---|---|---|---|---|---|---|
| | 0 h (Control) | $H_2O$ | | | | 150 nM INF1 | | | | |
| | | 3 h | 6 h | 12 h | 24 h | 3 h | 6 h | 12 h | 24 h | |
| Niben101Scf01084g05013 | 89.89 | 186.30 | 368.21 | 600.92 | 472.93 | 247.88 | 508.01 | 1076.55 | 1194.83 | Glycine-rich protein 3-like [*N. tabacum*] |
| Niben101Ctg13380g00001 | 385.51 | 364.03 | 550.20 | 828.68 | 578.02 | 521.03 | 865.15 | 1149.75 | 1027.51 | Peptidyl-prolyl cis-trans isomerase [*N. sylvestris*] |
| Niben101Scf00428g14006 | 474.80 | 275.27 | 518.21 | 548.99 | 594.95 | 152.92 | 330.73 | 314.27 | 885.58 | Lysine-rich arabinogalactan protein 19-like [*N. sylvestris*] |
| Niben101Scf01084g03004 | 103.60 | 123.50 | 202.01 | 202.13 | 176.49 | 189.34 | 477.48 | 786.50 | 513.54 | Glycine-rich protein-like [*N. sylvestris*] |
| Niben101Scf00897g03002 | 87.50 | 56.10 | 142.58 | 136.14 | 117.91 | 360.45 | 591.60 | 416.94 | 511.91 | Glutathione S-transferase [*N. sylvestris*] |
| Niben101Scf25768g00021 | 55.89 | 53.63 | 105.20 | 87.05 | 73.63 | 131.45 | 244.79 | 340.24 | 401.41 | Protein RALF-like 27 [*N. tabacum*] |
| Niben101Scf00646g07010 | 48.09 | 142.16 | 249.09 | 289.19 | 104.00 | 484.94 | 741.33 | 461.93 | 389.30 | Uncharacterized protein LOC104227479 [*N. sylvestris*] |
| Niben101Scf02771g01007 | 99.13 | 133.84 | 156.75 | 122.69 | 119.63 | 228.76 | 295.33 | 232.14 | 377.09 | Heat shock cognate 70 kDa protein 2-like [*N. attenuata*] |
| Niben101Scf06590g00003 | 96.07 | 104.47 | 170.95 | 177.97 | 188.18 | 131.28 | 214.77 | 201.22 | 362.39 | L-ascorbate peroxidase 2, cytosolic [*N. tabacum*] |
| Niben101Scf10505g00007 | 69.73 | 44.34 | 79.57 | 106.93 | 54.91 | 93.57 | 271.58 | 225.90 | 342.59 | Protein disulfide-isomerase-like [*N. tomentosiformis*] |
| Niben101Scf13188g00012 | 46.14 | 49.60 | 118.23 | 204.73 | 165.49 | 75.52 | 135.37 | 267.48 | 305.54 | Abscisic stress-ripening protein 2 [*N. sylvestris*] |
| Niben101Scf01048g00008 | 188.71 | 83.96 | 128.06 | 288.16 | 247.34 | 83.87 | 129.00 | 421.04 | 292.60 | Glycine-rich domain-containing protein 1-like [*N. tabacum*] |
| Niben101Scf02886g01003 | 85.05 | 58.15 | 115.12 | 139.94 | 110.30 | 93.31 | 229.52 | 335.41 | 286.02 | Metallothionein-like protein type 2 [*N. attenuata*] * |
| Niben101Scf04364g01014 | 72.58 | 96.56 | 108.35 | 88.80 | 89.59 | 155.92 | 208.27 | 157.23 | 279.79 | Heat shock cognate 70 kDa protein 2-like [*N. attenuata*] |
| Niben101Scf03052g00011 | 107.10 | 82.41 | 101.34 | 196.07 | 165.30 | 85.40 | 160.23 | 319.47 | 278.98 | Glycine-rich protein-like [*N. sylvestris*] |
| Niben101Scf00447g02008 | 61.17 | 72.22 | 98.31 | 84.84 | 75.39 | 129.86 | 148.76 | 106.96 | 260.77 | Eukaryotic translation initiation factor 1A-like [*N. tomentosiformis*] |
| Niben101Scf02230g01003 | 59.86 | 55.93 | 74.09 | 76.65 | 64.64 | 80.33 | 105.32 | 142.47 | 258.76 | Leucine-rich repeat protein 1-like [*N. tomentosiformis*] |
| Niben101Scf01445g00016 | 81.09 | 35.38 | 89.26 | 168.54 | 45.05 | 78.21 | 178.18 | 307.02 | 255.86 | Cysteine proteinase 3 precursor [*N. tabacum*] |
| Niben101Scf04209g01002 | 162.33 | 124.74 | 258.54 | 233.77 | 75.35 | 247.61 | 662.72 | 811.22 | 248.52 | Uncharacterized protein LOC109213470 [*N. attenuata*] |
| Niben101Scf07459g00017 | 93.61 | 83.09 | 96.49 | 74.88 | 82.39 | 108.76 | 225.24 | 136.32 | 239.18 | Phosphomannomutase [*N. tabacum*] |
| Niben101Scf06195g00002 | 71.82 | 71.44 | 101.03 | 110.29 | 100.23 | 97.39 | 144.20 | 156.80 | 214.46 | L-ascorbate peroxidase 2, cytosolic [*N. tabacum*] |
| Niben101Scf05152g01001 | 46.47 | 23.08 | 65.72 | 80.39 | 35.08 | 21.61 | 83.74 | 120.30 | 209.42 | Cytochrome b561 and DOMON domain-containing protein [*N. sylvestris*] |
| Niben101Scf01365g05003 | 124.04 | 85.94 | 146.82 | 173.34 | 127.48 | 159.98 | 400.55 | 383.10 | 205.56 | Sucrose transporter 1-1 [*N. tabacum*] |
| Niben101Scf06726g00034 | 132.48 | 88.13 | 163.55 | 284.80 | 170.43 | 78.64 | 263.36 | 464.14 | 204.99 | BURP domain protein USPL1-like [*N. tomentosiformis*] |
| Niben101Scf00578g05007 | 20.47 | 22.40 | 33.65 | 28.81 | 28.82 | 35.65 | 182.73 | 113.98 | 203.12 | Calcium-binding protein PBP1-like [*N. sylvestris*] |
| Niben101Scf03823g01005 | 99.70 | 49.02 | 115.17 | 168.62 | 96.83 | 64.64 | 181.33 | 173.01 | 203.12 | Pectinesterase/pectinesterase inhibitor U1 [*N. attenuata*] |
| Niben101Scf05340g00009 | 65.13 | 66.31 | 87.84 | 104.41 | 70.41 | 82.35 | 118.49 | 142.08 | 201.43 | Uncharacterized protein LOC109212171 [*N. attenuata*] |
| Niben101Scf00983g02001 | 104.98 | 93.37 | 129.98 | 196.45 | 99.13 | 196.76 | 316.33 | 383.35 | 196.22 | Isocitrate dehydrogenase [NADP] [*N. tomentosiformis*] |
| Niben101Scf16258g02004 | 17.98 | 49.15 | 98.45 | 183.09 | 166.74 | 58.12 | 146.27 | 190.12 | 190.14 | Defensin J1-2 [*N. attenuata*] |
| Niben101Scf01032g00006 | 155.41 | 51.44 | 75.47 | 141.24 | 115.99 | 65.34 | 128.11 | 254.36 | 184.82 | Dihydroflavonol-4-reductase-like [*Capsicum annuum*] |
| Niben101Scf02319g12008 | 44.63 | 62.15 | 142.00 | 167.34 | 58.37 | 165.06 | 326.62 | 289.52 | 176.93 | BURP domain protein RD22-like [*N. tomentosiformis*] |
| Niben101Scf01008g03019 | 126.96 | 43.86 | 114.68 | 128.05 | 115.01 | 33.33 | 103.93 | 122.64 | 173.20 | Homeobox protein 2-like [*N. attenuata*] |
| Niben101Scf37733g00002 | 37.21 | 40.31 | 24.25 | 77.09 | 108.03 | 29.12 | 45.62 | 82.44 | 169.55 | 5'-adenylylsulfate reductase 1, chloroplastic-like [*N. tabacum*] |
| Niben101Scf04174g01004 | 42.69 | 51.49 | 81.87 | 63.01 | 35.57 | 156.94 | 201.60 | 176.25 | 160.52 | Glutathione S-transferase [*N. attenuata*] |
| Niben101Scf04140g00003 | 66.27 | 42.60 | 71.56 | 88.99 | 86.85 | 130.40 | 161.65 | 289.30 | 158.74 | Uncharacterized protein LOC104227479 [*N. sylvestris*] |
| Niben101Scf00345g01021 | 73.47 | 25.51 | 61.11 | 156.47 | 108.70 | 29.74 | 69.97 | 153.50 | 151.19 | Glutamine synthetase isoform X1 [N. sylvestris] |
| Niben101Scf00215g02009 | 58.49 | 44.93 | 64.45 | 79.24 | 52.86 | 74.39 | 95.53 | 125.93 | 143.89 | Leucine-rich repeat protein 1-like [*N. tomentosiformis*] |
| Niben101Scf02508g01012 | 33.94 | 44.83 | 63.71 | 55.96 | 47.11 | 74.77 | 95.84 | 82.31 | 130.22 | Small acidic protein 1 [*Solanum tuberosum*] |
| Niben101Scf05442g03015 | 48.76 | 32.51 | 42.00 | 50.91 | 54.13 | 31.66 | 69.33 | 137.25 | 127.96 | Phenylalanine ammonia-lyase [*N. tomentosiformis*] |
| Niben101Scf06890g01022 | 12.39 | 22.97 | 40.71 | 17.77 | 16.09 | 64.87 | 100.34 | 75.55 | 126.57 | Endoplasmin homolog [*N. attenuata*] |
| Niben101Scf18771g00012 | 23.30 | 39.70 | 48.06 | 31.87 | 54.34 | 44.17 | 132.75 | 59.77 | 124.88 | Heme-binding-like protein, chloroplastic [*N. sylvestris*] |
| Niben101Scf03460g04004 | 10.17 | 11.28 | 41.10 | 59.91 | 46.00 | 13.47 | 72.99 | 94.26 | 124.70 | Lignin-forming anionic peroxidase [*N. tomentosiformis*] |
| Niben101Scf02222g01001 | 28.17 | 30.90 | 50.12 | 47.74 | 32.53 | 47.75 | 75.04 | 76.39 | 118.67 | Uncharacterized protein LOC109212171 [*N. attenuata*] |
| Niben101Scf01025g02004 | 63.19 | 42.64 | 52.19 | 80.93 | 83.61 | 71.18 | 178.45 | 184.97 | 116.54 | Arginine decarboxylase-like [*N. attenuata*] |
| Niben101Scf01520g04005 | 121.86 | 41.25 | 68.33 | 121.99 | 84.75 | 47.41 | 97.89 | 183.82 | 116.02 | Stem-specific protein TSJT1 [*N. sylvestris*] |
| Niben101Scf03425g00005 | 67.70 | 35.73 | 46.07 | 73.70 | 55.07 | 45.82 | 92.22 | 172.05 | 115.06 | Sucrose transporter 1-1 [*N. tabacum*] |
| Niben101Scf08341g01001 | 48.05 | 37.80 | 52.26 | 76.23 | 68.23 | 48.29 | 77.60 | 74.02 | 113.28 | GTP-binding nuclear protein Ran-B1 [*N. tabacum*] |
| Niben101Scf07253g02012 | 26.46 | 37.12 | 58.03 | 55.75 | 33.51 | 56.42 | 94.60 | 72.70 | 110.58 | Microsomal glutathione S-transferase 3 [*N. tomentosiformis*] |
| Niben101Scf00837g11004 | 14.74 | 23.75 | 47.37 | 22.54 | 17.49 | 58.95 | 112.53 | 59.95 | 109.15 | Endoplasmin homolog [*N. attenuata*] |
| Niben101Scf00107g03016 | 9.55 | 17.72 | 33.46 | 21.41 | 9.64 | 44.75 | 73.71 | 58.79 | 106.56 | Kynurenine formamidase-like [*N. tabacum*] |

*Top 50 genes highly expressed in INF1-treated leaves 24 after the treatment.

**Supplementary Table 4.** *Nicotiana benthamiana* genes categorized in cluster 4.

| Gene ID* | Expression (FPKM value) | | | | | | | | | Annotation (BlastX) |
|---|---|---|---|---|---|---|---|---|---|---|
| | 0 h | $H_2O$ | | | | 150 nM INF1 | | | | |
| | (Control) | 3 h | 6 h | 12 h | 24 h | 3 h | 6 h | 12 h | 24 h | |
| Niben101Scf06525g03013 | 22.02 | 46.50 | 112.94 | 125.94 | 87.30 | 231.76 | 386.77 | 728.54 | 1805.14 | NbSAR8.2m gene product [*N. benthamiana*] |
| Niben101Scf02041g00002 | 4.05 | 35.41 | 33.97 | 29.04 | 16.17 | 65.65 | 68.15 | 173.15 | 1291.63 | Acidic chitinase PR-Q [*N. tabacum*] |
| Niben101Scf12789g00006 | 0.94 | 5.04 | 22.55 | 30.87 | 28.06 | 95.08 | 239.38 | 616.75 | 760.73 | Cysteine-rich repeat secretory protein 55-like [*N. tabacum*] |
| Niben101Scf10834g03005 | 23.16 | 32.45 | 61.18 | 40.45 | 25.38 | 89.88 | 179.66 | 210.68 | 409.89 | Calreticulin-3 NbCRT3a2 [*N. benthamiana*] |
| Niben101Scf01400g00014 | 3.65 | 3.18 | 2.65 | 2.29 | 3.61 | 2.98 | 2.00 | 6.50 | 399.79 | Pathogenesis-related protein R minor form [*N. sylvestris*] |
| Niben101Scf01719g08010 | 6.20 | 11.49 | 5.22 | 7.81 | 5.29 | 28.79 | 44.42 | 171.94 | 215.06 | PDR-type ACB transporter NbPDR2a [*N. benthamiana*] |
| Niben101Scf00414g07005 | 16.81 | 22.84 | 24.80 | 23.02 | 21.83 | 127.01 | 150.73 | 235.03 | 210.88 | Farnesyl pyrophosphate synthase NbFPPS1b [*N. benthamiana*] |
| Niben101Scf10055g07005 | 12.17 | 27.62 | 24.07 | 14.25 | 13.98 | 81.54 | 89.96 | 182.21 | 207.87 | NADP-dependent malic enzyme [*N. sylvestris*] |
| Niben101Scf06256g01001 | 18.72 | 18.53 | 33.23 | 22.98 | 17.01 | 48.16 | 73.74 | 123.27 | 195.57 | Calreticulin-3 NbCRT3a1 [*N. benthamiana*] |
| Niben101Scf01051g09003 | 9.76 | 7.27 | 7.28 | 12.04 | 12.31 | 8.28 | 10.37 | 49.70 | 180.33 | FAM10 family protein [*N. attenuata*] |
| Niben101Scf00213g01002 | 3.05 | 4.67 | 12.01 | 8.80 | 5.68 | 24.84 | 60.17 | 116.48 | 163.47 | G-type lectin S-receptor-like serine/threonine-protein kinase RLK1 [*N. sylvestris*] |
| Niben101Scf02636g05003 | 1.18 | 2.44 | 1.97 | 1.19 | 1.91 | 4.30 | 5.00 | 59.29 | 158.80 | Uncharacterized protein LOC107818746 [*N. tabacum*] |
| Niben101Scf03990g00010 | 4.50 | 3.89 | 9.46 | 16.69 | 5.66 | 13.11 | 67.90 | 114.79 | 154.67 | Peroxidase 21 [*N. sylvestris*] |
| Niben101Scf14996g00009 | 4.08 | 3.71 | 1.56 | 2.06 | 2.42 | 3.11 | 2.62 | 20.86 | 149.69 | Catalase isozyme 1 [*N. sylvestris*] |
| Niben101Scf08130g00014 | 1.06 | 2.46 | 2.37 | 1.17 | 1.05 | 6.13 | 10.12 | 26.12 | 141.70 | Glutathione S-transferase [*N. attenuata*] |
| Niben101Scf05283g00016 | 7.62 | 14.52 | 21.35 | 15.58 | 11.48 | 29.88 | 68.00 | 88.02 | 129.93 | Carboxylesterase 17 [*N. attenuata*] |
| Niben101Scf04941g01011 | 3.28 | 2.92 | 5.41 | 4.39 | 4.30 | 14.29 | 13.67 | 39.57 | 127.61 | LysM domain receptor-like kinase 4 [*N. sylvestris*] |
| Niben101Scf02111g16018 | 12.24 | 13.05 | 14.87 | 16.25 | 14.79 | 25.36 | 46.74 | 129.36 | 127.43 | ATP-citrate synthase beta chain protein 2 [*N. sylvestris*] |
| Niben101Scf08510g01006 | 1.46 | 7.71 | 6.36 | 5.90 | 4.40 | 48.53 | 78.68 | 201.24 | 120.12 | Heavy metal-associated isoprenylated plant protein 39-like [*N. attenuata*] |
| Niben101Scf02786g03004 | 0.77 | 8.29 | 10.83 | 3.79 | 1.24 | 48.18 | 46.50 | 64.95 | 117.52 | Uncharacterized protein LOC104218427 [*N. sylvestris*] |
| Niben101Scf03147g10015 | 8.85 | 3.73 | 8.71 | 2.24 | 3.77 | 4.36 | 13.27 | 33.80 | 105.04 | Glutathione S-transferase [*N. attenuata*] |
| Niben101Scf02195g01001 | 4.10 | 4.37 | 5.56 | 24.94 | 10.59 | 7.60 | 39.67 | 92.75 | 102.70 | Calcium-binding protein CML41 [*N. sylvestris*] |
| Niben101Scf03816g01001 | 4.48 | 5.00 | 7.21 | 7.66 | 5.62 | 26.67 | 38.31 | 76.94 | 101.82 | LRR receptor-like serine/threonine/tyrosine-protein kinase SOBIR1 [*N. tabacum*] |
| Niben101Scf02658g01008 | 1.66 | 11.01 | 13.00 | 5.17 | 3.63 | 37.10 | 60.20 | 48.98 | 97.14 | Dirigent protein 22-like [*N. tabacum*] |
| Niben101Scf08939g02004 | 0.74 | 1.50 | 0.94 | 1.30 | 1.43 | 1.55 | 1.56 | 18.77 | 96.15 | Hypothetical protein [*N. attenuata*] |
| Niben101Scf03015g03010 | 7.54 | 15.24 | 14.06 | 11.68 | 9.16 | 25.39 | 21.97 | 69.36 | 94.56 | 14-3-3-like protein E [*N. tabacum*] |
| Niben101Scf01111g01003 | 0.96 | 4.50 | 2.85 | 2.22 | 1.45 | 71.49 | 45.95 | 124.19 | 93.74 | 3-hydroxy-3-methylglutaryl-CoA synthase NbHMGR1a [*N. benthamiana*] |
| Niben101Ctg16217g00002 | 1.80 | 2.85 | 2.55 | 2.33 | 2.10 | 20.76 | 20.89 | 30.95 | 92.37 | Uncharacterized protein LOC109221780 [*N. attenuata*] |
| Niben101Scf05601g00010 | 1.08 | 3.82 | 6.41 | 6.89 | 1.96 | 31.14 | 47.02 | 72.24 | 91.65 | Sugar transport protein 13 [*N. attenuata*] |
| Niben101Scf09846g01010 | 2.57 | 3.76 | 8.89 | 4.40 | 9.05 | 34.70 | 54.04 | 48.75 | 87.76 | Quinone-oxidoreductase homolog, chloroplastic [*N. sylvestris*] |
| Niben101Scf01742g05006 | 4.10 | 3.94 | 4.22 | 3.78 | 4.54 | 14.83 | 22.37 | 67.11 | 81.92 | ATP-citrate synthase alpha chain protein 2 [*N. sylvestris*] |
| Niben101Scf05872g02002 | 6.41 | 5.26 | 11.69 | 13.60 | 20.42 | 4.51 | 14.18 | 14.84 | 80.91 | Dirigent protein 22-like [*N. attenuata*] |
| Niben101Scf08940g01001 | 0.71 | 8.52 | 6.24 | 1.79 | 0.87 | 36.15 | 37.28 | 68.50 | 74.60 | Lysine histidine transporter 1-like [*N. sylvestris*] |
| Niben101Scf10735g00016 | 7.96 | 5.89 | 8.69 | 22.11 | 18.46 | 7.16 | 17.58 | 54.43 | 72.74 | Pathogenesis-related protein STH-2-like [*N. attenuata*] |
| Niben101Scf10336g01001 | 1.76 | 1.44 | 2.60 | 3.38 | 3.48 | 5.63 | 63.99 | 54.61 | 72.12 | Metalloendoproteinase 1-like [*N. sylvestris*] |
| Niben101Scf05060g07008 | 0.06 | 3.24 | 2.89 | 1.86 | 0.67 | 6.92 | 108.07 | 31.76 | 68.41 | Calcium-binding protein CML46 [*N. tomentosiformis*] |
| Niben101Scf02171g00008 | 5.65 | 5.57 | 3.34 | 4.61 | 5.75 | 9.60 | 41.70 | 84.12 | 66.46 | Wound-inuduced protein kinase [*N. benthamiana*] |
| Niben101Scf02349g03002 | 2.20 | 3.18 | 3.12 | 2.72 | 1.56 | 4.71 | 6.87 | 16.06 | 66.32 | Suberization-associated anionic peroxidase-like [*N. tabacum*] |
| Niben101Scf02063g05001 | 1.54 | 1.53 | 0.75 | 2.79 | 1.77 | 1.55 | 6.87 | 27.40 | 63.16 | Ethylene-responsive transcription factor 1B-like [*N. attenuata*] |
| Niben101Scf01015g01002 | 0.35 | 0.40 | 0.67 | 0.50 | 0.78 | 0.50 | 1.01 | 9.87 | 62.80 | Wound-induced protein WIN1-like [*N. sylvestris*] |
| Niben101Scf02622g07001 | 3.08 | 2.07 | 1.90 | 2.08 | 3.09 | 6.29 | 15.38 | 40.40 | 61.44 | Glycosyltransferase family protein 64 protein C5-like [*N. attenuata*] |
| Niben101Scf08020g06001 | 0.81 | 5.66 | 4.06 | 2.04 | 1.41 | 14.14 | 58.08 | 32.91 | 59.25 | Harpin inducing protein [*N. tabacum*] |
| Niben101Scf04015g01004 | 0.28 | 0.73 | 0.39 | 1.19 | 1.71 | 2.86 | 3.34 | 24.19 | 59.10 | Phosphoglycerate mutase-like protein AT74H [*N. tomentosiformis*] |
| Niben101Scf02639g03015 | 3.70 | 7.54 | 4.41 | 5.55 | 5.38 | 13.45 | 44.18 | 42.02 | 58.78 | ALA-interacting subunit 1-like [*N. attenuata*] |
| Niben101Scf03114g02002 | 2.62 | 4.64 | 4.37 | 5.50 | 8.01 | 13.91 | 38.47 | 55.30 | 58.48 | Heavy metal-associated isoprenylated plant protein 39-like [*N. attenuata*] |
| Niben101Scf01574g09001 | 1.39 | 3.74 | 3.68 | 3.27 | 1.76 | 11.21 | 55.25 | 50.41 | 58.32 | Uncharacterized protein LOC104235457 [*N. sylvestris*] |
| Niben101Scf02072g02017 | 9.11 | 4.71 | 7.86 | 13.01 | 9.46 | 6.11 | 10.56 | 38.39 | 58.01 | Spermidine coumaroyl-CoA acyltransferase [*N. tomentosiformis*] |
| Niben101Scf05378g01017 | 6.52 | 2.59 | 10.84 | 9.79 | 4.88 | 17.18 | 21.20 | 35.65 | 57.95 | Uncharacterized protein LOC104240896 [*N. sylvestris*] |
| Niben101Scf00225g00003 | 2.09 | 8.32 | 3.16 | 3.27 | 4.82 | 12.81 | 5.29 | 32.70 | 57.38 | Flavanone 3-dioxygenase [*N. sylvestris*] |
| Niben101Scf00700g09001 | 3.44 | 13.07 | 12.96 | 7.05 | 4.15 | 23.71 | 28.36 | 27.08 | 56.92 | Caffeic acid 3-O-methyltransferase-like [*N. attenuata*] |

*Top 50 genes highly expressed in INF1-treated leaves 24 after the treatment.

**Supplementary Table 5.** *Nicotiana benthamiana* genes categorized in cluster 10.

| Gene ID* | Expression (FPKM value) | | | | | | | | | Annotation (BlastX) |
|---|---|---|---|---|---|---|---|---|---|---|
| | 0 h (Control) | $H_2O$ | | | | 150 nM INF1 | | | | |
| | | 3 h | 6 h | 12 h | 24 h | 3 h | 6 h | 12 h | 24 h | |
| Niben101Scf02543g01008 | 12.44 | 153.43 | 100.46 | 13.32 | 6.30 | 448.61 | 268.75 | 195.65 | 546.31 | 1-aminocyclopropane-1-carboxylate oxidase NbACO1a [*N. benthamiana*] |
| Niben101Scf13429g03011 | 0.19 | 42.41 | 128.05 | 7.14 | 1.40 | 412.72 | 625.15 | 232.46 | 110.86 | 14 kDa proline-rich protein DC2.15-like [*N. attenuata*] |
| Niben101Scf08039g01005 | 0.46 | 28.97 | 13.08 | 1.91 | 1.04 | 163.21 | 74.73 | 51.93 | 103.35 | 1-aminocyclopropane-1-carboxylate oxidase NbACO1b [*N. benthamiana*] |
| Niben101Scf07103g01015 | 30.63 | 23.82 | 177.94 | 33.97 | 0.00 | 23.38 | 80.96 | 22.93 | 87.86 | ATP synthase CF1 epsilon subunit [*N. tabacum*] |
| Niben101Scf00148g00004 | 0.94 | 21.83 | 14.87 | 1.00 | 1.10 | 41.15 | 82.72 | 62.45 | 75.85 | 21 kDa protein-like [*N. tabacum*] |
| Niben101Scf05279g05006 | 0.34 | 20.97 | 7.76 | 1.01 | 0.41 | 39.81 | 41.60 | 43.19 | 37.01 | Uncharacterized protein LOC104231083 [*N. sylvestris*] |
| Niben101Scf01236g05023 | 15.21 | 23.39 | 22.59 | 20.52 | 21.26 | 24.72 | 0.00 | 30.89 | 31.88 | Hypothetical protein Lal_00000388 [*Lupinus albus*] |
| Niben101Scf01640g03006 | 0.23 | 58.93 | 7.90 | 0.72 | 0.48 | 79.67 | 9.54 | 10.33 | 28.87 | hypothetical protein A4A49_07063 [*N. attenuata*] |
| Niben101Scf07829g01008 | 0.51 | 20.59 | 2.44 | 0.57 | 0.17 | 51.84 | 48.36 | 29.95 | 27.43 | uncharacterized protein LOC104247702 [*N. sylvestris*] |
| Niben101Ctg15035g00004 | 0.04 | 4.70 | 4.56 | 0.41 | 0.28 | 53.87 | 26.39 | 36.10 | 25.73 | CASP-like protein PIMP1 [*N. attenuata*] |
| Niben101Scf03861g00008 | 0.05 | 1.92 | 0.20 | 0.21 | 0.11 | 6.95 | 12.04 | 10.89 | 16.82 | Peroxidase 4 [*Capsicum chinense*] |
| Niben101Scf05326g00010 | 0.19 | 3.51 | 3.03 | 0.48 | 0.35 | 8.02 | 9.15 | 21.07 | 14.71 | Lipoxygenase homology domain-containing protein 1-like [*N. tabacum*] |
| Niben101Scf00109g07005 | 0.00 | 1.36 | 0.39 | 0.03 | 0.04 | 1.06 | 4.46 | 9.35 | 12.93 | Calcium-binding protein CML19 [*N. attenuata*] |
| Niben101Scf00755g00004 | 8.66 | 6.30 | 9.69 | 4.36 | 0.00 | 5.35 | 12.22 | 4.48 | 12.65 | Unknown protein |
| Niben101Scf03647g03023 | 0.21 | 0.22 | 0.00 | 0.10 | 0.13 | 0.12 | 0.07 | 0.40 | 11.33 | Uncharacterized protein LOC104118395 [*N. tomentosiformis*] |
| Niben101Scf00390g03001 | 0.13 | 2.72 | 0.57 | 0.23 | 0.07 | 3.06 | 0.70 | 3.49 | 9.79 | Proline dehydrogenase 2, mitochondrial-like [*N. tabacum*] |
| Niben101Scf21484g01002 | 4.54 | 1.21 | 4.23 | 6.74 | 3.96 | 0.00 | 6.62 | 1.83 | 9.15 | Uncharacterized protein LOC104099480 [*N. tomentosiformis*] |
| Niben101Scf00050g01002 | 1.76 | 5.55 | 1.39 | 0.38 | 2.07 | 2.98 | 2.33 | 2.34 | 7.85 | Phylloplanin-like [*N. tabacum*] |
| Niben101Scf05118g11009 | 0.00 | 0.94 | 0.56 | 0.01 | 0.03 | 3.21 | 1.29 | 2.51 | 6.08 | E3 ubiquitin-protein ligase Hakai-like [*N. attenuata*] |
| Niben101Scf02613g05004 | 0.06 | 1.32 | 0.29 | 0.00 | 0.00 | 2.38 | 0.97 | 0.87 | 6.08 | EG45-like domain containing protein [*N. tabacum*] |
| Niben101Scf06578g00021 | 4.86 | 1.54 | 2.51 | 2.43 | 2.81 | 0.22 | 2.03 | 2.90 | 5.84 | Calcium-binding protein PBP1-like [*N. sylvestris*] |
| Niben101Scf07751g00003 | 0.19 | 2.65 | 0.84 | 0.28 | 0.03 | 7.94 | 6.27 | 6.11 | 5.73 | AT-hook motif nuclear-localized protein 17-like [*N. attenuata*] |
| Niben101Scf09078g00001 | 1.65 | 0.81 | 0.60 | 0.50 | 0.71 | 0.03 | 0.48 | 1.15 | 5.07 | Uncharacterized protein LOC108943634 [*N. tomentosiformis*] |
| Niben101Scf01376g04034 | 0.09 | 6.37 | 1.48 | 0.11 | 0.02 | 9.41 | 4.70 | 3.45 | 5.02 | Late blight resistance protein homolog R1B-16 isoform X2 [*N. tabacum*] |
| Niben101Scf02085g12016 | 0.01 | 2.20 | 0.25 | 0.11 | 0.01 | 13.05 | 2.98 | 3.36 | 4.48 | L-type lectin-domain containing receptor kinase S.4-like [*N.* attenuata] |
| Niben101Scf01196g00008 | 1.67 | 1.58 | 0.00 | 3.28 | 2.74 | 0.31 | 0.74 | 0.71 | 4.22 | Unknown protein |
| Niben101Scf00372g02011 | 0.01 | 1.36 | 0.07 | 0.09 | 0.03 | 7.28 | 7.62 | 5.52 | 4.11 | uncharacterized protein DDB_G0271670-like [*N. attenuata*] |
| Niben101Ctg10923g00001 | 0.24 | 0.31 | 0.21 | 0.29 | 0.24 | 0.19 | 0.02 | 0.81 | 3.87 | vacuolar-sorting receptor 6-like [*N. tomentosiformis*] |
| Niben101Scf05434g00011 | 3.25 | 3.59 | 1.06 | 1.07 | 0.00 | 14.44 | 4.78 | 4.88 | 3.79 | Sulfate transporter 3.3 [*N. attenuata*] |
| Niben101Scf01451g05015 | 2.19 | 1.05 | 0.00 | 2.29 | 1.35 | 0.52 | 1.35 | 2.31 | 3.74 | oligoribonuclease-like [*N. attenuata*] |
| Niben101Scf02191g03003 | 0.15 | 0.33 | 0.00 | 0.32 | 2.33 | 0.13 | 0.12 | 0.24 | 3.74 | hypothetical protein H5410_023541 [*Solanum commersonii*] |
| Niben101Scf08718g00004 | 6.76 | 0.35 | 1.23 | 0.93 | 0.08 | 1.51 | 2.96 | 3.62 | 2.80 | Unknown protein |
| Niben101Scf01212g01017 | 0.75 | 1.81 | 2.71 | 1.34 | 1.17 | 0.17 | 1.56 | 1.84 | 2.76 | photosystem II protein D1/D2 superfamily [*Helianthus annuus*] |
| Niben101Scf00484g03003 | 1.22 | 0.76 | 1.21 | 1.43 | 1.33 | 1.46 | 0.00 | 1.76 | 2.70 | serine/arginine-rich SC35-like splicing factor SCL33 isoform X2 [*N. attenuata*] |
| Niben101Scf02907g06004 | 0.02 | 1.99 | 0.80 | 0.00 | 0.95 | 3.76 | 2.31 | 0.47 | 2.43 | 8-hydroxygeraniol dehydrogenase-like [*N. tomentosiformis*] |
| Niben101Scf03213g01009 | 1.09 | 0.52 | 0.05 | 0.58 | 0.60 | 0.36 | 0.39 | 0.97 | 1.87 | hypothetical protein EJD97_005407 [*S. chilense*] |
| Niben101Scf02060g00010 | 0.15 | 0.13 | 0.24 | 0.21 | 0.00 | 0.30 | 0.23 | 0.70 | 1.78 | MADS-box transcription factor 23-like [*N. attenuata*] |
| Niben101Scf08465g03014 | 0.34 | 0.44 | 0.00 | 0.17 | 0.30 | 0.28 | 0.08 | 0.08 | 1.72 | GDSL esterase/lipase At5g37690 [*N. sylvestris*] |
| Niben101Scf10492g00004 | 0.08 | 0.08 | 0.06 | 0.10 | 0.13 | 0.00 | 0.08 | 0.46 | 1.69 | Glycosyltransferase At5g25310 [*N. attenuata*] |
| Niben101Scf09710g00026 | 0.65 | 0.60 | 1.31 | 2.00 | 0.78 | 0.58 | 0.11 | 1.01 | 1.57 | Hypothetical protein NitaMp145 [*N. tabacum*] |
| Niben101Scf02869g18005 | 0.12 | 0.24 | 0.19 | 0.00 | 0.00 | 0.27 | 0.08 | 0.37 | 1.54 | Elicitor-responsive protein 1-like [*N. sylvestris*] |
| Niben101Scf00428g16011 | 0.47 | 0.42 | 0.41 | 0.16 | 0.63 | 1.47 | 1.08 | 0.00 | 1.51 | V-type proton ATPase subunit d2 [*N. sylvestris*] |
| Niben101Scf01517g08025 | 0.40 | 0.83 | 0.70 | 1.82 | 0.72 | 0.00 | 1.72 | 1.38 | 1.44 | Uncharacterized protein LOC104232423 [*N. sylvestris*] |
| Niben101Scf07792g00002 | 0.28 | 0.49 | 1.37 | 0.42 | 0.52 | 0.00 | 1.72 | 0.33 | 1.41 | Vicilin-like seed storage protein [*N. attenuata*] |
| Niben101Scf04731g11004 | 0.02 | 0.25 | 0.02 | 0.00 | 0.02 | 0.06 | 0.03 | 0.06 | 1.26 | Citrate-binding protein-like [*N. tabacum*] |
| Niben101Scf03886g04003 | 0.92 | 0.48 | 0.45 | 1.18 | 0.54 | 0.04 | 0.56 | 1.03 | 1.15 | HVA22-like protein e [*N. tabacum*] |
| Niben101Scf02085g12014 | 0.06 | 13.99 | 0.77 | 0.05 | 0.01 | 29.16 | 1.91 | 0.96 | 1.07 | E3 ubiquitin-protein ligase Praja-2-like [*N. sylvestris*] |
| Niben101Scf06517g00020 | 0.30 | 0.25 | 0.00 | 0.15 | 0.13 | 0.25 | 0.11 | 0.37 | 1.00 | MADS-box transcription factor 23-like isoform X1 [*N. attenuata*] |
| Niben101Scf01111g03001 | 0.03 | 0.03 | 0.05 | 0.00 | 0.00 | 0.03 | 0.04 | 0.15 | 0.99 | MATE efflux family protein 5-like [*N. sylvestris*] |
| Niben101Scf04981g00001 | 0.02 | 0.22 | 0.08 | 0.00 | 0.03 | 0.23 | 0.11 | 1.20 | 0.96 | Glutathione transferase GST 23-like [*N. sylvestris*] |
| Niben101Scf03422g04036 | 0.02 | 1.05 | 0.07 | 0.00 | 0.09 | 5.04 | 0.98 | 2.11 | 0.93 | Plant cadmium resistance 8-like [*N. tomentosiformis*] |

*Top 50 genes highly expressed in INF1-treated leaves 24 after the treatment.

**Supplementary Table 6.** *Nicotiana benthamiana* genes categorized in cluster 14.

| Gene ID* | Expression (FPKM value) | | | | | | | | | Annotation (BlastX) |
|---|---|---|---|---|---|---|---|---|---|---|
| | 0 h (Control) | $H_2O$ | | | | 150 nM INF1 | | | | |
| | | 3 h | 6 h | 12 h | 24 h | 3 h | 6 h | 12 h | 24 h | |
| Niben101Scf08921g02024 | 0.47 | 0.45 | 2.92 | 3.21 | 1.98 | 3.83 | 11.26 | 181.07 | 963.96 | Zingipain-2-like [*N. attenuata*] |
| Niben101Scf03385g02011 | 1.98 | 4.01 | 4.21 | 2.75 | 2.62 | 39.14 | 52.84 | 211.72 | 690.42 | Uncharacterized protein LOC104232799 [*N. sylvestris*] |
| Niben101Ctg13736g00004 | 0.38 | 0.22 | 0.40 | 0.70 | 5.90 | 2.86 | 3.69 | 86.46 | 672.36 | Glucan endo-1,3-beta-glucosidase, acidic isoform GI9-like [*N. sylvestris*] |
| Niben101Scf01001g00003 | 0.27 | 0.16 | 0.27 | 0.71 | 5.81 | 2.74 | 3.55 | 76.80 | 626.78 | Glucan endo-1,3-beta-glucosidase, acidic isoform GI9-like [*N. sylvestris*] |
| Niben101Scf09740g00003 | 0.18 | 0.06 | 0.27 | 2.17 | 5.74 | 0.13 | 0.43 | 19.50 | 600.88 | Uncharacterized protein LOC107783307 [*N. tabacum*] |
| Niben101Scf01001g00005 | 0.26 | 0.12 | 0.25 | 0.74 | 5.36 | 2.67 | 3.21 | 73.94 | 570.31 | Glucan endo-1,3-beta-glucosidase, acidic isoform gi9 [*N. attenuata*] |
| Niben101Scf00107g03008 | 0.25 | 0.03 | 0.14 | 0.50 | 5.27 | 0.16 | 0.55 | 28.48 | 569.39 | Pathogenesis-related protein 1B [*N. tabacum*] |
| Niben101Scf03930g00018 | 0.97 | 26.93 | 14.27 | 3.22 | 1.74 | 214.84 | 83.54 | 192.51 | 566.23 | Cucumber peeling cupredoxin-like [*N. sylvestris*] |
| Niben101Scf02407g03010 | 1.56 | 5.27 | 5.21 | 4.44 | 3.72 | 34.22 | 48.64 | 201.52 | 560.15 | Uncharacterized protein LOC109240683 [*N. attenuata*] |
| Niben101Scf01001g00004 | 0.19 | 0.14 | 0.24 | 0.70 | 4.98 | 2.44 | 3.02 | 68.19 | 530.15 | Glucan endo-1,3-beta-glucosidase, acidic isoform GI9-like [*N. sylvestris*] |
| Niben101Scf00640g04023 | 0.00 | 0.73 | 0.73 | 0.79 | 0.35 | 9.14 | 17.16 | 86.86 | 434.56 | Bifunctional epoxide hydrolase 2-like [*N. tomentosiformis*] |
| Niben101Scf01084g03003 | 0.08 | 0.71 | 1.94 | 0.22 | 0.12 | 13.73 | 31.94 | 96.53 | 374.31 | NbSAR8.2d gene product [*N. benthamiana*] |
| Niben101Scf01934g02004 | 0.04 | 0.09 | 0.05 | 0.09 | 0.14 | 0.60 | 4.93 | 131.70 | 334.61 | Glucan endo-1,3-beta-glucosidase, basic vacuolar isoform GGIB50 [*N. sylvestris*] |
| Niben101Scf02877g02005 | 0.12 | 0.06 | 0.28 | 0.58 | 1.03 | 2.96 | 26.29 | 124.35 | 324.87 | Strictosidine synthase 1-like [*N. sylvestris*] |
| Niben101Scf02203g05002 | 1.33 | 6.71 | 4.77 | 2.78 | 1.69 | 82.45 | 94.85 | 222.74 | 295.51 | 3-hydroxy-3-methylglutaryl-coenzyme A reductase NbHMGR2 [*N. benthamiana*] |
| Niben101Scf02819g00005 | 0.70 | 4.95 | 7.08 | 2.14 | 1.49 | 58.05 | 53.59 | 109.74 | 280.57 | Cupredoxin-like [*N. tomentosiformis*] |
| Niben101Scf04869g03002 | 0.34 | 0.21 | 0.51 | 0.68 | 2.07 | 3.66 | 6.26 | 56.88 | 276.55 | Glucan endo-1,3-beta-glucosidase, acidic isoform GL161 [*N. sylvestris*] |
| Niben101Scf02918g00003 | 0.31 | 0.53 | 0.97 | 0.74 | 0.55 | 11.06 | 22.61 | 202.69 | 275.20 | 1-aminocyclopropane-1-carboxylate oxidase NbACO4 [*N. benthamiana*] |
| Niben101Scf35444g00004 | 0.56 | 5.66 | 5.06 | 3.01 | 2.47 | 19.32 | 27.38 | 81.16 | 263.68 | Glutathione S-transferase [*N. sylvestris*] |
| Niben101Scf10986g00001 | 0.73 | 1.38 | 2.20 | 3.17 | 4.16 | 11.74 | 21.50 | 142.36 | 219.62 | Cysteine-rich repeat secretory protein 55-like [*N. tabacum*] |
| Niben101Scf03993g05005 | 0.09 | 6.12 | 4.35 | 0.57 | 0.25 | 305.27 | 189.22 | 239.25 | 211.45 | 5-*epi*-aristolochene synthase NbEAS4 [*N. benthamiana*] |
| Niben101Scf04053g02006 | 0.03 | 0.00 | 0.00 | 0.04 | 0.08 | 0.01 | 0.24 | 21.12 | 206.72 | Basic form of pathogenesis-related protein 1-like [*N. sylvestris*] |
| Niben101Scf10488g01001 | 0.97 | 2.24 | 1.70 | 0.69 | 0.91 | 6.16 | 8.04 | 52.83 | 204.46 | Uncharacterized protein LOC104118395 [*N. tomentosiformis*] |
| Niben101Scf00712g02011 | 0.04 | 0.38 | 0.17 | 0.04 | 0.08 | 39.41 | 41.88 | 121.75 | 192.66 | 5-*epi*-aristolochene synthase NbEAS7 [*N. benthamiana*] |
| Niben101Scf07201g01009 | 0.85 | 8.24 | 3.64 | 2.95 | 0.89 | 31.77 | 32.00 | 104.23 | 182.99 | AAA-ATPase-like [*N. attenuata*] |
| Niben101Scf00577g09001 | 2.27 | 1.16 | 1.55 | 4.40 | 1.19 | 2.19 | 8.30 | 216.07 | 158.08 | Hypothetical protein A4A49_37911 [*N. attenuata*] |
| Niben101Scf01534g02014 | 0.00 | 0.21 | 0.75 | 0.24 | 0.06 | 2.76 | 7.63 | 69.46 | 150.34 | Zinc finger protein ZIC 2-like [*N. attenuata*] |
| Niben101Scf00821g16001 | 0.02 | 0.32 | 0.19 | 0.07 | 0.13 | 3.11 | 6.91 | 45.36 | 147.17 | Somatic embryogenesis receptor kinase 2-like [*N. tabacum*] |
| Niben101Scf07491g00003 | 0.15 | 0.10 | 0.05 | 0.07 | 0.30 | 0.31 | 1.52 | 33.80 | 144.26 | Endochitinase A [*N. tomentosiformis*] |
| Niben101Scf00313g07005 | 1.68 | 1.04 | 1.01 | 2.02 | 0.81 | 1.18 | 5.84 | 41.21 | 133.39 | Benzyl alcohol O-benzoyltransferase [*N. attenuata*] |
| Niben101Scf08057g00006 | 0.20 | 1.07 | 0.67 | 1.02 | 1.03 | 12.03 | 17.16 | 78.40 | 127.41 | Plant cadomiun resistance 2-like [*N. tabacum*] |
| Niben101Scf03374g06002 | 0.08 | 0.00 | 0.00 | 0.00 | 0.37 | 0.21 | 0.20 | 9.95 | 116.52 | Lipid transfer-like protein VAS [*N. sylvestris*] |
| Niben101Scf05404g09001 | 0.04 | 0.50 | 0.50 | 0.30 | 0.25 | 2.83 | 4.46 | 21.65 | 115.96 | Glutathione S-transferase [*N. sylvestris*] |
| Niben101Scf09044g01012 | 0.52 | 0.07 | 0.22 | 0.39 | 0.60 | 0.49 | 1.27 | 20.22 | 113.93 | Osmotin [*N. sylvestris*] |
| Niben101Scf07767g02011 | 0.77 | 0.67 | 2.90 | 1.64 | 1.60 | 2.09 | 14.79 | 80.84 | 112.55 | Cytochrome P450 71A1-like [*N. sylvestris*] |
| Niben101Scf01999g07002 | 0.07 | 0.10 | 0.11 | 0.58 | 0.74 | 0.44 | 1.68 | 24.87 | 112.37 | Pathogenesis-related protein 1C-like [*N. sylvestris*] |
| Niben101Scf04787g02002 | 0.01 | 0.03 | 0.02 | 0.00 | 0.00 | 1.58 | 10.50 | 84.68 | 111.26 | 9-divinyl ether synthase [*N. tabacum*] |
| Niben101Scf00354g01012 | 0.02 | 0.00 | 0.00 | 0.00 | 0.02 | 0.05 | 2.27 | 34.86 | 109.31 | deacetylvindoline O-acetyltransferase-like [*N. sylvestris*] |
| Niben101Scf00700g00005 | 0.01 | 1.27 | 0.24 | 0.06 | 0.03 | 41.47 | 29.76 | 73.28 | 105.38 | 5-*epi*-aristolochene synthase NbEAS6 [*N. benthamiana*] |
| Niben101Scf03993g06006 | 0.06 | 4.49 | 1.47 | 0.35 | 0.13 | 220.63 | 117.72 | 153.02 | 102.64 | 5-*epi*-aristolochene synthase NbEAS3 [*N. benthamiana*] |
| Niben101Scf12045g06025 | 0.54 | 0.84 | 1.81 | 1.67 | 0.95 | 3.03 | 4.61 | 7.59 | 101.52 | Pathogenesis-related protein PR-4B [*N. tomentosiformis*] |
| Niben101Scf03096g01022 | 0.00 | 0.43 | 1.11 | 0.07 | 0.07 | 14.71 | 29.48 | 14.21 | 87.06 | Cysteine-rich repeat secretory protein 38-like [*N. tabacum*] |
| Niben101Scf01942g04001 | 0.03 | 0.12 | 0.02 | 0.12 | 0.07 | 0.77 | 6.02 | 79.56 | 85.73 | WRKY transcription factor 51 [*N. sylvestris*] |
| Niben101Scf01249g06013 | 0.01 | 1.31 | 0.35 | 0.17 | 0.04 | 5.95 | 7.20 | 59.02 | 85.59 | Mitochondrial phosphate carrier protein 3 [*N. attenuata*] |
| Niben101Scf04944g05002 | 0.70 | 0.50 | 0.25 | 0.19 | 0.46 | 0.47 | 3.73 | 80.35 | 85.01 | WRKY transcription factor 40 [*N. sylvestris*] |
| Niben101Scf01084g01009 | 0.17 | 0.41 | 0.19 | 0.45 | 0.26 | 3.47 | 9.87 | 34.06 | 84.57 | NbSAR8.2b gene product [*N. benthamiana*] |
| Niben101Scf16114g01003 | 0.43 | 0.07 | 0.17 | 0.30 | 0.24 | 0.99 | 2.21 | 24.79 | 84.15 | GDSL esterase/lipase 5-like [*N. tabacum*] |
| Niben101Scf06583g03008 | 0.47 | 1.40 | 0.53 | 0.54 | 0.24 | 2.75 | 5.78 | 63.57 | 83.64 | PDR-type ACB transporter NbPDR2b [*N. benthamiana*] |
| Niben101Scf02410g00002 | 0.46 | 1.16 | 0.94 | 0.79 | 0.79 | 3.16 | 6.39 | 32.35 | 83.25 | basic endochitinase [*N. attenuata*] |
| Niben101Scf06424g01007 | 0.07 | 0.28 | 0.26 | 0.06 | 0.19 | 0.49 | 0.47 | 5.93 | 80.44 | Kunitz trypsin inhibitor 2-like [*N. attenuata*] |

*Top 50 genes highly expressed in INF1-treated leaves 24 after the treatment.

**Supplementary Table 7.** *Nicotiana benthamiana* genes categorized in cluster 16.

| Gene ID* | Expression (FPKM value) | | | | | | | | | Annotation (BlastX) |
|---|---|---|---|---|---|---|---|---|---|---|
| | 0 h (Control) | $H_2O$ | | | | 150 nM INF1 | | | | |
| | | 3 h | 6 h | 12 h | 24 h | 3 h | 6 h | 12 h | 24 h | |
| Niben101Scf01239g02002 | 15.48 | 24.48 | 37.27 | 20.16 | 12.63 | 266.74 | 275.50 | 301.44 | 221.61 | Caffeic acid 3-o-methyltransferase [*N. attenuata*] |
| Niben101Scf14679g00002 | 7.86 | 74.42 | 62.98 | 86.22 | 33.42 | 255.22 | 210.37 | 202.67 | 178.27 | Secoisolariciresinol dehydrogenase-like [*N. attenuata*] |
| Niben101Scf07242g07006 | 11.32 | 42.33 | 23.98 | 24.63 | 11.52 | 119.78 | 105.29 | 174.13 | 160.51 | Isoflavone 2'-hydroxylase-like [*N. sylvestris*] |
| Niben101Scf08127g08009 | 14.25 | 43.11 | 49.98 | 22.21 | 17.49 | 144.22 | 129.21 | 120.31 | 159.52 | 6-phosphogluconate dehydrogenase, decarboxylating 1 [*N. sylvestris*] |
| Niben101Scf02171g00007 | 0.44 | 8.28 | 8.73 | 2.60 | 2.79 | 119.65 | 104.75 | 112.09 | 156.17 | Endochitinase PR4-like [*N. sylvestris*] |
| Niben101Scf00225g00008 | 6.67 | 10.53 | 20.79 | 8.99 | 4.64 | 173.25 | 211.25 | 242.64 | 142.15 | Caffeic acid 3-o-methyltransferase [*N. attenuata*] |
| Niben101Scf02030g04003 | 14.33 | 43.93 | 84.07 | 15.93 | 15.11 | 131.06 | 171.13 | 71.67 | 128.26 | Luminal-binding protein 4-like [*N. sylvestris*] |
| Niben101Scf01100g01006 | 11.62 | 19.92 | 14.66 | 13.42 | 11.42 | 92.14 | 68.73 | 128.61 | 125.41 | acetyl-CoA acetyltransferase NbACAT1b [*N. benthamiana*] |
| Niben101Scf04083g04040 | 13.65 | 64.13 | 71.19 | 15.14 | 13.88 | 195.78 | 122.63 | 92.00 | 122.28 | Luminal-binding protein 5 precursor [*N. tabacum*] |
| Niben101Scf10688g01013 | 3.56 | 20.07 | 7.06 | 8.25 | 4.25 | 57.26 | 54.07 | 141.83 | 120.20 | Blue copper protein-like [*N. attenuata*] |
| Niben101Scf08590g00005 | 19.87 | 56.62 | 85.81 | 21.01 | 17.67 | 179.18 | 139.38 | 89.06 | 114.01 | Luminal-binding protein 5 precursor [*N. tabacum*] |
| Niben101Scf08196g01001 | 8.12 | 91.99 | 88.23 | 29.97 | 6.15 | 394.72 | 363.31 | 83.10 | 112.11 | Glutathione S-transferase parA [*N. sylvestris*] |
| Niben101Scf04563g01016 | 6.94 | 39.80 | 8.93 | 7.63 | 8.36 | 208.40 | 31.85 | 65.94 | 102.61 | Methyltransferase DDB_G0268948 [*N. tabacum*] |
| Niben101Scf00031g02001 | 4.49 | 31.11 | 17.22 | 7.03 | 5.77 | 199.46 | 169.58 | 139.04 | 101.99 | Epidermis-specific secreted glycoprotein EP1-like precursor [*N. tabacum*] |
| Niben101Scf22099g00006 | 3.42 | 23.84 | 10.41 | 4.18 | 5.02 | 65.97 | 70.73 | 90.85 | 91.44 | Cytochrome b561 and DOMON domain-containing protein [*N. attenuata*] |
| Niben101Scf14394g01020 | 10.40 | 40.39 | 55.62 | 14.56 | 13.58 | 125.73 | 98.74 | 41.34 | 90.46 | Luminal-binding protein [*N. sylvestris*] |
| Niben101Scf01729g01015 | 2.16 | 6.73 | 5.01 | 3.72 | 2.55 | 92.02 | 62.07 | 120.93 | 81.58 | 3-hydroxy-3-methylglutaryl-CoA synthase NbHMGS1b [*N. benthamiana*] |
| Niben101Scf00779g06009 | 9.06 | 70.58 | 22.10 | 13.65 | 8.46 | 193.65 | 60.42 | 68.57 | 81.37 | 12-oxophytodienoate reductase 2-like [*N. attenuata*] |
| Niben101Scf00372g05012 | 7.71 | 32.83 | 32.90 | 21.03 | 4.27 | 134.56 | 169.22 | 114.88 | 79.78 | Hypothetical protein A4A49_08095 [*N. attenuata*] |
| Niben101Scf14069g00002 | 5.78 | 83.18 | 44.21 | 13.01 | 3.72 | 126.80 | 70.81 | 53.78 | 77.97 | Cysteine-rich and transmembrane domain-containing protein A [*N. attenuata*] |
| Niben101Scf01719g08001 | 8.86 | 17.34 | 14.04 | 13.43 | 6.68 | 155.73 | 97.70 | 173.31 | 74.20 | PDR-type ACB transporter NbPDR1a [*N. benthamiana*] |
| Niben101Scf03438g02001 | 0.03 | 24.02 | 4.85 | 5.91 | 1.59 | 29.70 | 19.85 | 25.66 | 73.02 | Heat stress transcription factor B-3-like [*N. tabacum*] |
| Niben101Scf01596g12004 | 8.90 | 41.68 | 49.57 | 11.28 | 9.45 | 110.67 | 83.66 | 45.19 | 71.86 | Calnexin homolog 1-like [*N. sylvestris*] |
| Niben101Scf02907g06031 | 4.83 | 48.97 | 17.50 | 6.15 | 5.49 | 253.91 | 82.05 | 85.73 | 69.25 | Mannitol dehydrogenase [*N. tomentosiformis*] |
| Niben101Scf00173g05015 | 9.28 | 9.69 | 9.67 | 9.71 | 9.85 | 48.83 | 48.67 | 76.54 | 69.04 | Mevalonate-5-pyrophosphate decarboxylase NbMVD1b [*N. benthamiana*] |
| Niben101Scf02353g06039 | 0.84 | 14.62 | 12.57 | 6.24 | 4.57 | 77.78 | 70.00 | 57.84 | 65.98 | Kunitz trypsin inhibitor 2-like [*N. attenuata*] |
| Niben101Scf01241g02004 | 1.19 | 37.27 | 29.24 | 11.71 | 5.15 | 182.55 | 85.53 | 42.96 | 64.76 | Glutamate decarboxylase 4 [*N. tabacum*] |
| Niben101Scf07395g00029 | 10.75 | 35.72 | 36.11 | 20.32 | 12.80 | 150.39 | 71.41 | 62.27 | 63.51 | NRT1/PTR family 2.11-like [*N. tabacum*] |
| Niben101Scf04003g08001 | 5.54 | 11.79 | 6.94 | 5.34 | 5.42 | 57.96 | 98.26 | 131.20 | 62.16 | Caffeoyl-CoA O-methyltransferase 4 [*N. tabacum*] |
| Niben101Scf12738g00010 | 4.30 | 37.50 | 28.66 | 34.61 | 10.36 | 114.92 | 114.47 | 35.09 | 59.37 | Ferredoxin, root R-B2-like [*N. tomentosiformis*] |
| Niben101Scf09203g01012 | 6.75 | 10.90 | 31.05 | 6.78 | 6.87 | 53.81 | 70.40 | 38.49 | 57.78 | GTP-binding protein SAR2 [*N. tomentosiformis*] |
| Niben101Scf00031g01005 | 3.89 | 48.25 | 22.22 | 5.33 | 5.43 | 196.70 | 145.56 | 104.91 | 52.58 | Epidermis-specific secreted glycoprotein EP1-like [*N. tabacum*] |
| Niben101Scf04122g05014 | 6.07 | 17.43 | 4.74 | 2.65 | 2.68 | 34.44 | 17.00 | 43.74 | 52.09 | Sugar carrier protein C-like [*N. sylvestris*] |
| Niben101Scf00637g04001 | 10.19 | 67.10 | 42.47 | 32.03 | 13.12 | 260.39 | 219.03 | 112.55 | 51.33 | Polygalacturonase inhibitor-like [*N. attenuata*] |
| Niben101Scf03115g02008 | 5.17 | 26.45 | 31.24 | 5.74 | 5.14 | 84.14 | 53.32 | 32.52 | 51.30 | Luminal-binding protein 4 [*N. attenuata*] |
| Niben101Scf06394g00009 | 7.98 | 37.22 | 22.93 | 12.84 | 7.89 | 144.51 | 33.78 | 29.95 | 51.16 | 2-hydroxyacyl-CoA lyase [*N. attenuata*] |
| Niben101Scf06779g01002 | 3.52 | 5.55 | 11.40 | 3.64 | 3.81 | 36.54 | 35.96 | 28.46 | 51.04 | Inorganic phosphate transporter 1-4-like [*N. sylvestris*] |
| Niben101Scf00773g08003 | 1.97 | 5.80 | 3.74 | 1.79 | 2.29 | 47.79 | 51.32 | 55.86 | 50.33 | Wound-induced protein 1-like, partial [*N. sylvestris*] |
| Niben101Scf01789g03003 | 3.67 | 15.84 | 20.39 | 3.90 | 5.93 | 107.06 | 52.29 | 16.89 | 48.32 | Chitinase-3-like protein 1 [*N. sylvestris*] |
| Niben101Scf13180g01003 | 1.43 | 14.03 | 2.66 | 1.91 | 2.24 | 16.59 | 34.19 | 45.97 | 46.94 | 3-hydroxy-3-methylglutaryl-coenzyme A reductase NbHMGR1a [*N. benthamiana*] |
| Niben101Scf07451g00013 | 2.07 | 10.68 | 4.00 | 2.70 | 3.30 | 22.48 | 41.61 | 42.19 | 45.40 | Calcium-binding allergen Ole e 8-like [*N. tabacum*] |
| Niben101Scf03413g01011 | 11.91 | 35.44 | 25.78 | 16.18 | 11.32 | 135.13 | 74.37 | 58.95 | 44.34 | Mevalonate-5-pyrophosphate decarboxylase NbMVD1a [*N. benthamiana*] |
| Niben101Scf06009g00025 | 3.16 | 7.61 | 4.52 | 4.45 | 2.95 | 38.90 | 27.04 | 51.71 | 43.58 | SNAP25 homologous protein SNAP33 [*N. sylvestris*] |
| Niben101Scf01432g06023 | 3.81 | 8.60 | 4.55 | 3.52 | 3.08 | 29.46 | 88.79 | 82.72 | 43.45 | Caffeoyl-CoA O-methyltransferase 3 [*N. tabacum*] |
| Niben101Scf09030g01004 | 2.92 | 12.33 | 7.89 | 5.16 | 5.39 | 45.88 | 56.24 | 61.17 | 41.95 | Serine/arginine repetitive matrix protein 2-like [*N. sylvestris*] |
| Niben101Scf03710g10004 | 7.50 | 26.90 | 27.09 | 18.76 | 16.51 | 107.76 | 80.15 | 48.87 | 41.45 | Amino acid permease 3-like [*N. sylvestris*] |
| Niben101Scf06583g03009 | 1.24 | 28.38 | 5.84 | 4.09 | 1.65 | 162.75 | 59.45 | 80.33 | 40.47 | PDR-type ACB transporter NbPDR1b [*N. benthamiana*] |
| Niben101Scf00372g05013 | 5.08 | 28.84 | 43.72 | 14.18 | 2.58 | 155.11 | 189.61 | 98.97 | 39.84 | Hypothetical protein A4A49_08095 [*N. attenuata*] |
| Niben101Scf07586g00004 | 0.84 | 8.13 | 3.20 | 2.35 | 1.77 | 27.18 | 31.39 | 40.01 | 39.27 | Calcium-binding protein CAST-like [*N. sylvestris*] |
| Niben101Scf25768g00010 | 4.68 | 8.42 | 8.50 | 6.68 | 3.08 | 30.86 | 17.46 | 34.61 | 39.09 | Receptor-like protein kinase FERONIA [*N. tabacum*] |

*Top 50 genes highly expressed in INF1-treated leaves 24 after the treatment.

**Supplementary Table 8.** Predicted *Nicotiana benthamiana* genes for enzymes specifically involved in the production of capsidiol.

| Gene Name | Gene ID | Cluster ID[1] |
|---|---|---|
| **5-*epi*-Aristolochene synthase (EAS)** | | |
| *NbEAS1* | Niben101Scf07725g01004 | **14** |
| *NbEAS2* | Niben101Scf07725g00004 | **14** |
| *NbEAS3* | Niben101Scf03993g06006 | **14** |
| *NbEAS4* | Niben101Scf03993g05005 | **14** |
| *NbEAS5* | Niben101Scf06245g01017 | **14** |
| *NbEAS6* | Niben101Scf00700g00005 | **14** |
| *NbEAS7* | Niben101Scf00712g02011 | **14** |
| *NbEAS8* | Niben101Scf04362g07009 | **14** |
| *NbEAS9* | Niben101Scf01683g03005 | **14** |
| *NbEAS10* | Niben101Scf03400g02010 | **14** |
| **5-*epi*-Aristolochene dihydroxylase (EAH)** | | |
| *NbEAH1* | Niben101Scf00072g06001 | **14** |
| *NbEAH2* | Niben101Scf04362g07011 | **14** |
| *NbEAH3* | Niben101Scf00072g05003 | **14** |
| *NbEAH4* | Niben101Scf00072g05004 | **14** |
| *NbEAH5* | Niben101Scf00994g00001 | **14** |
| *NbEAH6* | Niben101Scf04869g00002 | n/a |

[1] Cluster numbers for INF1-induced genes are shown in bold.
n/a, not assigned.

**Supplementary Table 9.** Predicted genes for enzymes in mevalonate pathway and farnesylpyrophosphate synthase in *Nicotiana benthamiana.*

| Gene Name | Gene ID | **Cluster ID**[1] |
|---|---|---|
| **Acetyl-CoA thiolase (ACAT)** | | |
| *NbACAT1a*[2] | Niben101Scf04727g03006 | 3 |
| *NbACAT1b* | Niben101Scf01100g01006 | **16** |
| *NbACAT2a* | Niben101Scf06423g02006 | 8 |
| *NbACAT2b* | Niben101Scf01974g00019 | 7 |
| *NbACAT3* | Niben101Scf05433g00004 | 9 |
| **3-Hydroxy-3-methylglutaryl-CoA synthase (HMGS)** | | |
| *NbHMGS1a*[2] | Niben101Scf01111g01003 | **4** |
| *NbHMGS1b* | Niben101Scf01729g01015 | **16** |
| *NbHMGS2a* | Niben101Scf03321g01012 | 9 |
| *NbHMGS2b* | Niben101Scf02361g01002 | 2 |
| *NbHMGS3* | Niben101Scf10595g01007 | 13 |
| **3-Hydroxy-3-methylglutaryl-CoA reductase (HMGR)** | | |
| *NbHMGR1a* | Niben101Scf13180g01003 | **16** |
| *NbHMGR1b* | Niben101Scf09686g00013 | **2** |
| *NbHMGR2*[2] | Niben101Scf02203g05002 | **14** |
| *NbHMGR3a* | Niben101Scf09883g01009 | 13 |
| *NbHMGR3b* | Niben101Scf00163g01009 | 1 |
| **Mevalonate-5-kinase (MVK)** | | |
| *NbMVK1a* | Niben101Scf25893g00005 | **2** |
| *NbMVK1b* | Niben101Scf00370g03023 | **4** |
| **Phosphomevalonate kinase (PMVK)** | | |
| *NbPMVK1a* | Niben101Scf07030g04004 | **2** |
| *NbPMVK1b* | Niben101Scf09628g00020 | 8 |
| **Mevalonate-5-pyrophosphate decarboxylase (MVD)** | | |
| *NbMVD1a*[2] | Niben101Scf03413g01011 | **16** |
| *NbMVD1b* | Niben101Scf00173g05015 | **16** |
| **Isopentenyl pyrophosphate isomerase (IPPS)** | | |
| *NbIPPI1a* | Niben101Scf17839g02005 | **2** |
| *NbIPPI1b* | Niben101Scf05848g05012 | **4** |
| *NbIPPI2a* | Niben101Scf02499g03007 | **2** |
| *NbIPPI2b* | Niben101Scf01514g04018 | 9 |
| **Farnesylpyrophosphate synthase (FPPS)** | | |
| *NbFPPS1a*[2] | Niben101Scf04739g01006 | **16** |
| *NbFPPS1b* | Niben101Scf00414g07005 | **4** |
| *NbFPPS2a* | Niben101Scf04847g02011 | 7 |
| *NbFPPS2b* | Niben101Scf04444g09012 | **4** |

[1] Cluster numbers for INF1-induced genes are shown in bold.

[2] Identified as an essential gene for the resistance to *P. infestans* in Shibata et al., 2016.

**Supplementary Table 10.** Predicted genes for enzymes involved in methionine cycle and ethylene production in *Nicotiana benthamiana* .

| Gene Name | Gene ID | Cluster ID[1] |
|---|---|---|
| **Cystathionine gamma-synthase** | | |
| *NbCGS1a* [2] | Niben101Scf10866g01001 | **2** |
| *NbCGS1b* | Niben101Scf00136g03005 | 9 |
| *NbCGS2a* | Niben101Scf04133g01027 | 13 |
| *NbCGS2b* | Niben101Scf06457g00005 | 13 |
| **Cystathionine beta-lyase** | | |
| *NbCBL1a* | Niben101Scf08333g07030 | 9 |
| *NbCBL1b* | Niben101Scf03006g06008 | 9 |
| *NbCBL2* [3] | Niben101Scf05535g03002 | 7 |
| **Methionine synthase** | | |
| *NbMS1a* | Niben101Scf09725g00022 | 1 |
| *NbMS1b* | Niben101Scf01812g02025 | 1 |
| *NbMS2* | Niben101Scf07438g04014 | 13 |
| *NbMS3* | Niben101Scf03634g04009 | 13 |
| *NbMS4a* | Niben101Scf00054g01013 | 13 |
| *NbMS4b* | Niben101Scf12280g00002 | 13 |
| *NbMS5a* | Niben101Scf09813g00005 | 8 |
| *NbMS5b* | Niben101Scf05629g01024 | 8 |
| *NbMS6* [3] | Niben101Scf10678g00010 | 8 |
| **S-adenosylmethionine (SAM) synthetase** | | |
| *NbSAMS1a* [2, 4] | Niben101Scf03535g01001 | 1 |
| *NbSAMS1b* | Niben101Scf04643g02010 | 11 |
| *NbSAMS2a* [5] | Niben101Scf11751g01007 | 3 |
| *NbSAMS2b* | Niben101Scf02502g04001 | 3 |
| *NbSAMS3a* | Niben101Scf01236g02016 | 13 |
| *NbSAMS3b* | Niben101Scf01820g00025 | 13 |
| *NbSAMS4a* | Niben101Scf01861g00002 | **2** |
| *NbSAMS4b* | Niben101Scf00402g04010 | 9 |
| *NbSAMS5* | Niben101Scf00285g00001 | 13 |
| *NbSAMS6* | Niben101Scf01334g06003 | 7 |
| *NbSAMS7a* | Niben101Scf15054g01008 | 17 |
| *NbSAMS7b* | Niben101Scf04473g14015 | n. a. |
| *NbSAMS8* | Niben101Scf01671g03004 | 12 |
| *NbSAMS9* [3] | Niben101Ctg06829g00002 | 13 |
| *NbSAMS10* [3] | Niben101Scf00402g04005 | 9 |
| **S-adenosylhomocycteine (SAH) hydrolase** | | |
| *NbSAHH1a* [2, 6] | Niben101Scf10608g02009 | 1 |
| *NbSAHH1b* | Niben101Scf09136g00004 | 1 |
| *NbSAHH2a* | Niben101Scf01171g03016 | 7 |
| *NbSAHH2b* | Niben101Scf01592g01001 | 13 |
| *NbSAHH3* | Niben101Scf06737g00025 | 5 |
| *NbSAHH* [3] | Niben101Scf02853g05019<br>Niben101Scf02853g05018 | 5 |
| **Aminocyclopropane carboxylate (ACC) synthase** | | |
| *NbACS1* | Niben101Scf09512g03008 | **14** |
| *NbACS2a* | Niben101Scf06180g00015 | **2** |

| | | |
|---|---|---|
| *NbACS2b* | Niben101Scf02334g00004 | 11 |
| *NbACS3* | Niben101Scf00254g00004 | 8 |
| *NbACS4* | Niben101Scf03226g01003 | **2** |
| *NbACS5a* | Niben101Scf05348g01023 | 19 |
| *NbACS5b* | Niben101Scf05977g01005 | 19 |
| *NbACS6a* | Niben101Scf03907g01018 | 13 |
| *NbACS6b* | Niben101Scf00485g02022 | 8 |
| *NbACS7a* | Niben101Scf00388g07005 | 20 |
| *NbACS7b* | Niben101Scf15817g01015 | 20 |
| *NbACS8a* | Niben101Scf02740g15001 | n. a. |
| *NbACS8b* | Niben101Scf11516g00011 | n. a. |
| *NbACS9a* | Niben101Scf04773g00021 | n. a. |
| *NbACS9b* | Niben101Scf02081g03012 | n. a. |
| *NbACS10a* | Niben101Scf02053g00003 | n. a. |
| *NbACS10b* | Niben101Ctg13861g00002 | n. a. |
| *NbACS11a* | Niben101Scf05449g01004 | n. a. |
| *NbACS11b* | Niben101Scf00108g05008 | n. a. |
| *NbACS12a* | Niben101Scf00926g03006 | n. a. |
| *NbACS12b* | Niben101Scf08472g00003 | n. a. |
| *NbACS13a* | Niben101Scf03238g02003 | n. a. |
| *NbACS13b* | Niben101Scf02816g05004 | n. a. |
| *NbACS14* | Niben101Scf39461g00001 | n. a. |
| **Aminocyclopropane carboxylate (ACC) oxydase** | | |
| *NbACO1a* [2] | Niben101Scf02543g01008 | **10** |
| *NbACO1b* [2] | Niben101Scf08039g01005 | **10** |
| *NbACO2a* [2] | Niben101Scf09590g03007 (LC008355)[7] | 3 |
| *NbACO2b* | Niben101Scf02433g06003 (LC659235)[7] | 9 |
| *NbACO3a* | Niben101Scf09217g00024 | 19 |
| *NbACO3b* | Niben101Scf13622g01020 | 9 |
| *NbACO4* | Niben101Scf02918g00003 | **14** |
| *NbACO5a* | Niben101Scf19336g00004 | n. a. |
| *NbACO5b* | Niben101Scf00313g01001 | n. a. |
| *NbACO6* | Niben101Scf00313g01004 | n. a. |
| *NbACO7a* | Niben101Scf02597g08002 | 19 |
| *NbACO7b* | Niben101Scf00430g00002 | n. a. |
| *NbACO8a* | Niben101Scf14041g00001 | 1 |
| *NbACO8b* | Niben101Scf03710g07016 | 19 |
| *NbACO9a* | Niben101Scf09834g00001 | 8 |
| *NbACO9b* | Niben101Scf15670g00014 | **4** |

[1] Cluster numbers for INF1-induced genes are shown in bold.

[2] Genes essential for the resistance to *P. infestans* (Shibata et al. 2016)

[3] Probable pseudogene or non-coding sequence.

[4] Most homologous to *NbSAM1* in Ismayil et al. (2018).

[5] Most homologous to *NbSAM2* and *NbSAM3* in Ismayil et al. (2018).

[6] Reported as *NbSAHH* in Carmen Cañizares et al. (2013).

[7] The sequences with the Niben101 ID numbers are partial sequences. Full-length cDNA sequences were constructed by *de novo* assembly of RNAseq data and registered under accession nos. shown in parentheses.

n/a, not assigned.

**Supplementary Table 11.** Gene list for predicted AP2/RAV family transcription factors in *Nicotiana benthamiana* .

| Family | Gene ID | Gene Name | Arabidopsis homologues in same branch of phylogenetic tree | | Cluster |
|---|---|---|---|---|---|
| AP2 | Niben101Scf07223g00002 | *NbAP2-1* | | | 12 |
| | Niben101Scf07050g00010 | *NbAP2-2a* | | | 13 |
| | Niben101Scf00496g02009 | *NbAP2-2b* | | | 8 |
| | Niben101Scf07944g03004 | *NbAP2-3a* | AP2 | At4g36920.1 | 13 |
| | Niben101Scf01692g01003 | *NbAP2-3b* | TOE3 | At5g67180.1 | 8 |
| | Niben101Scf01432g01005 | *NbAP2-4a* | | | 7 |
| | Niben101Scf07444g03006 | *NbAP2-4b* | | | 13 |
| | Niben101Scf00280g01007 | *NbAP2-5a* | | | 13 |
| | Niben101Scf25996g00006 | *NbAP2-5b* | | | 13 |
| | Niben101Scf01035g04005 | *NbAP2-6a* | | | 1 |
| | Niben101Scf02702g00005 | *NbAP2-6b* | TOE1/RAP2.7 | At2g28550.3 | 8 |
| | Niben101Scf11756g01019 | *NbAP2-7a* | TOE2 | At5g60120.2 | 7 |
| | Niben101Scf03883g05010 | *NbAP2-7b* | SNZ | At2g39250.1 | 8 |
| | Niben101Scf02369g04004 | *NbAP2-8a* | SMZ | At3g54990.1 | 11 |
| | Niben101Scf00482g08004 | *NbAP2-8b* | | | 18 |
| | Niben101Scf00396g00013 | *NbAP2-9* | | | 19 |
| | Niben101Scf07196g01001 | *NbAP2-10* | PLT1 | At3g20840.1 | n. a. |
| | Niben101Scf07196g01008 | *NbAP2-11a* | PLT2 | At1g51190.1 | 9 |
| | Niben101Scf07196g01007 | *NbAP2-11b* | | | 9 |
| | Niben101Scf04386g00002 | *NbAP2-12* | | | n. a. |
| | Niben101Scf11138g01008 | *NbAP2-13a* | - | At2g41710.3 | 8 |
| | Niben101Scf02156g03004 | *NbAP2-13b* | | | 9 |
| | Niben101Scf07030g00017 | *NbAP2-14a* | | | 13 |
| | Niben101Scf15227g00026 | *NbAP2-14b* | | | 1 |
| | Niben101Scf03437g02045 | *NbAP2-15a* | WRI3 | At1g16060.1 | 13 |
| | Niben101Scf04218g00013 | *NbAP2-15b* | WRI4 | At1g79700.2 | 9 |
| | Niben101Scf00546g00002 | *NbAP2-16* | | | 7 |
| | Niben101Scf05109g04005 | *NbAP2-17a* | | | n. a. |
| | Niben101Scf08876g00008 | *NbAP2-17b* | | | n. a. |
| | Niben101Scf02891g04012 | *NbAP2-18a* | WRI1 | At3g54320.1 | 7 |
| | Niben101Scf13694g00020 | *NbAP2-18b* | | | 8 |
| | Niben101Scf01774g10021 | *NbAP2-19a* | | | 9 |
| | Niben101Scf03419g03008 | *NbAP2-19b* | ANT | At4g37750.1 | 8 |
| | Niben101Scf00388g03006 | *NbAP2-20a* | | | 7 |
| | Niben101Scf07169g02008 | *NbAP2-20b* | | | 9 |
| | Niben101Scf11515g00016 | *NbAP2-21a* | | | 12 |
| | Niben101Scf07511g02006 | *NbAP2-21b* | AIL6 | At5g10510.3 | n. a. |
| | Niben101Scf04574g07001 | *NbAP2-22a* | AIL7 | At5g65510.1 | 1 |
| | Niben101Scf00173g09005 | *NbAP2-22b* | | | n. a. |
| | Niben101Scf01438g04007 | *NbAP2-23a* | AIL5 | At5g57390.1 | 13 |
| | Niben101Scf02540g00013 | *NbAP2-23b* | | | 7 |
| | Niben101Scf04691g00002 | *NbAP2-24a* | BBM | At5g17430.1 | n. a. |
| | Niben101Scf06407g01001 | *NbAP2-24b* | | | n. a. |
| | Niben101Scf05123g00001 | *NbAP2-25a* | AIL1 | At1g72570.1 | 11 |
| | Niben101Scf02749g05039 | *NbAP2-25b* | | | 1 |
| Soloist I | Niben101Scf03479g03001 | *NbAP2-S-1a* | APD1 | At4g13040.3 | 8 |
| | Niben101Scf00113g03002 | *NbAP2-S-1b* | | | **2** |
| RAV | Niben101Scf10767g03010 | *NbRAV-1* | | | 8 |
| | Niben101Scf06059g01001 | *NbRAV-2a* | RAV2/RAP2.8 | At1g68840.1 | 11 |
| | Niben101Scf16806g00003 | *NbRAV-2b* | EDF3 | At3g25730.1 | n. a. |
| | Niben101Scf03150g06002 | *NbRAV-3a* | TEM1 | At1g25560.1 | **16** |
| | Niben101Scf00872g03005 | *NbRAV-3b* | RAV1/EDF4 | At1g13260.1 | 11 |
| | Niben101Scf07498g01011 | *NbRAV-4a* | - | At1g50680.1 | n. a. |
| | Niben101Scf03488g06004 | *NbRAV-4b* | - | At1g51120.1 | n. a. |

[1] Cluster numbers for INF1-induced genes are shown in bold.
n. a., not assigned.

**Supplementary Table 12.** Gene list for predicted ERF family transcription factors in *Nicotiana benthamiana*.

| Sub family | | Gene ID | Gene Name | Arabidopsis homologues in same branch of phylogenetic tree | | Cluster[1] |
|---|---|---|---|---|---|---|
| ERF-I, -IIa | (DREB A-6) | Niben101Scf01177g01001 | *NbERF-I-1a* | | | 18 |
| | | Niben101Scf09459g02008 | *NbERF-I-1b* | ERF15 | At4g31060.1 | 6 |
| | | Niben101Scf13695g01014 | *NbERF-I-2a* | ERF61 | At1g64380.1 | 13 |
| | | Niben101Scf18347g00023 | *NbERF-I-2b* | ERF62 | At4g13620.1 | 13 |
| | | Niben101Scf03850g01005 | *NbERF-I-3a* | | | 11 |
| | | Niben101Scf02972g00008 | *NbERF-I-3b* | | | 3 |
| | | Niben101Scf06119g02002 | *NbERF-I-4a* | TG (ERF55) | At1g36060.1 | 8 |
| | | Niben101Scf01980g04003 | *NbERF-I-4b* | ERF56 | At2g22200.1 | 11 |
| | | Niben101Scf02219g02002 | *NbERF-I-5a* | WIND4 (ERF57) | At5g65130.1 | 7 |
| | | Niben101Scf10283g01004 | *NbERF-I-5b* | RAP2.4D/WIND2 (ERF58) | At1g22190.1 | 13 |
| | | Niben101Scf03510g06011 | *NbERF-I-6a* | RAP2.4/WIND1 (ERF59) | At1g78080.1 | 8 |
| | | Niben101Scf00846g02001 | NbERF-I-6b | ERF60 | At4g39780.1 | 8 |
| | | Niben101Scf05440g02002 | *NbERF-I-7a* | | | 9 |
| | | Niben101Scf34675g00005 | *NbERF-I-7b* | ERF53 | At2g20880.1 | n. a. |
| | | Niben101Scf06117g01012 | *NbERF-I-8a* | ERF54 | At4g28140.1 | 13 |
| | | Niben101Scf06120g00005 | *NbERF-I-8b* | | | 3 |
| | | Niben101Scf10763g00021 | *NbERF-I-9* | | | n.a. |
| | (DREB A-5) | Niben101Scf01692g03007 | *NbERF-II-1a* | RAP2.1 (ERF6) | At1g46768.1 | 8 |
| | | Niben101Scf04826g01005 | *NbERF-II-1b* | RAP2.9 (ERF7) | At4g06746.1 | 13 |
| | | Niben101Scf07066g03005 | *NbERF-II-2a* | DEAR3 (ERF8) | At2g23340.1 | 9 |
| | | Niben101Scf05135g06001 | *NbERF-II-2b* | RAP2.10 (ERF9) | At4g36900.1 | 3 |
| | | Niben101Scf06444g02001 | *NbERF-II-3a* | DEAR2 (ERF10) | At5g67190.1 | 7 |
| | | Niben101Scf03314g00004 | *NbERF-II-3b* | CEJ1 (ERF11) | At3g50260.1 | 1 |
| | | Niben101Scf03223g00002 | *NbERF-II-4* | | | 19 |
| ERF-IIb, -IIc | (DREB A-5) | Niben101Scf15963g01004 | *NbERF-II-5a* | ERF19 | At1g22810.1 | n. a. |
| | | Niben101Scf06473g01014 | *NbERF-II-5b* | ERF20 | At1g71520.1 | n. a. |
| | | Niben101Scf06596g03021 | *NbERF-II-6a* | DREB26 (ERF12) | At1g21910.1 | 1 |
| | | Niben101Scf09087g00009 | *NbERF-II-6b* | ERF13 | At1g77640.1 | 12 |
| | | | | ERF14 | At1g44830.1 | 1 |
| | | Niben101Scf00298g00001 | *NbERF-II-7a* | | | n. a. |
| | | Niben101Scf02437g08005 | *NbERF-II-7b* | ERF16 | At5g21960.1 | 17 |
| | | Niben101Scf11694g01005 | *NbERF-II-8* | ERF17 | At1g19210.1 | n. a. |
| | | Niben101Scf05773g00002 | *NbERF-II-9* | ORA47 (ERF18) | At1g74930.1 | n. a. |
| | | Niben101Scf02744g00004 | *NbERF-II-10a* | | | n. a. |
| | | Niben101Scf08651g04017 | *NbERF-II-10b* | | | n. a. |
| ERF-III | (DREB A-4) | Niben101Scf01611g11002 | *NbERF-III-1a* | | | 13 |
| | | Niben101Scf05156g01011 | *NbERF-III-1b* | ERF43 | At4g32800.1 | 13 |
| | | Niben101Scf08776g00003 | *NbERF-III-2* | ESE2 (ERF42) | At2g25820.1 | n. a. |
| | | Niben101Scf09363g00010 | *NbERF-III-3a* | ERF37 | At1g77200.1 | n. a. |
| | | Niben101Scf01374g05001 | *NbERF-III-3b* | TINY (ERF40) | At5g25810.1 | **2** |
| | | Niben101Scf01539g00001 | *NbERF-III-4* | TINY2 (ERF41) | At5g11590.1 | 11 |
| | | Niben101Scf06969g00003 | *NbERF-III-5* | ERF23 | At1g01250.1 | **2** |
| | | Niben101Scf04337g01005 | *NbERF-III-6* | ERF36 | At3g16280.1 | 13 |
| | | Niben101Scf00603g02001 | *NbERF-III-7* | | | 13 |
| | | Niben101Scf05855g06014 | *NbERF-III-8a* | | | 11 |
| | | Niben101Scf00315g02001 | *NbERF-III-8b* | | | n. a. |
| | | Niben101Scf07693g00011 | *NbERF-III-9a* | ERF34 | At2g44940.1 | 6 |
| | | Niben101Scf06436g05001 | *NbERF-III-9b* | ERF35 | At3g60490.1 | 3 |
| | | Niben101Scf15704g00006 | *NbERF-III-10* | ERF38 | At2g35700.1 | 13 |
| | | Niben101Scf06157g01005 | *NbERF-III-11a* | ERF39 | At4g16750.1 | 11 |
| | | Niben101Scf03772g00001 | *NbERF-III-11b* | | | n. a. |
| | (DREB A-1) | Niben101Scf03321g01009 | *NbERF-III-12a* | | | 20 |
| | | Niben101Scf03092g00007 | *NbERF-III-12b* | | | n. a. |
| | | Niben101Scf05459g00001 | *NbERF-III-13* | | | n. a. |
| | | Niben101Ctg16385g00003 | *NbERF-III-14* | | At1g12610.1 | n. a. |
| | | Niben101Scf03245g04004 | *NbERF-III-15a* | DDF1 (ERF33) | At1g63030.1 | n. a. |
| | | Niben101Scf03245g04005 | *NbERF-III-15b* | DDF2 (ERF32) | At4g25490.1 | n. a. |
| | | Niben101Scf16007g00010 | *NbERF-III-16a* | CBF1/DREB1B (ERF29) | At4g25470.1 | n. a. |
| | | Niben101Scf16007g00011 | *NbERF-III-16b* | CBF2/DREB1C (ERF30) | At4g25480.1 | n. a. |
| | | Niben101Scf00282g02006 | *NbERF-III-17* | CBF3/DREB1A (ERF31) | At5g51990.1 | 3 |
| | | Niben101Scf00085g00008 | *NbERF-III-18a* | | | n. a. |
| | | Niben101Scf06081g02026 | *NbERF-III-18b* | | | n. a. |
| | (DREB A-4) | Niben101Scf04217g07002 | *NbERF-III-19* | | | n. a. |
| | | Niben101Scf08840g00003 | *NbERF-III-20a* | | | 6 |
| | | Niben101Scf08840g00008 | *NbERF-III-20b* | | | **16** |
| | | Niben101Scf16506g00004 | *NbERF-III-21* | HRD (ERF24) | At2g36450.1 | n. a. |
| | | Niben101Scf01111g02001 | *NbERF-III-22* | ERF25 | At5g52020.1 | n. a. |
| | | Niben101Scf03245g06003 | *NbERF-III-23a* | ERF26 | At1g63040.1 | 11 |
| | | Niben101Scf02577g03014 | *NbERF-III-23b* | ERF27 | At1g12630.1 | n. a. |
| | | Niben101Scf00784g01003 | *NbERF-III-24* | | | n. a. |

| | | | | | | |
|---|---|---|---|---|---|---|
| | | Niben101Scf03580g01003 | *NbERF-III-25a* | | | n. a. |
| | | Niben101Scf11954g01002 | *NbERF-III-25b* | | | n. a. |
| | | Niben101Scf06848g08006 | *NbERF-III-26a* | | | n. a. |
| | | Niben101Scf02182g19004 | *NbERF-III-26b* | FUF1 (ERF21) | At1g71450.1 | n. a. |
| | | Niben101Scf06848g09002 | *NbERF-III-27a* | ERF22 | At1g33760.1 | n. a. |
| | | Niben101Scf02182g21007 | *NbERF-III-27b* | | | n. a. |
| ERF-IV | (DREB-A2) | Niben101Scf00735g02005 | *NbERF-IV-1a* | | | 1 |
| | | Niben101Scf05993g01013 | *NbERF-IV-1b* | DREB2F (ERF51) | At3g57600.1 | 12 |
| | | Niben101Scf04505g02007 | *NbERF-IV-2* | | | 15 |
| | | Niben101Scf00177g02016 | *NbERF-IV-3* | DREB2G (ERF50) | At5g18450.1 | 11 |
| | | Niben101Scf01701g03007 | *NbERF-IV-4a* | | | n. a. |
| | | Niben101Scf08033g01007 | *NbERF-IV-4b* | DREB2D (ERF49) | At1g75490.1 | n. a. |
| | | Niben101Scf07226g08005 | *NbERF-IV-5* | | | 11 |
| | | Niben101Scf10169g00004 | *NbERF-IV-6a* | | | 8 |
| | | Niben101Scf01237g06003 | *NbERF-IV-6b* | DREB2B (ERF44) | At3g11020.1 | **10** |
| | | Niben101Scf06378g04004 | *NbERF-IV-7a* | DREB2A (ERF45) | At5g05410.1 | 8 |
| | | Niben101Scf08060g00005 | *NbERF-IV-7b* | DREB19 (ERF46) | At2g38340.1 | 8 |
| | | Niben101Scf03202g13015 | *NbERF-IV-8a* | DREB2H (ERF47) | At2g40350.1 | n. a. |
| | | Niben101Scf01297g01001 | *NbERF-IV-8b* | DREB2C (ERF48) | At2g40340.1 | n. a. |
| | | Niben101Scf05849g00002 | *NbERF-IV-9* | | | **2** |
| ERF-V | | Niben101Scf04131g01028 | *NbERF-V-1a* | | | 17 |
| | | Niben101Scf04131g01007 | *NbERF-V-1b* | | | n. a. |
| | | Niben101Scf05800g00005 | *NbERF-V-2* | | | n. a. |
| | | Niben101Scf07926g07004 | *NbERF-V-3a* | ESE3 (ERF3) | At5g25190.1 | 7 |
| | | Niben101Scf04437g00010 | *NbERF-V-3b* | | | 17 |
| | | Niben101Scf05618g01002 | *NbERF-V-4* | | | n. a. |
| | | Niben101Scf01142g04009 | *NbERF-V-5a* | | | 8 |
| | | Niben101Scf06190g06004 | *NbERF-V-5b* | | | 3 |
| | | Niben101Scf02397g00007 | *NbERF-V-6a* | | | n. a. |
| | | Niben101Scf09659g03018 | *NbERF-V-6b* | | | n. a. |
| | | Niben101Scf01669g04022 | *NbERF-V-7* | | | n. a. |
| | | Niben101Scf02475g08017 | *NbERF-V-8a* | | | n. a. |
| | | Niben101Scf03732g03003 | *NbERF-V-8b* | | | n. a. |
| | | Niben101Scf04763g00013 | *NbERF-V-9a* | | | 1 |
| | | Niben101Scf04763g00014 | *NbERF-V-9b* | RAP2.11 (ERF2) | At5g19790.1 | 13 |
| | | Niben101Scf06026g00002 | *NbERF-V-10* | | | **2** |
| | | Niben101Scf17051g00007 | *NbERF-V-11* | | | n. a. |
| | | Niben101Scf15962g00025 | *NbERF-V-12a* | | | n. a. |
| | | Niben101Scf06348g00034 | *NbERF-V-12b* | | | n. a. |
| | | Niben101Scf11303g01019 | *NbERF-V-13a* | | | 8 |
| | | Niben101Scf11341g00009 | *NbERF-V-13b* | | | **16** |
| | | Niben101Scf09882g01001 | *NbERF-V-14a* | | | 5 |
| | | Niben101Ctg12291g00001 | *NbERF-V-14b* | | | n. a. |
| | | Niben101Scf03930g01015 | *NbERF-V-15* | SHN1/WIN1 (ERF1) | At1g15360 | n. a. |
| | | Niben101Scf07913g00005 | *NbERF-V-16* | SHN2 (ERF4) | At5g11190.1 | 15 |
| | | Niben101Scf12609g01006 | *NbERF-V-17a* | SHN3 (ERF5) | At5g25390.2 | 11 |
| | | Niben101Scf03832g02022 | *NbERF-V-17b* | | | 11 |
| ERF-VI | | Niben101Ctg15705g00001 | *NbERF-VI-1a* | | | 13 |
| | | Niben101Ctg15705g00004 | *NbERF-VI-1b* | | | 13 |
| | | Niben101Scf00725g02012 | *NbERF-VI-2* | | | 13 |
| | | Niben101Scf01075g01003 | *NbERF-VI-3* | | | 19 |
| | | Niben101Scf00753g02005 | *NbERF-VI-4* | CRF9 (ERF117) | At1g49120.1 | 9 |
| | | Niben101Scf04409g00010 | *NbERF-VI-5a* | | | 11 |
| | | Niben101Scf03817g12030 | *NbERF-VI-5b* | | | 7 |
| | | Niben101Scf08575g01003 | *NbERF-VI-6a* | | | 13 |
| | | Niben101Scf02222g00019 | *NbERF-VI-6b* | | | 17 |
| | | Niben101Scf00278g05001 | *NbERF-VI-7a* | | | 11 |
| | | Niben101Scf00530g01001 | *NbERF-VI-7b* | | | 8 |
| | | Niben101Scf08278g00007 | *NbERF-VI-8* | | | 7 |
| | | Niben101Scf02726g03006 | *NbERF-VI-9a* | CRF1 (ERF63) | At4g11140.1 | 1 |
| | | Niben101Scf01543g03001 | *NbERF-VI-9b* | CRF2/TMO3 (ERF64) | At4g23750.1 | 13 |
| | | Niben101Scf00503g07005 | *NbERF-VI-10a* | CRF3 (ERF65) | At5g53290.1 | **4** |
| | | Niben101Scf03773g00009 | *NbERF-VI-10b* | CRF4 (ERF66) | At4g27950.1 | **4** |
| | | Niben101Ctg13994g00001 | *NbERF-VI-11a* | CRF6 (ERF67) | At3g61630.1 | 19 |
| | | Niben101Scf01817g01001 | *NbERF-VI-11b* | CRF5 (ERF68) | At2g46310.1 | 19 |
| | | Niben101Scf03985g01005 | *NbERF-VI-12* | CRF7 (ERF69) | At1g22985.1 | 13 |
| | | Niben101Scf01001g06011 | *NbERF-VI-13a* | CRF8 (ERF70) | At1g71130.1 | 9 |
| | | Niben101Scf09004g00001 | *NbERF-VI-13b* | | | 9 |
| | | Niben101Scf09004g00003 | *NbERF-VI-14* | | | n. a. |
| | | Niben101Scf01518g08004 | *NbERF-VI-15a* | | | 8 |
| | | Niben101Scf00577g11004 | *NbERF-VI-15b* | | | 9 |
| | | Niben101Scf00262g04006 | *NbERF-VI-16a* | | | 3 |
| | | Niben101Scf02316g03001 | *NbERF-VI-16b* | CRF10 (ERF118) | At1g68550.1 | 3 |
| | | Niben101Scf04860g02011 | *NbERF-VI-17* | CRF11 (ERF119) | At3g25890 | n. a. |
| | | Niben101Scf01695g01005 | *NbERF-VI-18* | CRF12 (ERF116) | At1g25470.1 | 8 |

| | | | | | |
|---|---|---|---|---|---|
| | Niben101Scf12874g00001 | *NbERF-VI-19* | | | n. a. |
| | Niben101Scf08196g00012 | *NbERF-VI-20a* | | | **4** |
| | Niben101Scf01085g03003 | *NbERF-VI-20b* | | | **4** |
| ERF-VII | Niben101Scf06913g00001 | *NbERF-VII-1a* | | | 3 |
| | Niben101Scf01956g05006 | *NbERF-VII-1b* | | | **2** |
| | Niben101Scf01430g00007 | *NbERF173* [2]*/NbERF-VII-2* | ERF71 | At2g47520.1 | 3 |
| | Niben101Scf05827g07015 | *NbERF-VII-3a* | AtEBP/RAP2.3 (ERF72) | At3g16770.1 | 9 |
| | Niben101Scf11139g00001 | *NbERF-VII-3b* | ERF73 | At1g72360.1 | 9 |
| | Niben101Scf19733g00001 | *NbERF-VII-4a* | RAP2.12 (ERF74) | At1g53910.1 | 3 |
| | Niben101Scf05123g02008 | *NbERF-VII-4b* | RAP2.2 (ERF75) | At3g14230.1 | 7 |
| | Niben101Scf00870g05003 | *NbERF-VII-5* | | | 7 |
| ERF-VIII | - | - | ERF112 | At2g33710.2 | n. a. |
| | | | ESR1 (ERF89) | At1g12980.1 | n. a. |
| | Niben101Scf01818g07006 | *NbERF-VIII-1a* | | | n. a. |
| | Niben101Scf10785g01002 | *NbERF-VIII-1b* | | | n. a. |
| | Niben101Scf10785g01014 | *NbERF-VIII-2* | | | n. a. |
| | Niben101Scf01818g08019 | *NbERF-VIII-3a* | | | n. a. |
| | Niben101Ctg14204g00003 | *NbERF-VIII-3b* | LEP (ERF85) | At5g13910.1 | n. a. |
| | Niben101Scf12639g00001 | *NbERF-VIII-4a* | PUCHI (ERF86) | At5g18560.1 | n. a. |
| | Niben101Scf01295g05006 | *NbERF-VIII-4b* | ERF87 | At1g28160.1 | n. a. |
| | Niben101Scf00941g00007 | *NbERF-VIII-5a* | ERF88 | At1g12890.1 | n. a. |
| | Niben101Scf19091g00016 | *NbERF-VIII-5b* | DRNL (ERF90) | At1g24590.1 | n. a. |
| | Niben101Scf03045g14001 | *NbERF-VIII-6* | | | 13 |
| | Niben101Scf02720g08001 | *NbERF-VIII-7a* | | | 8 |
| | Niben101Scf08050g00012 | *NbERF-VIII-7b* | | | 3 |
| | Niben101Scf08597g01023 | *NbERF-VIII-8a* | | | 13 |
| | Niben101Scf03657g02001 | *NbERF-VIII-8b* | | | n. a. |
| | Niben101Scf02543g00002 | *NbERF-VIII-9* | | | 8 |
| | Niben101Scf12483g01009 | *NbERF-VIII-10a* | | | 13 |
| | Niben101Scf01767g03004 | *NbERF-VIII-10b* | ERF3 (ERF82) | At1g50640.1 | 13 |
| | Niben101Scf07310g01001 | *NbERF-VIII-11* | ERF7 (ERF83) | At3g20310.1 | 3 |
| | Niben101Scf05692g06003 | *NbERF-VIII-12a* | | | 3 |
| | Niben101Scf02073g01009 | *NbERF-VIII-12b* | | | n. a. |
| | Niben101Scf09236g00001 | *NbERF-VIII-13a* | | | 8 |
| | Niben101Scf01752g05003 | *NbERF-VIII-13b* | | | 3 |
| | Niben101Scf06200g01006 | *NbERF-VIII-14a* | | | n. a. |
| | Niben101Scf19266g01015 | *NbERF-VIII-14b* | | | n. a. |
| | Niben101Scf06200g00001 | *NbERF-VIII-15a* | | | **2** |
| | Niben101Scf19266g00006 | *NbERF-VIII-15b* | | | n. a. |
| | Niben101Scf13779g00003 | *NbERF-VIII-16a* | ERF11 (ERF76) | At1g28370.1 | 13 |
| | Niben101Scf07896g01006 | *NbERF-VIII-16b* | ERF10 (ERF77) | At1g03800.1 | 3 |
| | Niben101Scf09704g02001 | *NbERF-VIII-17a* | ERF4/RAP2.5 (ERF78) | At3g15210.1 | **16** |
| | Niben101Scf04350g04004 | *NbERF-VIII-17b* | ERF8 (ERF79) | At1g53170.1 | 3 |
| | Niben101Scf11424g01001 | *NbERF-VIII-18a* | ERF9 (ERF80) | At5g44210.1 | 3 |
| | Niben101Scf09191g00005 | *NbERF-VIII-18b* | ERF12 (ERF81) | At1g28360.1 | 3 |
| | Niben101Scf01426g00001 | *NbERF-VIII-19a* | | | **4** |
| | Niben101Scf03035g01006 | *NbERF-VIII-19b* | | | **2** |
| | Niben101Scf05480g00003 | *NbERF-VIII-20a* | | | **4** |
| | Niben101Scf01008g00004 | *NbERF-VIII-20b* | | | **4** |
| ERF-IX | Niben101Scf07761g00004 | *NbERF-IX-1a* | | | n. a. |
| | Niben101Scf01956g12014 | *NbERF-IX-1b* | - | - | n. a. |
| | Niben101Scf01177g04007 | *NbERF-IX-2* | | | 13 |
| | Niben101Scf08965g00003 | *NbERF-IX-3a* | | | 3 |
| | Niben101Scf06413g01002 | *NbERF-IX-3b* | | | n. a. |
| | Niben101Scf36372g00002 | *NbERF-IX-4a* | | | 7 |
| | Niben101Scf06413g01001 | *NbERF-IX-4b* | | | n. a. |
| | Niben101Scf01947g01005 | *NbERF-IX-5a* | | | 19 |
| | Niben101Scf13455g01002 | *NbERF-IX-5b* | | | 15 |
| | Niben101Scf01947g01008 | *NbERF-IX-6* | ERF1 (ERF92) | At3g23240.1 | **4** |
| | Niben101Scf15451g00001 | *NbERF-IX-7* | ERF15 (ERF93) | At2g31230.1 | **14** |
| | Niben101Scf08141g01011 | *NbERF-IX-8a* | ORA59 (ERF94) | At1g06160.1 | **10** |
| | Niben101Ctg04506g00002 | *NbERF-IX-8b* | | | **2** |
| | Niben101Scf01795g06002 | *NbERF-IX-9a* | | | n. a. |
| | Niben101Scf06328g00004 | *NbERF-IX-9b* | | | n. a. |
| | Niben101Scf02063g05001 | *NbERF-IX-10a* | | | **4** |
| | Niben101Scf07761g02006 | *NbERF-IX-10b* | | | **4** |
| | Niben101Scf01090g02001 | *NbERF-IX-11a* | | | n. a. |
| | Niben101Scf17204g01001 | *NbERF-IX-11b* | | | **14** |
| | Niben101Scf07105g04007 | *NbERF-IX-12* | ESE1 (ERF95) | At3g23220.1 | n. a. |
| | Niben101Scf07761g00005 | *NbERF-IX-13* | ERF96 | At5g43410.1 | n. a. |
| | Niben101Scf01428g04013 | *NbERF-IX-14a* | ERF14 (ERF97) | At1g04370.1 | n. a. |
| | Niben101Scf01956g10015 | *NbERF-IX-14b* | | | n. a. |
| | Niben101Scf17204g01003 | NbERF-IX-15a | TDR1 (ERF98) | At3g23230.1 | n. a. |
| | Niben101Scf01090g00001 | NbERF-IX-15b | | | n. a. |

| | | | | | | |
|---|---|---|---|---|---|---|
| | | Niben101Scf01212g03005 | *NbERF-IX-16a* | | | **14** |
| | | Niben101Scf01094g03014 | *NbERF-IX-16b* | | | n. a. |
| | | Niben101Ctg16023g00001 | *NbERF-IX-17a* | | | n. a. |
| | | Niben101Scf01956g12018 | *NbERF-IX-17b* | ERF91 | At4g18450.1 | n. a. |
| | | Niben101Scf06413g02012 | *NbERF-IX-18* | | | n. a. |
| | | Niben101Scf01212g03003 | *NbERF-IX-19a* | | | n. a. |
| | | Niben101Scf01094g03016 | *NbERF-IX-19b* | | | n. a. |
| | | Niben101Scf05173g07004 | *NbERF-IX-20* | | | n. a. |
| | | Niben101Scf01159g00002 | *NbERF-IX-21a* | | | n. a. |
| | | Niben101Scf05173g05003 | *NbERF-IX-21b* | | | n. a. |
| | | Niben101Scf05173g07005 | *NbERF-IX-22* | | | n. a. |
| | | Niben101Scf05173g08003 | *NbERF-IX-23* | | | n. a. |
| | | Niben101Scf05173g08002 | *NbERF1/NbERF-IX-24* [3] | | | n. a. |
| | | Niben101Scf01082g07003 | *NbERF-IX-25a* | | | 11 |
| | | Niben101Scf05173g09004 | *NbERF-IX-25b* | | | n. a. |
| | | Niben101Scf01082g04001 | *NbERF-IX-26a* | ERF13 (ERF99) | At2g44840.1 | n. a. |
| | | Niben101Scf05855g00002 | *NbERF-IX-26b* | ERF1A (ERF100) | At4g17500.1 | n. a. |
| | | Niben101Scf06951g00004 | *NbERF-IX-27* | AtERF2 (ERF101) | At5g47220.1 | 11 |
| | | Niben101Scf19057g00005 | *NbERF-IX-28* | | | n. a. |
| | | Niben101Scf00428g10006 | *NbERF-IX-29a* | | | **14** |
| | | Niben101Scf02775g00003 | *NbERF-IX-29b* | | | n. a. |
| | | Niben101Scf02775g00001 | *NbERF-IX-30* | | | 20 |
| | | Niben101Scf00428g09009 | *NbERF-IX-31a* | | | 3 |
| | | Niben101Scf02775g00002 | *NbERF-IX-31b* | | | **16** |
| | | Niben101Scf08546g00001 | *NbERF-IX-32* | | | n. a. |
| | | Niben101Scf00454g04003 | *NbERF-IX-33a* | | | **4** |
| | | Niben101Scf06865g00004 | *NbERF-IX-33b* | | | **16** |
| | | Niben101Scf01177g04031 | *NbERF-IX-34* | | | 3 |
| | | Niben101Scf00454g03001 | *NbERF-IX-35a* | | | **16** |
| | | Niben101Scf12210g07001 | *NbERF-IX-35b* | | | **4** |
| | | Niben101Scf00163g22003 | *NbERF-IX-36* | AtERF5 (ERF102) | At5g47230.1 | n. a. |
| | | Niben101Scf00163g22004 | *NbERF-IX-37* | AtERF6 (ERF103) | At4g17490.1 | **4** |
| | | Niben101Scf08546g05004 | *NbERF-IX-38a* | ERF104 | At5g61600.1 | **16** |
| | | Niben101Scf08546g05002 | *NbERF-IX-38b* | ERF105 | At5g51190.1 | **16** |
| | | Niben101Scf00163g22002 | *NbERF-IX-39a* | | | **16** |
| | | Niben101Scf08546g03002 | *NbERF-IX-39b* | | | **4** |
| | | Niben101Scf23241g00003 | *NbERF-IX-40a* | | | **2** |
| | | Niben101Scf03310g00003 | *NbERF-IX-40b* | | | **4** |
| | | Niben101Scf08546g05005 | *NbERF-IX-41* | | | **4** |
| | | Niben101Scf08249g02004 | *NbERF-IX-42* | ERF106 | At5g07580.1 | 7 |
| | | Niben101Scf00454g02008 | *NbERF-IX-43a* | DEWAX (ERF107) | At5g61590.1 | 1 |
| | | Niben101Scf00454g02007 | *NbERF-IX-43b* | | | 6 |
| ERF-X | | Niben101Scf04634g00008 | *NbERF-X-1a* | | | 6 |
| | | Niben101Ctg15010g00002 | *NbERF-X-1b* | | | 6 |
| | | Niben101Scf01453g11005 | *NbERF-X-2a* | | | 18 |
| | | Niben101Scf01942g08004 | *NbERF-X-2b* | | | 18 |
| | | Niben101Scf12210g08011 | *NbERF-X-3a* | RAP2.6 (ERF108) | At1g43160.1 | **16** |
| | | Niben101Scf02242g03012 | *NbERF-X-3b* | ERF110 | At5g50080.1 | 20 |
| | | Niben101Scf10438g00010 | *NbERF-X-4a* | ABR1 (ERF111) | At5g64750.1 | n. a. |
| | | Niben101Scf10438g00009 | *NbERF-X-4b* | RAP2.6L (ERF113) | At5g13330.1 | 20 |
| | | Niben101Scf11706g00010 | *NbERF-X-5* | ERF114 | At5g61890.1 | 20 |
| | | Niben101Scf06621g02032 | *NbERF-X-6a* | ERF115 | At5g07310.1 | 9 |
| | | Niben101Scf03548g02037 | *NbERF-X-6b* | | | 9 |
| | | Niben101Scf07579g01012 | *NbERF-X-7* | | | **2** |
| | | Niben101Scf14233g00008 | *NbERF-X-8a* | | | **16** |
| | | Niben101Scf00725g06004 | *NbERF-X-8b* | | | n. a. |
| | | Niben101Scf00321g03008 | *NbERF-X-9a* | | | n. a. |
| | | Niben101Scf02483g02007 | *NbERF-X-9b* | | | n. a. |
| | | Niben101Scf06203g05020 | *NbERF-X-10* | RRTF1 (ERF109) | At4g34410.1 | n. a. |
| | | Niben101Scf02408g00021 | *NbERF-X-11a* | ERF120 | At2g20350.1 | n. a. |
| | | Niben101Scf07172g00019 | *NbERF-X-11b* | ERF121 | At5g67010.1 | 18 |
| | | Niben101Scf05140g01007 | *NbERF-X-12a* | ERF122 | At5g67000.1 | n. a. |
| | | Niben101Scf01350g07008 | *NbERF-X-12b* | | | n. a. |
| | | Niben101Scf00839g00006 | *NbERF-X-13* | | | n. a. |
| Soloist II | | Niben101Scf08516g01005 | *NbERF-S-1* | ERF84 | At1g80580.1 | n. a. |
| Soloist III | (DREB-A3) | Niben101Scf05357g00008 | *NbERF-S-2a* | ABI4 (ERF52) | At2g40220.1 | n. a. |
| | | Niben101Scf06984g02002 | *NbERF-S-2b* | | | n. a. |

[1] Cluster numbers for INF1-induced genes are shown in bold.

[2] NbERF173/NbERF-VII-2 was reported as a gene involved in the resistace of *N. benthamiana* against *Phytophthora parasitica* (Yu et al., 2020).

[3] NbERF1/NbERF-IX-24 was reported as a gene involved in the MeJA-induced nicotine biosynthesis (Todd et al., 2010).

n. a., not assigned.
